# Structural insights into DHODH-mediated catalysis and drug resistance

**DOI:** 10.64898/2026.09.23.753858

**Authors:** Ruchika Pokhriyal, Hyuk-Soo Seo, Mio Murakoso, Sonia E. Trojan, Laura Evans, Jillian K. O’Neil, Komal Chauhan, Shakchhi Joshi, Hanna Meyer, Anders Friberg, Judith Günther, Sven Giese, Sven Christian, Xiaoping Yang, Robert E. Lintner, David E. Root, Marcia C. Haigis, Matthew Vander Heiden, Marco Mravic, Sirano Dhe-Paganon, Julie-Aurore Losman

## Abstract

Dihydroorotate dehydrogenase (DHODH), which catalyzes the rate-limiting step in *de novo* pyrimidine biosynthesis, is a validated therapeutic target in cancer, autoimmune disorders and infectious diseases. DHODH is hypothesized to utilize a ‘ping-pong’ catalytic mechanism. However, available crystal structures of DHODH in complex with small molecule inhibitors are irreconcilable with this model, showing simultaneous occupancy of both substrate binding sites. To elucidate the structural basis for DHODH-mediated catalysis, we resolved the structures of two DHODH holoenzymes. These structures capture novel conformational states that show mutually exclusive substrate binding. The holoenzyme structures also suggest that conformational changes in the catalytic loop of DHODH play a critical role in regulating enzyme activity. To map the functional landscape underlying DHODH inhibitor resistance, we performed deep mutational scanning drug-resistance screens with two clinically-relevant structurally distinct DHODH inhibitors, BAY2402234 and brequinar. Resistance variants cluster in the inhibitor binding site and at a previously unappreciated surface pocket on DHODH that allosterically regulates ubiquinone binding. Together, our findings provide structural evidence for the ‘ping pong’ model of catalysis, define the mutational landscape governing inhibitor resistance, and reveal novel structural vulnerabilities in DHODH that can be utilized for future drug development.

## Introduction

*De novo* pyrimidine biosynthesis and pyrimidine salvage are the two mechanisms by which cells can acquire pyrimidines (**Extended Data Fig. 1a**)^1,2^. Rapidly proliferating cells have high pyrimidine requirements and are dependent on *de novo* synthesis, and many human pathogens do not express salvage enzymes, making *de novo* synthesis their only means of acquiring pyrimidines^3,4^. This has prompted extensive efforts to target *de novo* pyrimidine biosynthesis for the treatment of various diseases, including autoimmune disorders, cancer, and microbial infections. Dihydroorotate dehydrogenase (DHODH), the rate-limiting enzyme in *de novo* pyrimidine biosynthesis, is highly amenable to small molecule inhibition, and numerous DHODH inhibitors have been developed^3–6^. However, the structure and catalytic mechanism of DHODH and the structural basis for on-target resistance to DHODH inhibitors is incompletely understood. This knowledge is critical to guide the development of drugs that can effectively target DHODH.

DHODH is embedded in the external surface of the inner mitochondrial membrane by an N-terminal domain that consists of a mitochondrial localization signal (MLS), a putative transmembrane α-helix (TM-α1), and two amphipathic α-helices (α2 and α3) (**Extended Data Fig. 1b**)^7–11^. The TM-α1 helix is not required for activity and is thought to stabilize the membrane interactions of DHODH^12^. The α2 and α3 helices form a ligand entry tunnel that extends from the membrane-embedded region of the enzyme to its active site. A loop structure (L1) connects the N-terminal domain to a C-terminal α/β-barrel that forms the active site. Flavin mononucleotide (FMN) sits on the floor of the active site and a second loop structure (L2) in the α/β-barrel serves as a ceiling over the active site.

Enzyme kinetic studies suggest that DHODH uses a two-step two-site double-displacement ‘ping-pong’ mechanism of catalysis (**Extended Data Fig. 1c**)^13,14^. In the first reaction, dihydroorotate (DHO) binds in the first redox site and is deprotonated by an active site serine, S214, which results in oxidation of DHO to orotate (ORO) and reduction of FMN to FMNH_2_^15^. ORO is then released and ubiquinone is recruited from the inner mitochondrial membrane, through the amphipathic tunnel, to the second redox site located on the opposite surface of FMN. In the second reaction, FMNH_2_ is re-oxidized to FMN with concomitant reduction of ubiquinone to ubiquinol. Ubiquinol then exits the tunnel.

There is a significant discrepancy between the over 100 published crystal structures of DHODH and its purported mechanism of catalysis. Because ubiquinone is highly hydrophobic, the structural studies were all performed using inactive enzyme in complex with different inhibitors^9,14,16–18^. These inhibitors all bind in the ubiquinone access tunnel, and the crystal structures all show ORO simultaneously occupying the first redox site. However, ‘ping-pong’ catalysis is predicated on the assumption that occupancy of the first and second redox sites are mutually exclusive^19^. This discrepancy has led to speculation that DHODH inhibitors induce a conformation of DHODH that does not reflect its ubiquinone-bound state^9,17^.

To determine whether our understanding of the structure of DHODH is incomplete, we solved the crystal structure of human DHODH in complex with decyl-ubiquinone (DCU), a biochemically active structural analog of ubiquinone (**Extended Data Fig. 1d-e**)^20^; and we solved the crystal structure of human DHODH bound to ORO with an empty ubiquinone access tunnel. We also used deep mutational scanning (DMS) to comprehensively map single amino acid substitutions in DHODH that confer resistance to one or both of two structurally distinct inhibitors, BAY2402234 and brequinar (**Extended Data Fig. 1f**)^18,21^. The DMS screens identified numerous mutations in the ubiquinone access tunnel and on the surface of DHODH at a distance from the tunnel that confer resistance by impairing drug binding. Taken together, these studies provide structural evidence for the ‘ping-pong’ model of catalysis and provide novel insights into the catalytic mechanism of DHODH. These studies also identify a pocket on the surface of DHODH that allosterically regulates the ubiquinone binding site. These findings have the potential to guide the development of new pharmacologic strategies to target DHODH.

### Structural characterization of human DHODH holoenzymes

To structurally characterize catalytically-active DHODH, we generated recombinant human wild-type DHODH protein lacking the MLS and TM-α1 helix (DHODH^WT-ΔN^) (**Extended Data Fig. 1b**). We confirmed that DHODH^WT-ΔN^ is catalytically active by dichlorophenolindophenol (DCIP) absorbance assay^22^, using a catalytic-dead S214 mutant (DHODH^S214A-ΔN^) as a negative control (**Extended Data Fig. 1g-h**). We then co-incubated DHODH^WT-ΔN^ with a saturating concentration of DCU and solved the crystal structure of DHODH^WT-ΔN^ in complex with DCU (DHODH-DCU) at 2.4 Å resolution (**Fig. 1a, Extended Data Fig. 2, and Extended Data Table 1**). DHODH-DCU exhibits the same overall secondary structure as previously published inhibitor- and ORO-containing ternary structures. The central α/β-barrel forms the first redox site and is connected to the N-terminal domain by the L1 loop (**indicted by a red arrow in Fig. 1a**). DCU binds in the ubiquinone access tunnel similarly to inhibitor but makes fewer contacts with the tunnel (**Fig. 1b and Extended Data Table 2**).

**Fig. 1:**
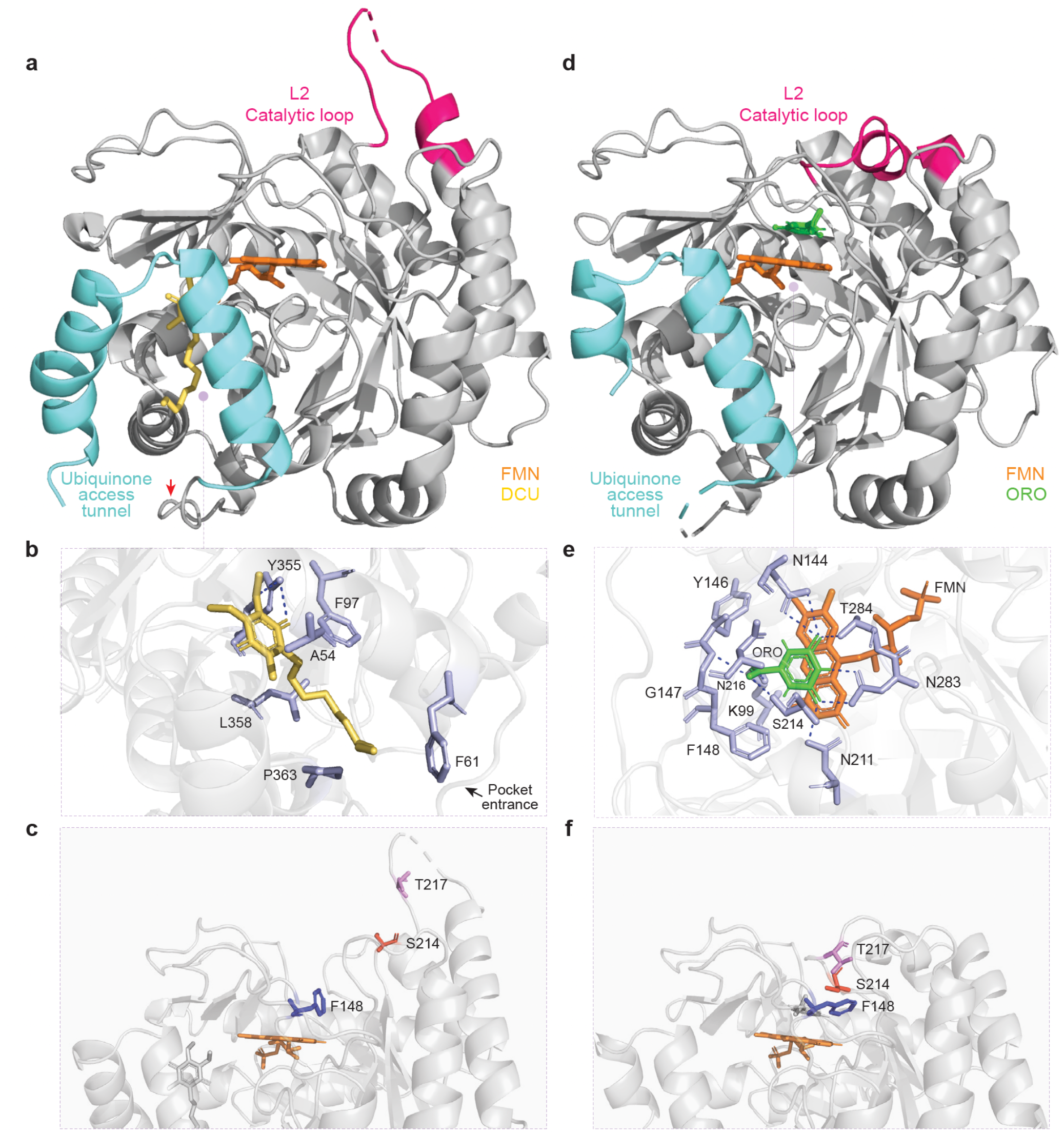
DHODH holoenzyme structures reveal that conformational changes in the loop over the first redox site are critical for catalysis. **(a)** Ribbon representation of DHODH-DCU showing the L2 loop over the empty first redox site in an ‘open’ conformation. The L1 loop is indicated by a red arrow. **(b)** Interactions of DCU with tunnel residues. DCU forms a hydrogen bond (dashed line) with Y355 and makes contacts with surrounding hydrophobic residues (cyan). **(c)** Position of catalytic residue S214 (red) and accessory residues T217 (pink) and F148 (blue) in DHODH-DCU. **(d)** Ribbon representation of DHODH-ORO showing the L2 loop in a ‘closed’ conformation. **(e)** Interactions of ORO with residue side-chains in the first redox site. ORO forms hydrogen bonds with K99, N144, G147, F148, N211, S214, N283 (violet; dashed lines). **(f)** Position of catalytic residue S214 (red) and accessory residues T217 (pink) and F148 (blue) in DHODH-ORO. The ubiquinone access tunnel (cyan), L2 loop (magenta), DCU (yellow), FMN (orange), and ORO (green) are highlighted. Ribbon representations were prepared using PyMOL. Protein-ligand interactions were determined using PLIP.

The most striking difference between DHODH-DCU and published DHODH-ORO-inhibitor structures is in the conformation of the L2 loop formed by residues 212 to 226. In DHODH-ORO-inhibitor structures, the L2 loop occludes the opening of the first redox site, with different loop residues being more or less resolved depending on which inhibitor is bound^9,14,16–18^. The structural heterogeneity of the L2 loop is presumed to reflect the fact that the loop has sufficient ‘breathing room’ to permit DHO to enter and ORO to exit the first redox site and to allow S214 to adopt an optimal conformation for catalysis^8,23,24^. In DHODH-DCU, the L2 loop over the unoccupied first redox site is in a wide-open conformation (**Fig. 1a**). This not only provides ample space for substrate to enter and exit the active site but also changes the position of S214 such that it juts out of the site (**Fig. 1c**). The open-loop conformation also moves another critical residue, T217, out of the active site. T217 is hypothesized to cooperate with F148 in the floor of the active site to enhance the basicity and reactivity of S214^9,25^. The overall effect of the open L2 loop appears to be to disable the first redox site when the site is unoccupied. Of note, DHODH-DCU recapitulates the hypothesized fifth step of the ‘ping-pong’ model of catalysis (**Extended Data Fig. 1c**).

Contrary to the ‘ping-pong’ model, published DHODH-ORO-inhibitor structures all show the two redox sites occupied simultaneously. To recapitulate these structures with DCU, we co-incubated DHODH^WT-ΔN^, DCU, and ORO in saturating crystallography solutions. Diffraction of the crystals failed to detect any ternary complexes but we were able to solve the structure of DHODH^WT-ΔN^ in complex with ORO alone (DHODH-ORO) at 3.0 Å resolution (**Fig. 1d, Extended Data Fig. 3, and Extended Data Table 1**). DHODH-ORO exhibits the same ‘closed’ L2 loop conformation as DHODH-ORO-inhibitor structures, with closure of the loop bringing S214 into close proximity to T217 and F148 (**Fig. 1e-f**). Notably, DHODH-ORO recapitulates the hypothesized third step of the ‘ping-pong’ model of catalysis (**Extended Data Fig. 1c**).

In DHODH-ORO-inhibitor structures, the α2- and α3-helices of the tunnel and the L1 loop are highly ordered, and the L1 loop occludes the opening of the tunnel^26^. By contrast, in DHODH-ORO, the proximal end of the α2-helix and the L1 loop are too disordered to be fully resolved (**Fig. 1d and Extended Data Fig. 3**). This suggests that substrate binding stabilizes the conformation of the tunnel and, moreover, that the structural dynamics of the tunnel are, at least in part, a function of tunnel occupancy. Notably, substrate binding to both redox sites of DHODH induces significant conformational changes in adjoining loop structures that appear to ‘close’ each reaction chamber prior to catalysis (**Extended Data Fig. 4**). This is consistent with previous reports that enzymes adopt ‘closed’ conformations during catalysis in order to create sealed reaction chambers that maximize the efficiency of electron transfer^23,27–30^.

It has been suggested that, with the exception of the ubiquinone access tunnel and the L2 loop over the first redox site, DHODH undergoes minimal conformational changes during catalysis^23,31^. However, examination of the surface representations of DHODH-DCU (**Extended Data Fig. 5**), DHODH-ORO (**Extended Data Fig. 6**), and a representative DHODH-ORO-inhibitor structure (**Extended Data Fig. 7**) reveals two previously unappreciated pockets on DHODH that extend deep into the core of the enzyme that have different conformations in the three complexes (**Fig. 2a-b and Extended Data Fig. 8**). Pocket 1, which is apparent in DHODH-DCU and in many DHODH-ORO-inhibitor structures but is effaced in DHODH-ORO, likely reflects an allosteric effect of tunnel occupancy. Pocket 2 is effaced in DHODH-ORO-inhibitor structures but remains accessible throughout the opening and closing of the first redox site in catalytically-active DHODH (**Extended Data Fig. 9**). We visualized pocket 2 in DHODH-DCU using the MOLEonline server (**Fig. 2c-d**)^32^. Pocket 2 is 22.5 Å long and extends from the surface of the enzyme to its catalytic core. The opening of the pocket is hydrophilic with a radius of 3.6 Å, and the pocket then becomes more hydrophobic and tapers to a bottleneck radius of 1.2 Å. This narrow aperture suggests that pocket 2 is unlikely to be a channel for movement of ligands into and out of the reaction site. However, it could serve as a binding site for allosteric regulators of DHODH activity.

**Fig. 2:**
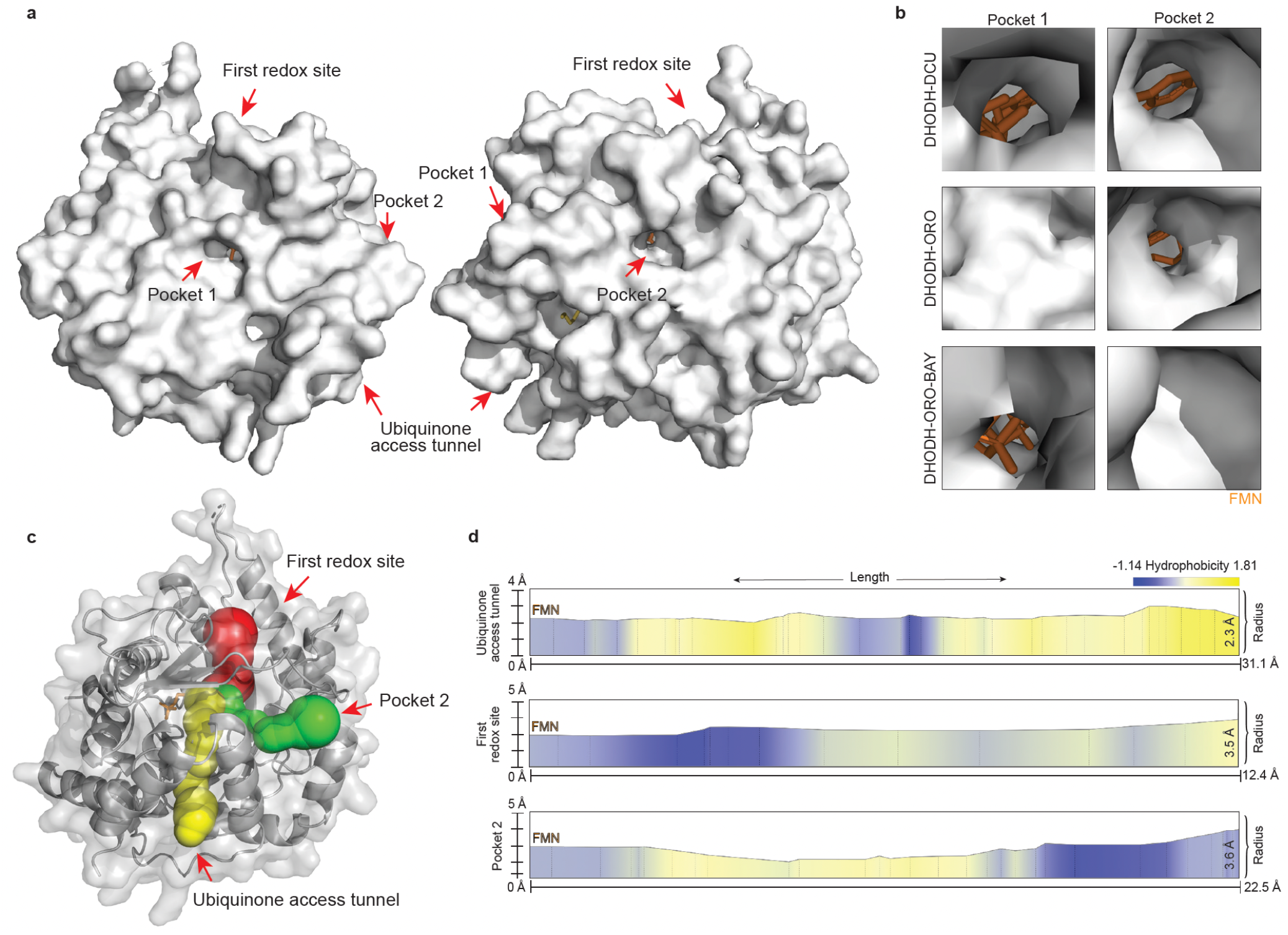
DHODH holoenzyme structures reveal cryptic surface pockets on DHODH that extend into the core of the enzyme. **(a)** Stereo-view surface representations of DHODH-DCU showing the locations of the first redox site, the ubiquinone access tunnel, pocket 1, and pocket 2, as indicated. The two views are rotated 180° around the vertical axis. **(b)** Close-up views of the locations corresponding to pocket 1 (left) and pocket 2 (right) in DHODH-DCU (top), DHODH-ORO (middle) and DHODH-ORO-BAY (PDB 6QU7; bottom). FMN (orange) is highlighted. Analogous data for other DHODH-ORO-inhibitor structures are presented in **Supplementary Fig. 8**. **(c)** Annotation of surface-accessible pockets on DHODH-DCU that extend into the FMN binding site. The first redox site (red), ubiquinone access tunnel (yellow), and pocket 2 (green) are shown. **(d)** Radii and hydrophobicities of each of the surface pockets from (c). Surface representations were prepared using PyMOL. Pocket annotation was performed using MOLEonline server.

### Systematic structure-function analysis of human DHODH

To complement our crystallographic studies, we undertook DMS drug resistance screens using BAY2402234 (BAY) and brequinar (BRQ) (**Extended Data Fig. 1f**)^18,21^. We generated a cDNA library encoding all 6,688 single amino acid variants of full-length wild-type human DHODH (DHODH^WT^) and expressed the library in TF-1 cells, an inhibitor-sensitive cell line. We then treated the cells for 8 days with doses of BAY or BRQ that are sufficient to kill TF-1 cells overexpressing DHODH^WT^ (**Extended Data Fig. 10**). The cytotoxicity of these doses of inhibitor can be fully rescued by supraphysiologic concentrations of the pyrimidine salvage substrate uridine, confirming that, at these high doses, the inhibitors act ‘on-target’. We then performed three independent replicates of each DMS screen and assessed the relative resistance of each variant to BAY or BRQ by determining the fold change of abundance of each variant compared to DMSO control. The read counts of each replicate of each screen correlated highly (**Extended Data Fig. 11**). Using a statistical threshold of enrichment of >3-fold, we identified numerous variants that confer resistance to one or both inhibitor (**Extended Data Fig. 12 and Extended Data Table 3**). Most of the resistance mutations localize to three distinct regions in DHODH: the ubiquinone/inhibitor-binding site, the first redox site, and the protein surface (**Fig. 3 and Extended Data Fig. 13**). We also identified resistance mutations that localize to the L2 loop and to the core of the α/β-barrel. Notably, at numerous residues, substituting any one of several structurally and chemically different amino acids confers resistance to BAY and/or BRQ (**Fig. 3a and 3f, Extended Data Fig. 13a and 13f, and Extended Data Table 4**).

**Fig. 3:**
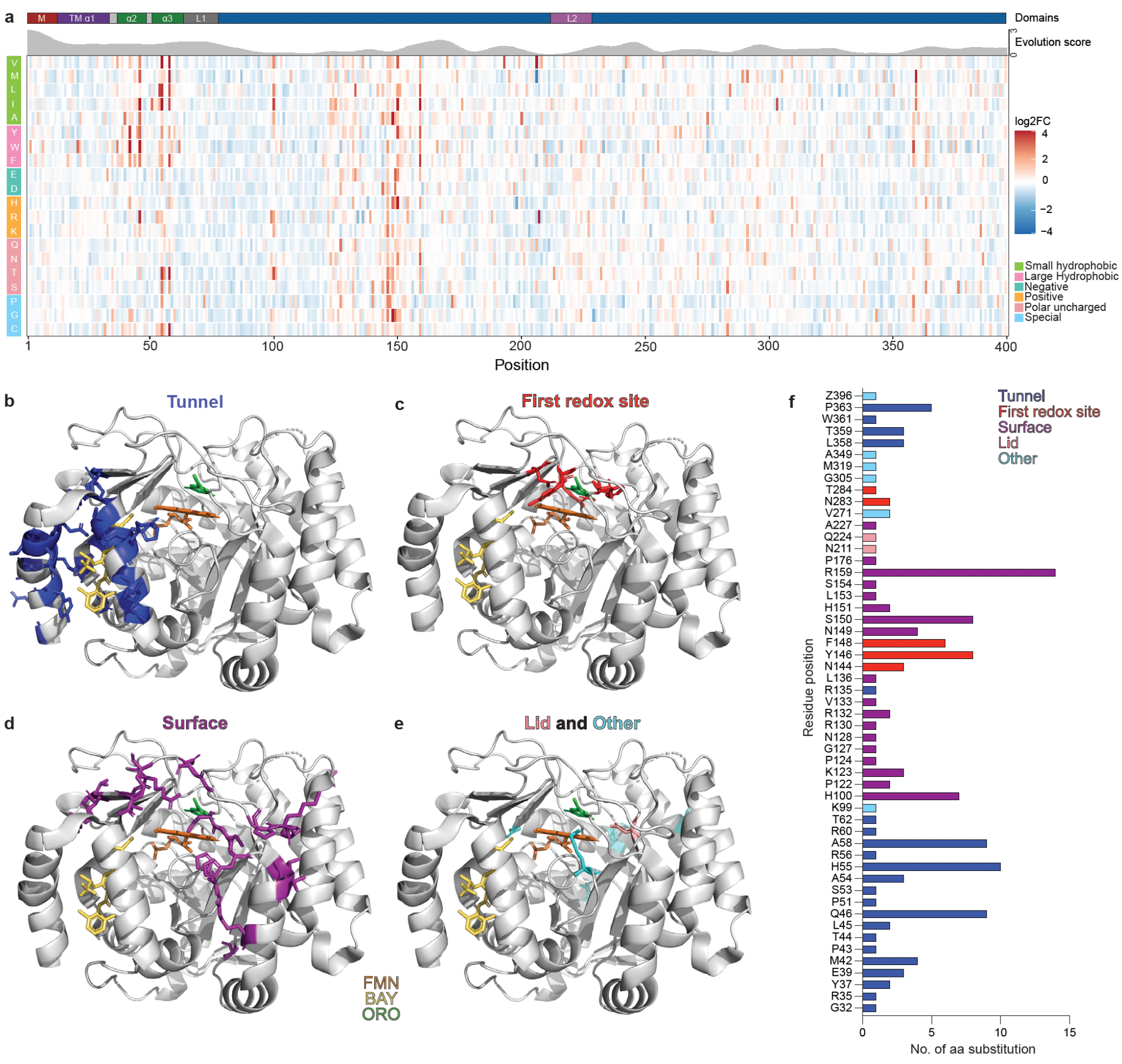
Structural mapping of BAY-resistant DHODH variants. **(a)** Heatmap showing the log2 fold change (log2FC) in variant enrichment (red) or depletion (blue) on day 8 of the BAY DMS screen relative to day 8 of the vehicle-control DMS screen. Each column represents an amino acid position and each row represents a substituted residue. Residues are grouped biochemically: small hydrophobic (green), large hydrophobic (pink), negatively charged (teal), positively charged (orange), polar uncharged (coral), and special (light blue). The locations in the DHODH secondary structure of the mitochondrial localization signal (M), transmembrane α1 helix (TM-α1), α2 and α3 helices, L1 loop, and L2 loop, are annotated above the heatmap. Shown below the annotation is the evolution score of each residue, as determined using Aminode. Lower scores indicate more conserved residues. **(b-e)** Ribbon representations of DHODH-ORO-BAY (PDB 6QU7) highlighting the locations of mutations that confer resistance to BAY in the ubiquinone access/drug-binding tunnel (blue) (b); first redox site (red) (c); surface (purple) (d); and lid (pink) and other locations (teal) (e). FMN (orange), BAY (yellow) and ORO (green) are highlighted. **(f)** Number of different substitutions that confer resistance to BAY at each indicated residue position. Locations of the residues in the 3-dimensional structure of DHODH are color-coded as in (b-e). Analogous data from the BRQ DMS screen are presented in **Supplementary Fig. 13**. Ribbon representations were prepared using PyMOL.

### Characterization of drug-binding-domain resistance mutations

Residue R135, located at the apex of the ubiquinone access tunnel (**Fig. 4a**), is one of only a handful of residues that are 100% conserved across a diverse array of DHODH orthologs (**Extended Data Fig. 14**), and structural studies suggest that R135 plays a critical role in catalysis by binding the ubiquinone head-group^9^. Nevertheless, multiple R135 substitutions were selectively enriched in the BRQ DMS screen, including the known loss-of-function mutant DHODH^R135C^ (**Extended Data Table 4 and Extended Data Fig. 13f**)^33–35^. *In-silico* protein free energy perturbation (FEP) calculations suggested that substitution R135C would impair BAY binding by 3.8 kcal/mol and BRQ binding by 15.5 kcal/mol (**Extended Data Table 5**)^36,37^. We cloned and characterized DHODH^R135C^ and found that, although DHODH^R135C-ΔN^ does have lower *in vitro* catalytic activity than DHODH^WT-ΔN^, it is selectively resistant to BRQ (**Fig. 4b**), and expression of full-length DHODH^R135C^ in cells confers selective resistance of cells to BRQ (**Fig. 4c-d**). To ask if R135C specifically impairs BRQ but not BAY binding, we optimized a thermal shift assay (TSA) for recombinant DHODH^ΔN^ proteins and a cellular thermal shift assay (CETSA) for full-length DHODH proteins (**Extended Data Fig. 15**)^38,39^. DHODH^R135C-ΔN^ aggregates at low temperatures and is not amenable to TSA. However, CETSA analysis found that BAY, but not BRQ, induces a significant DHODH^R135C^ thermal shift (**Fig. 4e**). Taken together, these results suggest that DHODH^R135C^ is a hypomorphic mutant that is sufficiently active to support *de novo* synthesis when endogenous DHODH is inhibited, and R135C confers resistance to BRQ by impairing BRQ binding.

**Fig. 4:**
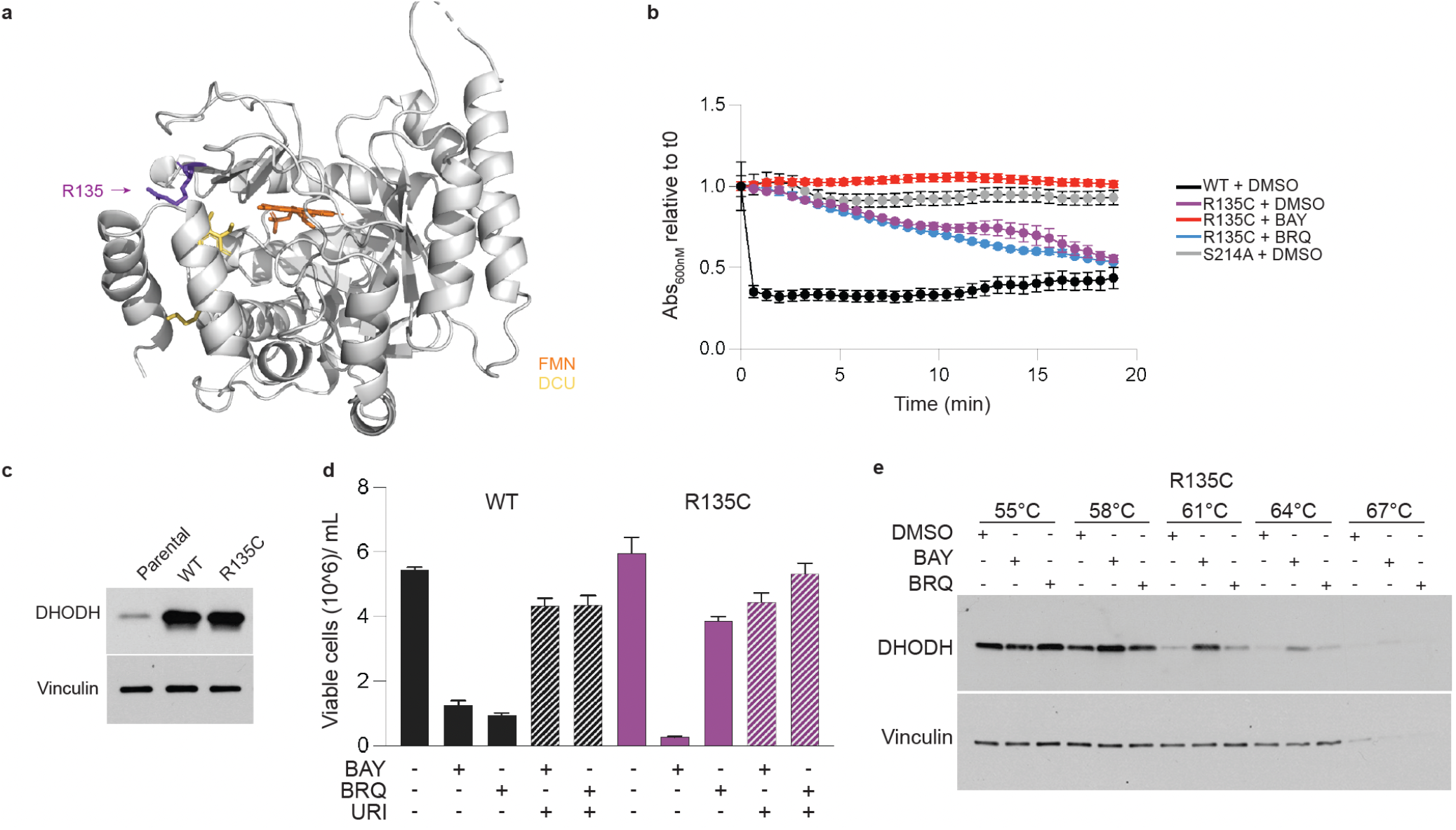
Loss-of-function mutation *DHODH^R135C^* impairs BRQ binding and confers resistance to BRQ. **(a)** Ribbon representation of DHODH-DCU showing the location of residue R135 (purple). FMN (orange) and DCU (yellow) are shown. **(b)** *In vitro* catalytic activity of recombinant DHODH^WT-ΔN^, DHODH^R135C-ΔN^, and DHODH^S214A-ΔN^ proteins in the presence of vehicle (DMSO), BAY 10 µM, or BRQ 10 µM, as indicated, as measured by DCIP assay. **(c)** Immunoblot analysis of DHODH (top) and vinculin (bottom) expression in parental TF-1 cells (Parental) and TF-1 cells expressing DHODH^WT^ (WT) or DHODH^R135C^ (R135C). **(d)** Growth of the cells from (c) after six-day treatment with DMSO, BAY 50 nM, or BRQ 5 µM, with or without uridine (URI) 50 µM, as indicated. **(e)** CETSA of lysates from TF-1 cells expressing DHODH^R135C^ treated *in vitro* with DMSO, BAY 50 nM, or BRQ 50 nM, as indicated. Immunoblots for DHODH (top) and vinculin (bottom) are shown. Shown are representative results from three or more independent experiments.

The DMS screens identified multiple substitutions at residue A58, located in the α3-helix, that confer resistance to both BAY and BRQ (**Fig. 3f, Extended Data Fig. 13f, and Extended Data Table 4**). FEP calculations of two of the substitutions, A58H and A58T, suggested that A58H would significantly impair BAY and BRQ binding, by 86.2 and 165.0 kcal/mol, respectively, whereas A58T would only modestly impair BRQ binding, by 1.9 kcal/mol, and would have no significant effect on BAY binding (**Extended Data Table 5**). We cloned and characterized the two substitutions. Consistent with the DMS screen results, DHODH^A58H^ is resistant to BAY and BRQ *in vitro* and in cells (**Extended Data Fig. 16a-c**), and consistent with the FEP calculations, neither BAY nor BRQ induce a significant DHODH^A58H^ thermal shift (**Extended Data Fig. 16d**). DHODH^A58T-ΔN^ is likewise resistant to BAY and BRQ *in vitro*, but both BAY and BRQ induce a DHODH^A58T-ΔN^ thermal shift that is comparable to the shift induced by binding of DCU, DHO or ORO to DHODH^WT-ΔN^ (**Fig. 5a-b**). Moreover, DHODH^A58T^ expression confers resistance of cells to both BAY and BRQ, but both inhibitors induce a DHODH^A58T^ thermal shift (**Fig. 5c-e**).

**Fig. 5:**
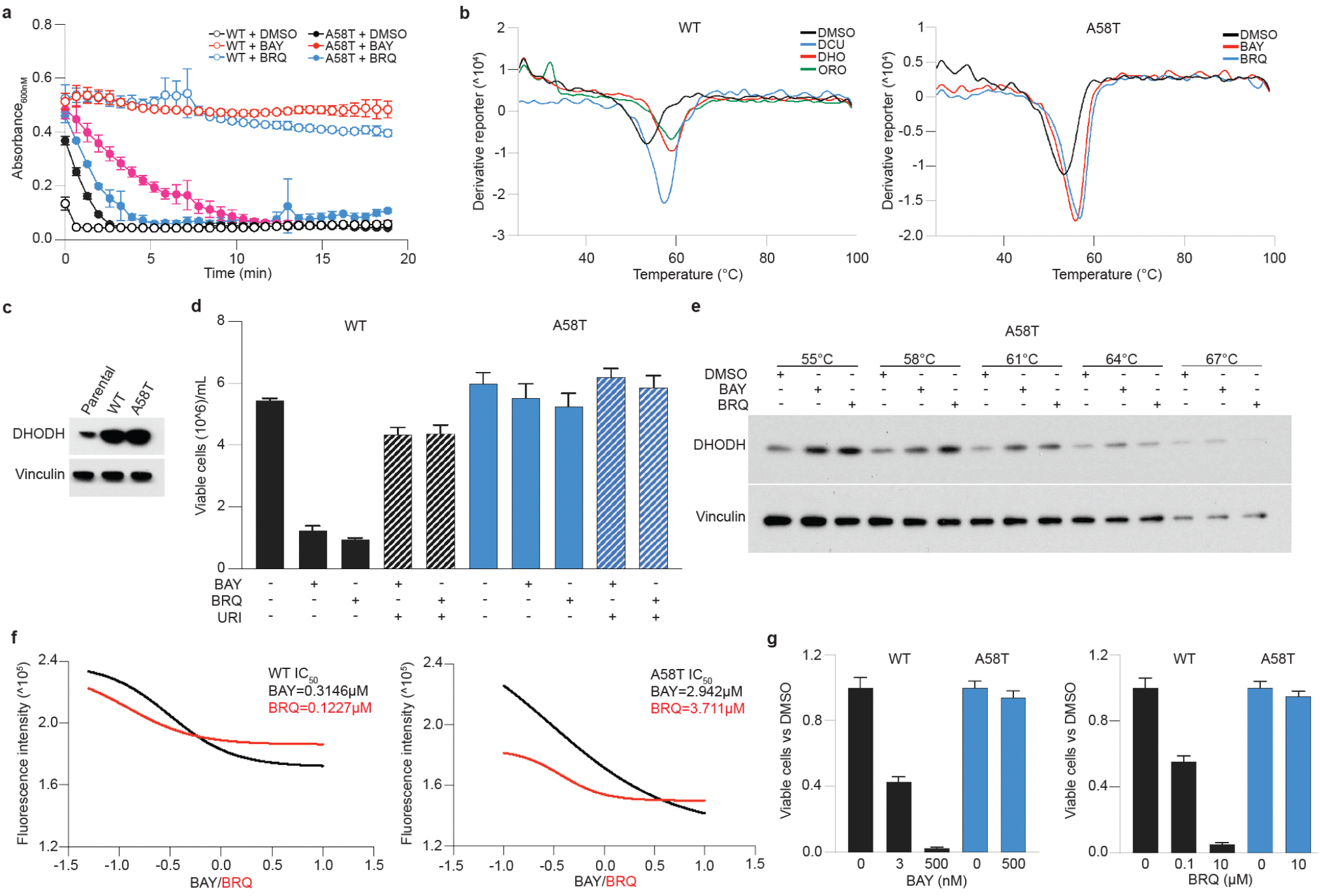
Drug-resistance mutation *DHODH^A58T^* partially impairs inhibitor binding and increases DCU binding. **(a)** *In vitro* catalytic activity of recombinant DHODH^WT-ΔN^ and DHODH^A58T-ΔN^ in the presence of vehicle (DMSO), BAY 10 µM, or BRQ 10 µM, as indicated, as measured by DCIP assay. **(b)** Left: TSA of DHODH^WT-ΔN^ treated with DMSO, DCU 100 μM (blue), DHO 100 μM (red), or ORO 100 μM (green), as indicated. Right: TSA of DHODH^A58T-ΔN^ treated with DMSO (black), BAY 10 μM (red), or BRQ 10 μM (blue), as indicated. **(c)** Immunoblot analysis of DHODH (top) and vinculin (bottom) expression in parental TF-1 cells (Parental) and TF-1 cells expressing DHODH^WT^ (WT) or DHODH^A58T^ (A58T). **(d)** Growth of the cells from (c) after six-day treatment with DMSO, BAY 50 nM, or BRQ 5 µM, with or without uridine 50 µM, as indicated. **(e)** CETSA of lysates from TF-1 cells expressing DHODH^A58T^ treated *in vitro* with DMSO, BAY 50 nM, or BRQ 50 nM, as indicated. Immunoblots for DHODH^A58T^ (top) and vinculin (bottom) are shown. **(f)** Inhibition curves of DHODH^WT-ΔN^ (left) and DHODH^A58T-ΔN^ (right) treated with a range of concentrations of BAY or BRQ, as indicated, as measured by OFA. IC_50_ values were calculated for each variant and each drug using the GraphPad Prism dose-response inhibition equation. **(g)** Growth of TF-1 cells expressing DHODH^WT^ (black) or DHODH^A58T^ (blue) after six-day treatment with the indicated concentrations of BAY (left) or BRQ (right). Shown are representative results from three or more independent experiments.

A58T is not the only substitution that confers resistance without abolishing inhibitor binding. Two substitutions at residue Q46, located in the α2-helix, behave similarly. Q46L was specifically enriched in the BRQ DMS screen and Q46R was specifically enriched in the BAY DMS screen (**Extended Data Fig. 12d**). DHODH^Q46L^ is selectively resistant to BRQ *in vitro* and in cells but both BAY and BRQ induce DHODH^Q46L^ thermal shifts (**Extended Data Fig. 17a-e**). Likewise, DHODH^Q46R^ is selectively resistant to BAY, but both inhibitors induce DHODH^Q46R^ thermal shifts (**Extended Data Fig. 17f-i**).

The observation that several DHODH mutants can bind inhibitors to which they are resistant begs the question, how do these mutants remain catalytically active in the presence of drug? Although all eukaryotic DHODH enzymes are quinone-dependent, gram-positive bacteria and archaea express DHODH orthologs that utilize soluble substrates, including fumarate and molecular oxygen, as terminal electron acceptors^2,40^. This prompted us to ask if drug-binding drug-resistant mutants acquire the ability to use fumarate or molecular oxygen as terminal electron acceptors. To address this question, we first had to develop a quinone-dependent *in vitro* assay for DHODH activity. The DCIP absorbance assay commonly used to quantify DHODH activity cannot be used to assess the quinone-dependence of DHODH because FMNH_2_ can directly reduce DCIP, and the assay is therefore not dependent on addition of a quinone to the reaction mixture (**Extended Data Fig. 18a-b**). To assess the quinone-dependence of drug-binding drug-resistant mutants, we modified an orotate fluorescence assay (OFA)^41^ to directly measure orotate production by DHODH. Using DHODH^WT-ΔN^ as a positive control and DHODH^S214A-ΔN^ as a negative control, we tested the ability of oxygen and fumarate to act as co-substrates of DHODH^A58T-ΔN^. In the absence of DCU, neither DHODH^WT-ΔN^ nor DHODH^A58T-ΔN^ have detectable OFA activity under ambient oxygen conditions, and fumarate cannot support orotate production by DHODH^WT-ΔN^ or DHODH^A58T-ΔN^ (**Extended Data Fig. 18c**). These results suggest that the mechanism of resistance of drug-binding mutants is not that they acquire the ability to use molecular oxygen or fumarate as terminal electron acceptors.

Drug resistance mutations that do not impair drug binding can act by either increasing the catalytic activity of the drug target or, in cases where the drug acts as a competitive inhibitor of an essential co-substrate, can increase the affinity of the target for its co-substrate^42–44^. Drug resistance mutations can also alter the structure of a drug target in such a way as to allow co-substrate binding even when drug is bound, effectively turning a competitive inhibitor into a non-competitive binder^45^. To ask if any of these mechanisms contribute to the drug resistance of DHODH^A58T^, we determined the maximum velocity (V_max_) and the K_m_ value for DCU of DHODH^WT-ΔN^ and DHODH^A58T-ΔN^ by OFA. DHODH^WT-ΔN^ has a V_max_ of 3.58 µM/sec and a DCU K_m_ value of 16.54 µM whereas DHODH^A58T-ΔN^ has a V_max_ of 2.62 µM/sec and a DCU K_m_ value of 9.8 µM (**Extended Data Fig. 19a-b**). This indicates that DHODH^A58T-ΔN^ has lower intrinsic catalytic activity but higher affinity for DCU than does DHODH^WT-ΔN^. Finally, we undertook to determine if inhibitor binding to DHODH^A58T-ΔN^ is non-competitive with DCU. The standard dose of DCU used for *in vitro* activity assays is significantly higher than the minimum DCU concentration required to support orotate production. We therefore repeated the OFA using a limiting concentration of DCU. Under DCU-limiting conditions, BAY acts as competitive inhibitor of both DHODH^WT-ΔN^ and DHODH^A58T-ΔN^ (**Extended Data Fig. 19c-d**). Notably, although A58T increases the *in vitro* IC_50_ values of BAY and BRQ by 10- and 30-fold respectively, DHODH^A58T^-expressing cells are completely resistant to 100-fold higher concentrations of BAY or BRQ than those that inhibit DHODH^WT^-expressing cells (**Fig. 5f-g**). The disproportionate *in cellulo* resistance of DHODH^A58T^ suggests that even a modest shift in the relative affinity of DHODH for inhibitor versus ubiquinone is sufficient to allow ubiquinone to outcompete inhibitor.

### Characterization of allosteric drug resistance mutations

The DMS screens identified numerous allosteric mutations that confer resistance to BAY and/or BRQ (**Fig. 3d-f and Extended Data Fig. 13d-f**). Allosteric resistance mutations have two known mechanisms of action. They can alter the conformation of the drug binding site at a distance^46,47^. Alternatively, in cases where the inhibitor selectively binds the inactive conformation of its target, the mutations can stabilize the active conformation, thereby locking the target in a low drug-binding-affinity state^48,49^. Among the allosteric resistance mutations identified in the DMS screens were multiple substitutions at four residues located in and around the cryptic pocket, pocket 2, we identified in DHODH-DCU and DHODH-ORO: H55, H100, S150, and R159 (**Fig. 6a and Extended Data Table 4**). Given that pocket 2 is ‘closed’ in DHODH-ORO-inhibitor structures and is ‘open’ in DHODH-DCU and DHODH-ORO (**Fig. 2b and Extended Data Fig. 8**), we hypothesized that pocket 2 mutations confer drug resistance either by allosterically impairing drug binding or by stabilizing the ‘open’ conformation of the pocket, *i.e.*, the active state of the enzyme.

**Fig. 6:**
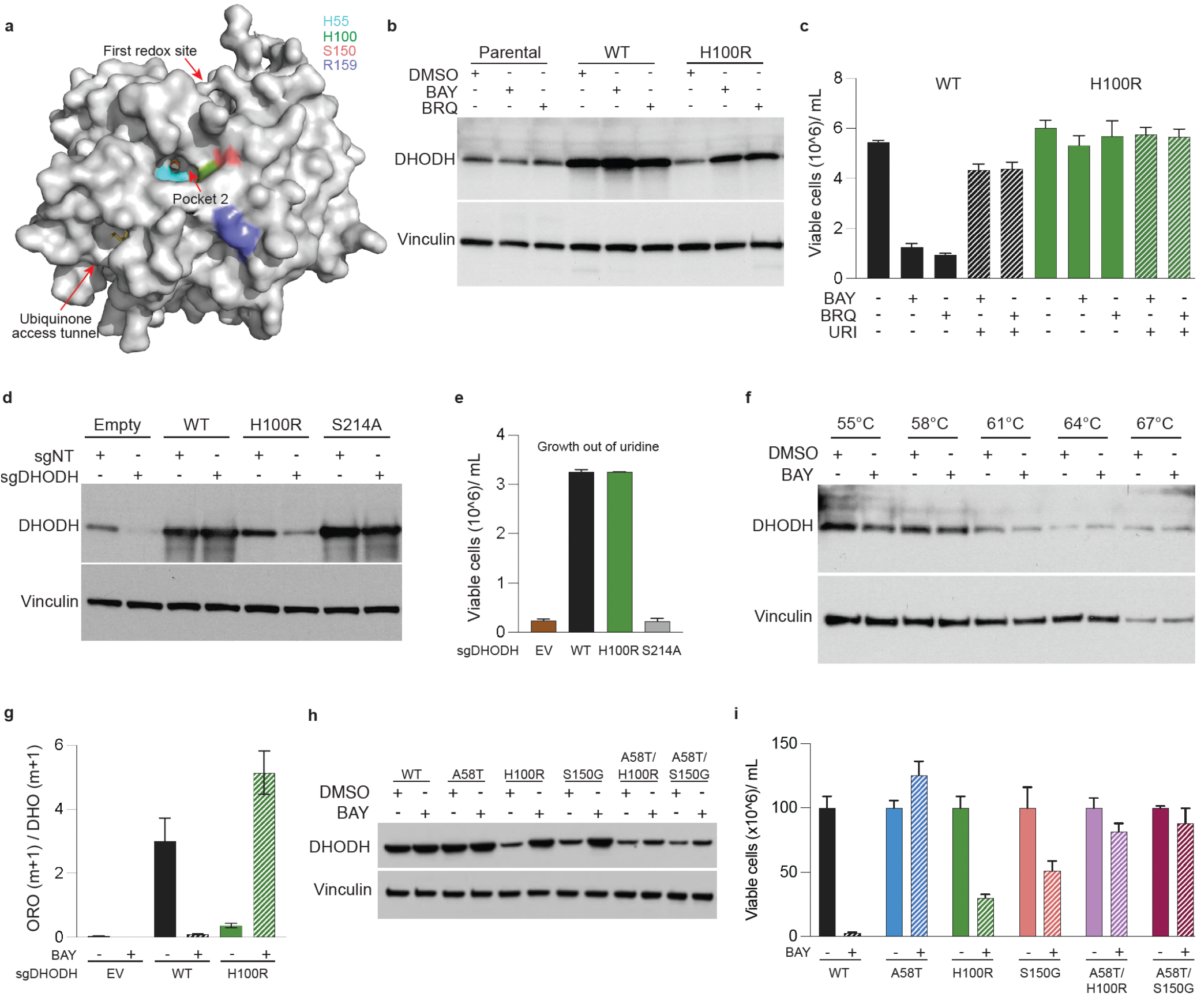
Allosteric drug-resistance mutations destabilize DHODH and impair drug binding. **(a)** Surface representation of DHODH-DCU highlighting the location of H55 (cyan), H100 (green), S150 (red), and R159 (violet) in relation to the ubiquinone access tunnel, pocket 1, and pocket 2 (red arrows). **(b)** Immunoblot analysis of DHODH (top) and vinculin (bottom) expression in parental TF-1 cells (Parental) and TF-1 cells expressing DHODH^WT^ (WT) or DHODH^H100R^ (H100R) following overnight treatment with vehicle (DMSO), BAY 50 nM, or BRQ 5 μM, as indicated. **(c)** Growth of the cells from (b) after six-day treatment with DMSO, BAY 50 nM, or BRQ 5 µM, with or without uridine (URI) 50 µM, as indicated. **(d)** Immunoblot analysis of DHODH (top) and vinculin (bottom) expression in TF-1 cells co-expressing empty vector (Empty) or an sgRNA-resistant cDNA encoding DHODH^WT^ (WT), DHODH^H100R^ (H100R), or DHODH^S214A^ (S214A), along with Cas9 and either a non-targeting control sgRNA (sgNT) or an sgRNA targeting DHODH (sgDHODH), as indicated. **(e)** Growth of the DHODH-knockout cells from (d) after six-day culture in media lacking supplemental uridine. **(f)** CETSA of lysates from TF-1 cells co-expressing DHODH^H100R^, Cas9, and an sgRNA targeting DHODH, treated *in vitro* with DMSO or BAY 50 nM, as indicated. Immunoblots for DHODH^H100R^ (top) and vinculin (bottom) are shown. **(g)** Quantification of DHODH activity in TF-1 cells co-expressing empty vector (Empty) or an sgRNA-resistant cDNA encoding DHODH^WT^ (WT) or DHODH^H100R^ (H100R), along with Cas9 and an sgRNA targeting DHODH (sgDHODH), following 16-hour pre-treatment with BAY 50 nM or DMSO, as measured by amide-^15^N glutamine tracing. Shown are orotate (m+1)/dihydro-orotate (m+1) ratios, as measured by LC-MS**. (h)** Immunoblot analysis of DHODH (top) and vinculin (bottom) expression in TF-1 cells expressing the indicated DHODH variants: wild-type (WT), DHODH^A58T^ (A58T), DHODH^H100R^ (H100R), DHODH^S150G^ (S150G), DHODH^A58T/H100R^ (A58T/H100R) or DHODH^A58T/S150G^ (A58T/S150G) following overnight treatment with DMSO or BAY 50 nM, as indicated. **(i)** Growth of the cells from (h) after six-day treatment with DMSO or BAY 50 nM. Shown are representative results from three or more independent experiments.

To assess the effect of perturbing pocket 2 on drug binding, we performed FEP calculations on four pocket 2 substitutions that were enriched in the BAY and BRQ DMS screens: H55Y, H100R, S150G, and R159M. Three of the substitutions, H55Y, H100R, and R159M, were calculated to modestly impair BAY binding, by 3.6, 0.9, and 0.2 kcal/mol, respectively, and BRQ binding, by 0.5, 3.25, and 0.4 kcal/mol, respectively (**Extended Data Table 5**). FEP analysis suggested that the S150G substitution would have no effect on binding of either drug.

To assess the effect of perturbing pocket 2 on the structural dynamics of DHODH, we performed DynaMut and protein message passing neural network (ProteinMPNN) analyses^50,51^. Either or both analyses suggested that H55Y, H100R, S150G, and R159M would destabilize DHODH protein (**Extended Data Table 6**). We cloned and further characterized DHODH^H100R^, which DynaMut and ProteinMPNN suggested would induce a ΔΔG free energy change of -1.25 and -18.6 kcal/mol, respectively. Consistent with the *in-silico* analyses, DHODH^H100R-ΔN^ is poorly expressed in bacteria (data not shown) and introduction of a DHODH^H100R^ expression construct in TF-1 cells does not result in an appreciable increase in DHODH expression when compared to parental TF-1 cells (**Fig. 6b**). However, when TF-1 cells harboring the DHODH^H100R^ cDNA are treated with BAY or BRQ, DHODH^H100R^ expression is upregulated to levels approaching those achieved in TF-1 cells ectopically expressing DHODH^WT^, and DHODH^H100R^-expressing cells are resistant to BAY and BRQ (**Fig. 6b-c**). DHODH^S150G^ and DHODH^R159M^ are likewise poorly expressed in cells, are induced by BAY or BRQ treatment, and confer resistance of cells to both drugs (**Extended Data Fig. 20**). Despite being calculated to be unstable by ProteinMPNN, DHODH^H55Y^ is expressed at levels similar as DHODH^WT^. Of note, we also performed DynaMut and ProteinMPNN analyses on the five drug-binding-domain substitutions we characterized, and ProteinMPNN erroneously calculated that two of substitutions, Q46L and A58H, would destabilize DHODH (**Extended Data Table 6**).

One explanation for the behavior of the unstable pocket 2 mutants is that there is a small fraction of mutant enzyme that folds spontaneously, is catalytically-active, and has low binding affinity for inhibitor; and inhibitor treatment binds and stabilizes, but does not induce folding of, the larger fraction of misfolded mutant protein. In this scenario, drug resistance is conferred by the spontaneously-folded mutant enzyme. An alternative explanation for the behavior of the unstable pocket 2 mutants is that inhibitor treatment induces folding of misfolded mutant protein into catalytically-active drug-resistant enzyme. To differentiate between these two possibilities, we introduced an sgRNA-resistant DHODH^H100R^ cDNA into TF-1 cells, cultured the cells in media containing uridine, and deleted endogenous DHODH from the cells. Untreated DHODH knock-out cells harboring the DHODH^H100R^ cDNA (DHODH^Δ/H100R^ cells) express very low but detectable levels of DHODH^H100R^, and the DHODH activity in untreated DHODH^Δ/H100R^ cells is sufficient to rescue the cells from the cytotoxic effects of uridine withdrawal (**Fig. 6d-e**). We then performed CETSA using lysates from untreated DHODH^Δ/H100R^ cells and found that BAY does not induce a significant thermal shift of spontaneously-folded DHODH^H100R^ protein (**Fig. 6f**). Taken together, these observations suggest that, in DHODH^Δ/H100R^ cells, there is a small fraction of DHODH^H100R^ protein that folds spontaneously, is catalytically active, and has impaired inhibitor binding.

Next, we sought to determine if ‘drug-induced’ DHODH^H100R^ protein is catalytically-active and drug-resistant. We reasoned that, if inhibitor treatment merely stabilizes misfolded DHODH^H100R^ protein, DHODH activity would be similar in inhibitor-treated and vehicle-treated DHODH^Δ/H100R^ cells. However, if inhibitor treatment induces folding of drug-resistant DHODH^H100R^ protein, DHODH activity would be higher in inhibitor-treated than vehicle-treated DHODH^Δ/H100R^ cells. To quantify cellular DHODH activity, we performed glutamine tracing experiments in DHODH^Δ/H100R^ cells after 16-hour pre-treatment with BAY or vehicle. We found that DHODH^Δ/H100R^ cells pretreated with BAY have significantly higher DHODH activity when compared to DHODH^Δ/H100R^ cells pretreated with vehicle (**Fig. 6g**). This indicates that inhibitor treatment increases both the level of DHODH^H100R^ protein and the level of DHODH^H100R^ activity in cells lacking endogenous DHODH.

Finally, we asked if inhibitor treatment promotes folding of misfolded DHODH proteins by directly binding and nucleating folding or by inducing the expression of endogenous chaperones. To distinguish between these two possibilities, we cloned a double-mutant DHODH variant that harbors both an H100R and an A58T substitution (DHODH^A58T/H100R^). We reasoned that, if endogenous chaperones non-specifically bind and induce folding of misfolded DHODH^H100R^ protein, then DHODH^A58T/H100R^ would be induced by inhibitor treatment. However, if direct binding of inhibitor to misfolded protein is required for folding, then further impairing inhibitor binding would prevent inhibitor-induced upregulation of DHODH^A58T/H100R^ expression. We found that BAY treatment does not significantly induce expression of DHODH^A58T/H100R^ (**Fig. 6h-i**). We obtained similar results with double-mutant DHODH^A58T/S150G^. These observations suggest that DHODH inhibitors can act as pharmacologic chaperones, directly binding and inducing folding of unstable DHODH mutants.

### Molecular dynamic simulations of DHODH^WT^, DHODH^A58T^, and DHODH^H100R^

None of the drug-resistant mutants we functionally characterized, including DHODH^A58T^, can be expressed and purified sufficiently to perform crystallography studies. We therefore undertook molecular dynamic (MD) simulations to gain insights into the structural effects of two of the substitutions, A58T and H100R. First, we modeled the consequences of A58T on ligand binding in the ubiquinone access tunnel using fixed-backbone mutagenesis, evaluating our DHODH-DCU structure alongside DHODH-ORO-BAY (PDB 6QU7) and DHODH-ORO-BRQ (PDB 1D3G)^9,18^. We found that the bulkier threonine sidechain in DHODH^A58T^ constricts the tunnel and clashes with DCU, BAY, and BRQ in each of their respective structures (**Fig. 7a**). This suggests that, for DHODH^A58T^ to be catalytically active, either ubiquinone needs to be positioned differently in the tunnel or the α-helices of the tunnel need to rearrange to permit ubiquinone binding.

**Fig. 7:**
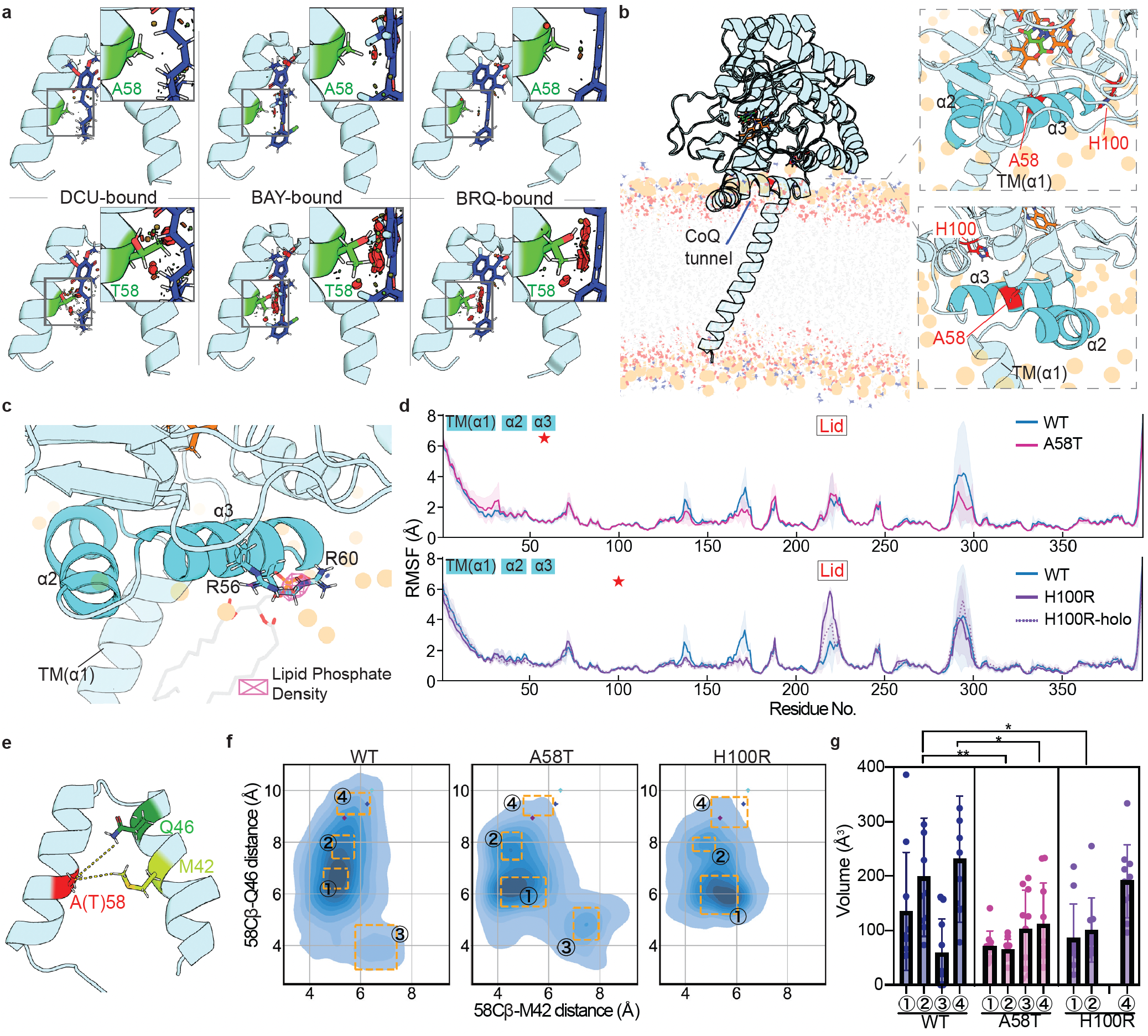
MD simulation of DHODH^WT^, DHODH^A58T^ and DHODH^H100R^. **(a)** Ubiquinone access tunnel structures of DHODH^WT-ΔN^ (top) or DHODH^A58T-ΔN^ (bottom) in different ligand-bound states: DCU-bound (left), BAY-bound (center), or BRQ-bound (right). Potential clashes with ligands are indicated by red discs. DHODH^WT-ΔN^ structures are experimentally determined X-ray crystal structures obtained in this study (DCU-bound) and previous studies (BAY-bound, 6QU7; BRQ-bound, 1D3G). DHODH^A58T-ΔN^ conformations were modeled using fixed-backbone mutagenesis (PyMOL) of DHODH^WT-ΔN^ structures without minimization. **(b)** A representative frame from an MD simulation of full-length DHODH^WT^ embedded in a palmitoyl-oleoyl-phosphatidylethanolamine (POPE) bilayer (left) and zoomed-in snapshots of the same MD simulation showing two different views of the membrane-embedded helices (right). POPE phosphates (tan spheres), POPE fatty acid chains (red, oxygen atoms; gray, aliphatic atoms), FMN (orange), orotate (green), α2 and α3 helices (cyan), and residues A58 and H100 (red) are indicated. **(c)** Averaged volumetric density of POPE phosphates at the α3-helix of DHODH^WT^ (n=3, 1-μs simulations), represented as a pink mesh. One representative frame is aligned. **(d)** Top: average root-mean-square-fluctuations (RMSF) of inhibitor-free DHODH^WT^ (WT) overlaid with inhibitor-free DHODH^A58T^ (A58T). Bottom: RMSF of inhibitor-free DHODH^WT^, inhibitor-free DHODH^H100R^ (H100R), and inhibitor-bound DHODH^H100R^ (H100R). The data were derived from three 1-μs simulations for each variant, with standard deviations overlaid (transparent shading). Cyan blocks indicate residues that form the transmembrane helix (TM-α1) and the membrane-embedded helices (α2 and α3). The boxed red “Lid” text indicates residues that form the L2 loop. Red asterisks indicate mutated residues A58 and H100. **(e)** Schematic cartoon illustrating the locations of Q46 (dark green), M42 (light green) and A58 or T58 (red) in the ubiquinone access tunnel. **(f)** Contour plots of tunnel width as measured by distances between the β-carbon of residue 58 (58Cβ) and residue M42 (58Cβ-M42) (X-axis) or between 58Cβ and residue Q46 (58Cβ-Q46) (Y-axis) taken every 0.2 ns during simulations (n=3) of inhibitor-free DHODH^WT^ (left), inhibitor-free DHODH^A58T^ (center), and inhibitor-free DHODH^H100R^ (right). The 58Cβ-M42 and 58Cβ-Q46 distances determined from DHODH crystal structures are indicated by dots: purple, DHODH-ORO; cyan, DHODH-DCU; blue, DHODH-ORO-BRQ (PDB 1D3G). Unique tunnel conformations observed in the contour plots are indicated by orange boxes (labeled 1, 2, 3, and 4). **(g)** Cavity volumes of tunnel locations highlighted in (f) in 10 representative frames from each cluster. Average volumes (bar plots) and standard deviations (error bars) are indicated. Statistical significance was assessed by Welch’s t-test between corresponding clusters in different variants (*p ≤ 0.05, **p ≤ 0.01).

To better understand the effects of A58T and H100R on inhibitor binding and to model the impact of the substitutions on the overall structure of DHODH in the context of its membrane microenvironment, we constructed full-length DHODH models and conducted all-atom MD simulations in simplified lipid 1-palmitoyl-2-oleoyl-sn-glycero-3-phosphethanolamine (POPE) bilayers (**Fig. 7b**). To build full-length membrane-embedded DHODH models, we used AlphaFold to predict the conformation of the missing TM-α1 helix and combined it with known DHODH^WT-ΔN^ crystal structures. We first determined the most stable membrane orientation of our modeled full-length wild-type structure (DHODH^WT-FL^) using simulations starting from four distinct membrane-oriented geometries (**Extended Data Fig. 21a**). Regardless of the initial configuration, DHODH^WT-FL^ converges to a similar conformation and membrane-embedding depth across >15 replicate 1-µs simulations, indicating a stable and preferred membrane geometry (**Extended Data Fig. 21b-d**). These conformations contrast with results of a previous MD simulation study that reported that portions of the α2 and α3 helices are lifted away from the membrane surface and are unstable^23^, but they agree with the conformation predicted by the PPM 3.0 server, wherein the α2 and α3 helices are stably associated with the membrane surface^52^.

Using the most stable predicted membrane geometry, we next performed triplicate 1-µs simulations of DHODH^WT-FL^, DHODH^A58T-FL^ and DHODH^H100R-FL^. Density analysis of POPE phosphate groups identified multiple residues that contribute to membrane interactions. Although individual lipid contacts varied across replicates (**Extended Data Fig. 21e**), modeling of all three variants consistently captured stable lipid interactions with the RxxxR motif (residues 56 to 60) of the α3 helix (**Fig. 7c and Extended Data Fig. 21f**). This suggests that R56 and R60 create a strong phospholipid head-group binding site. In addition, in simulations of DHODH^H100R-FL^, mutant residue R100 directs its sidechain down towards the phospholipid head-group layer of the membrane, resulting in gain-of-interaction lipid density (**Extended Data Fig. 21f**). This suggests that, whereas H100 in DHODH^WT-FL^ contributes to an intra-protein hydrogen bond network that stabilizes pocket 2 (**Extended Data Fig. 22a**), R100 in DHODH^H100R-FL^ trades this stabilization for additional lipid interactions. Although DHODH^A58T-FL^ and DHODH^H100R-FL^ vary in their lipid engagements, the overall protein orientation relative to membrane is similar between DHODH^WT-FL^, DHODH^A58T-FL^, and DHODH^H100R-FL^ (**Extended Data Fig. 21g**).

To determine if A58T or H100R induces allosteric effects or alters local protein stability, we analyzed root-mean-square fluctuations of DHODH^WT-FL^, DHODH^A58T-FL^, and DHODH^H100R-FL^. DHODH^WT-FL^ and DHODH^A58T-FL^ exhibit near-identical conformations and overall dynamics in their soluble domains (**Fig. 7d, top**). In contrast, DHODH^H100R-FL^ displays increased flexibility compared to DHODH^WT-FL^, particularly in the L2 loop residues 215 to 225 (**Fig. 7d, bottom**). Consistent with our observation that inhibitor treatment stabilizes DHODH^H100R^, inhibitor-bound DHODH^H100R-FL^ simulations generated from DHODH-ORO-BRQ show reduced lid flexibility compared to inhibitor-free simulations. In DHODH^WT-FL^ and DHODH^A58T-FL^, H100 forms stable interactions with H151, whereas K99 dynamically alternates between interactions with ORO and the H100–H151 pair (**Extended Data Fig. 22a-c**). In contrast, in DHODH^H100R-FL^, K99 adopts a different conformation that decreases its interactions with ORO and H151 (**Extended Data Fig. 22a-c**), This results in decreased ORO binding and dissociation of ORO from its binding site (**Extended Data Fig. 22b**). Our simulations predict that the overall effect of H100R is to disrupt local interaction networks within pocket 2 involving ORO and neighboring residues, which reduces ORO binding stability and increases lid flexibility. Inhibitor binding decreases the dynamics within pocket 2 and favors a more wild-type-like interaction network by bringing H55 in the α3 helix in closer proximity to K99 (**Extended Data Fig. 22b**). This results in reduced lid flexibility and stabilized ORO engagement. These results provide an explanation for the inherent instability of DHODH^H100R^ and for the observation that DHODH^H100R^ is stabilized upon inhibitor binding. Although H100R does impact local interaction networks, neither H100R nor A58T induces large-scale backbone rearrangements on the 1-µs timescale.

Finally, we analyzed ubiquinone access tunnel conformations in DHODH^WT-FL^, DHODH^A58T-FL^, and DHODH^H100R-FL^ to assess the potential impact of the mutations on inhibitor binding. Tunnel width was quantified by measuring the distances between residue 58 (either alanine or threonine) and two residues, M42 and Q46, located in the opposing α2 helix (**Fig. 7e**). Tunnel volume was quantified by measuring solvent-accessible cavity size. Across simulations, we identified several distinct tunnel conformations that vary in relative occupancy between wild-type and mutant protein ensembles (**Fig. 7f and Extended Data Fig. 22d**). Although the most populated conformations of DHODH^WT-FL^, DHODH^A58T-FL^, and DHODH^H100R-FL^ are not drastically different (*e.g*., in tunnel volume), on average, drug-resistant mutants more frequently adopt narrower tunnel geometries, particularly in the second, third and fourth most populated conformations (**Fig. 7g**). Interestingly, tunnel geometries of simulated full-length proteins in the context of a biologically realistic membrane are significantly narrower than tunnel geometries observed in experimentally-derived crystal structures of soluble truncated inhibitor-bound and inhibitor-free proteins (**Fig. 7f**). This indicates that, contrary to previous studies that suggested that the membrane microenvironment enhances tunnel flexibility^26,53^, membrane association may actually constrain tunnel geometry. This observation provides new context for our understanding of the structural mechanisms that mediate drug resistance and advocates for consideration of a contribution of the mitochondrial membrane to drug resistance. With that in mind, we aligned representative DHODH^A58T-FL^ frames to inhibitor-bound structures and found that, consistent with our observation that DHODH^A58T-ΔN^ has a higher affinity for DCU than does DHODH^WT-ΔN^, DCU is easily accommodated in the DHODH^A58T-FL^ tunnel and faces fewer potential steric clashes compared to BAY or BRQ, especially in the narrower conformation 2 and conformation 3 (**Extended Data Fig. 22e**). This suggests that the impact of resistance mutations on ligand binding is smaller for DCU than for inhibitors (**Extended Data Fig. 22e**). The fact that certain tunnel conformations discourage inhibitor binding but can easily accommodate quinone binding could explain why DHODH^A58T^ is significantly more drug resistant in cells than *in vitro*.

## Discussion

Herein, we report the crystal structures of two DHODH holoenzymes, DHODH bound to the biochemically active quinone co-substrate DCU and DHODH bound to its reaction product ORO. Unlike previously reported crystal structures of inactive DHODH complexes, these holoenzyme structures fully recapitulate the ‘ping-pong’ model of catalysis. Notably, our DHODH-DCU structure is the first crystal structure to show DHODH with an empty first redox site. Previous DHODH-ORO-inhibitor crystal structures and our DHODH-ORO structure all show ORO occupying the first redox site, which is sealed closed by the L2 loop. In DHODH-DCU, the L2 loop is in a wide-open conformation. It is likely that the high degree of structural similarity between DCU and ubiquinone allowed us to capture a conformation of DHODH that clearly demonstrates the ‘lid’ function of the L2 loop. The distinct conformations of the L2 loop in DHODH-DCU and DHODH-ORO also suggest that the position of the lid determines the catalytic competence of DHODH. Lid opening disables DHODH by shifting key catalytic residues away from the active site, including S214, which is displaced 14 Å away from the floor of the site. Lid closure restores the catalytic competence of DHODH by bringing S214 into close proximity to the active site and accessory catalytic residues. These observations suggest that the L2 loop not only controls substrate access to the first redox site but also plays a critical role in regulating the activity of DHODH.

DHODH has, for decades, been investigated as a therapeutic target in inflammatory diseases, cancer, malaria, and other ailments^2,5,6^. These efforts have largely focused on targeting the ubiquinone access tunnel because, in addition to being a deep pocket that is amenable to small molecule docking, the tunnel has a highly variable amino acid sequence that makes species-specific targeting feasible. Despite extensive efforts, only two human-specific DHODH inhibitors, leflunomide and teriflunomide, are currently in clinical use, for the treatment of rheumatoid and psoriatic arthritis and multiple sclerosis^54–57^. Several plasmodium-specific DHODH inhibitors have been developed that potently suppress multi-drug-resistant malarial parasitemia^58–60^. However, they failed to advance clinically due to the rapid emergence of drug resistance^61–63^. Our studies suggest that human DHODH is similarly prone to developing resistance to drugs that act as competitive inhibitors of ubiquinone. This is perhaps not surprising. The high degree of variability in the quinone binding sites of DHODH orthologs suggests that either quinones do not have rigid binding conformations and can tolerate significant structural variability in their binding sites, or that the quinone access tunnel is inherently flexible.

One example of a substitution that significantly alters the ubiquinone access tunnel but is sufficiently well-tolerated to confer drug-resistance is substitution A58T. Although the bulky threonine side-chain in DHODH^A58T^ constricts the tunnel and decreases the affinity of DHODH for inhibitor, it also increases the affinity of DHODH for DCU. It is impossible to know the relative contribution of each of these mechanisms to the drug-resistance of DHODH^A58T^. However, the fact that DHODH^A58T^ is significantly more drug-resistant in cells than *in vitro* suggests that the enhanced affinity of DHODH^A58T^ for ubiquinone plays a role in drug resistance. The only previously reported example of a drug-resistance mutation that acts by increasing the affinity of an enzyme for its co-substrate is the ‘gatekeeper’ T790M mutation in epidermal growth factor receptor (EGFR)^44^. T790M confers resistance to tyrosine kinase inhibitors (TKI) by both decreasing the affinity of EGFR for drug and increasing the affinity of EGFR for ATP. The drug-resistance of EGFR^T790M^ can be successfully overcome by irreversible TKI inhibitors that form a covalent bond with cysteine-797 in the ATP binding site^64^. It is notable that, in the case of DHODH, such a strategy would not be possible as there is not a single cysteine residue in human DHODH.

Our studies, along with the failure of efforts to develop effective plasmodium-specific DHODH inhibitors, suggest that targeting quinone binding is not an effective way to inhibit DHODH in diseases such as cancer and microbial infections where there is a strong selective pressure for drug resistance. Our finding that the L2 loop opens and closes during the catalytic cycle of DHODH and that closure of the loop is required for catalysis suggests that hindering lid movement would inhibit DHODH activity. Moreover, the fact that many L2 loop residues are highly conserved (**Extended Data Fig. 14 and 23**) could indicate that the catalytic lid is less tolerant to mutations than the ubiquinone access tunnel. It is also notable that G225, a hinge residue in mammalian DHODH orthologs that is critical for the sharp bending required to open and close the catalytic lid, is not conserved in microbial DHODH orthologs, including malaria (**Extended Data Fig. 14**). This indicates that it may be possible to develop species-selective inhibitors that target the catalytic lid. Another potential alternative site for small molecule targeting of DHODH is pocket 2 on the surface of DHODH. Our structure-function data suggest that pocket 2 allosterically regulates the ubiquinone access tunnel, and it may be possible to target pocket 2 in such a way as to impair ubiquinone binding and inhibit DHODH activity. Given that many of the residues around pocket 2 are highly conserved (**Extended Data Fig. 14 and 23b**) and that mutating the region around pocket 2 destabilizes DHODH, pocket 2 may be relatively resistant to acquisition of drug resistance mutations.

Taken together, these studies provide novel insights into the catalytic mechanism of DHODH and identify novel potentially targetable sites in DHODH that could enhance the efficacy of DHODH inhibitors.

---

**Extended Data Fig. 1:**
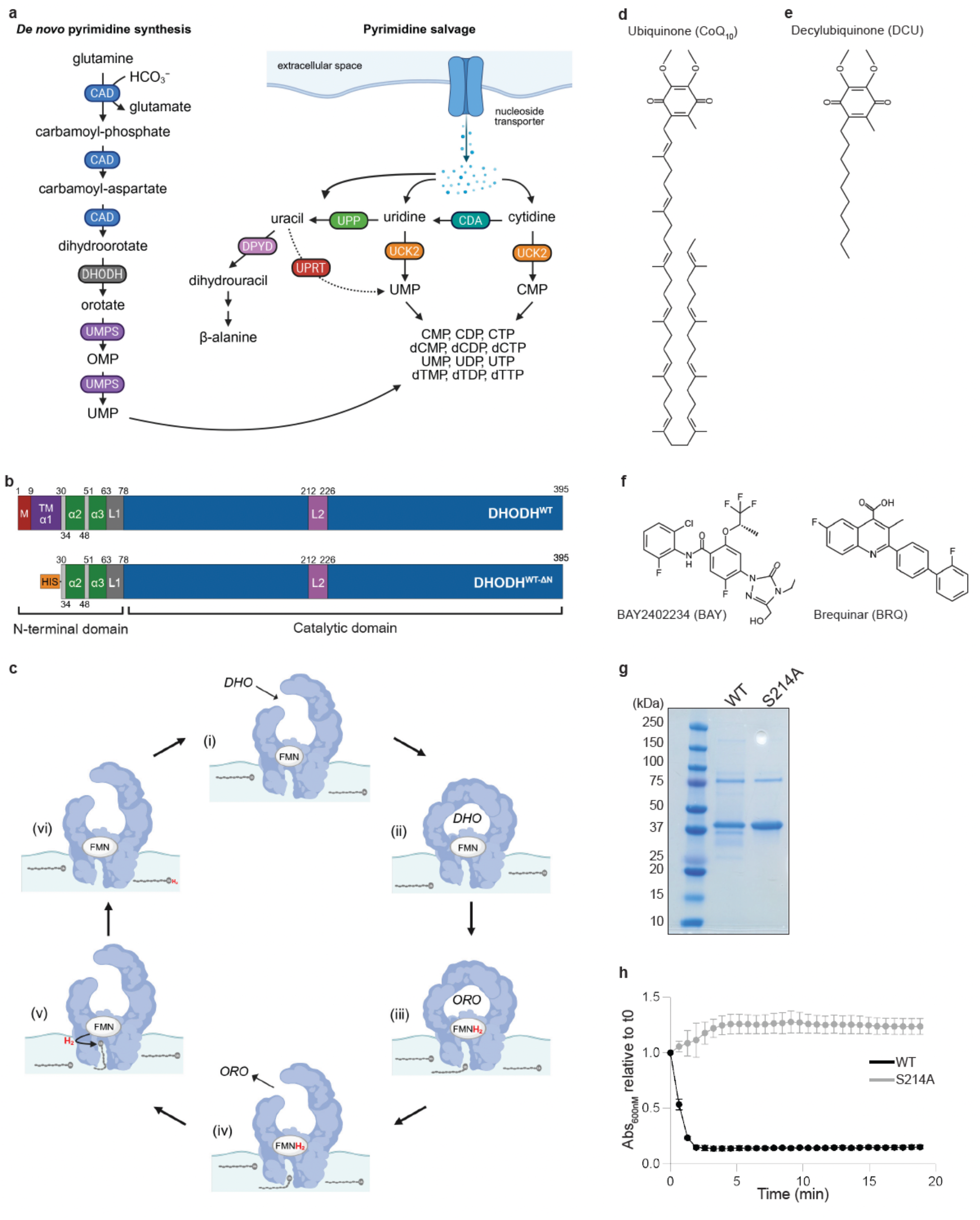
The reaction catalyzed by DHODH. **(a)** Schematic representation of the *de novo* and salvage pathways by which cells acquire pyrimidines. DHODH catalyzes the fourth and rate-limiting step in the *de novo* pathway, converting dihydroorotate to orotate. **(b)** Schematic representation of the domain structure of full-length human DHODH (top) and the soluble N-terminal-truncated recombinant human DHODH protein used for *in vitro* studies (bottom). M, mitochondrial localization signal; TM-α1, transmembrane helix; α2 and α3, α-helices that form the ubiquinone access tunnel; L1, loop structure that connects the N-terminal membrane-associated domain to the C-terminal catalytic domain; L2, loop structure that overlays the first redox site. **(c)** Schematic representation of the ‘ping-pong’ double-displacement model of catalysis. In the first half-reaction, dihydroorotate (DHO) binds in the first redox site (i), the reaction chamber is sealed (ii), and DHO is oxidized to orotate (ORO) and flavin mononucleotide (FMN) is reduced to FMNH_2_ (iii). In the second half-reaction, after ORO is released, ubiquinone binds in the second redox site (iv), FMNH_2_ is oxidized to FMN and ubiquinone is reduced to ubiquinol (v), and ubiquinol is released (vi). **(d-f)** Chemical structures of ubiquinone, the physiologic co-substrate of DHODH (d), decyl-ubiquinone, the *in vitro* co-substrate of DHODH used for *in vitro* studies (e), and BAY 2402234 (BAY) and brequinar (BRQ), two DHODH inhibitors that acts as competitive inhibitors of ubiquinone (f). **(g)** Coomassie blue-stained SDS-PAGE analysis of purified recombinant N-terminal-truncated wild-type (WT) and catalytic-dead mutant (S214A) DHODH. **(h)** *In vitro* catalytic activity of the recombinant DHODH variants from (g), as measured by DCIP assay. Panels (a-f) were generated using BioRender.

**Extended Data Fig. 2: Structure of human DHODH holoenzyme bound to DCU.** Rotating ribbon representation of DHODH in complex with DCU. The catalytic lid over the first redox site (pink) is in an ‘open’ conformation and DCU (yellow) occupies the space between the α2 and α3 helices that forms the ubiquinone access tunnel (cyan). FMN (orange) is shown. The ribbon diagram was prepared using PyMOL. [FILM]

**Extended Data Fig. 3: Structure of human DHODH holoenzyme bound to ORO.** Rotating ribbon representation of DHODH in complex with ORO. ORO (green) occupies the first redox site, the catalytic lid over the first redox site (pink) is in a ‘closed’ conformation, the ubiquinone access tunnel (cyan) is empty, and the proximal end of the α2 helix of the ubiquinone access tunnel and the L1 loop are unstructured. FMN (orange) is shown. The ribbon diagram was prepared using PyMOL. [FILM]

**Extended Data Fig. 4: The L2 loop over the first redox site and the α-2 helix of the ubiquinone access tunnel are highly flexible in holo-DHODH.** Film-overlay of the ribbon representations of DHODH-DCU and DHODH-ORO showing the movement of the L2 loop over the first redox site (pink) and the movement of the resolved portions of the α-helices of the ubiquinone access tunnel (cyan). [FILM]

**Extended Data Fig. 5: Surface conformation of the DHODH-DCU holoenzyme.** Rotating surface representation of DHODH-DCU. The L2 loop over the first redox site (pink) is in an ‘open’ conformation and DCU (yellow) occupies the space between the α2 and α3 helices that forms the ubiquinone access tunnel (cyan). FMN (orange) is shown. The surface diagram was prepared using PyMOL. [FILM]

**Extended Data Fig. 6: Surface conformation of the DHODH-ORO holoenzyme.** Rotating surface representation of DHODH-ORO. ORO (unseen) occupies the first redox site, the catalytic lid over the first redox site (pink) is in a ‘closed’ conformation, and the ubiquinone access tunnel (cyan) is empty. The proximal end of the α2 helix of the ubiquinone access tunnel is unstructured. FMN (orange) is shown. The surface diagram was prepared using PyMOL. [FILM]

**Extended Data Fig. 7: Surface conformation of a representative DHODH-ORO-inhibitor ternary complex.** Rotating surface representation of DHODH-ORO-BAY (PDB 6QU7). ORO (unseen) occupies the first redox site, the catalytic lid over the first redox site (pink) is in a ‘closed conformation, and BAY2402234 (yellow) occupies the space between the α2 and α3 helices that forms the ubiquinone access tunnel (cyan). FMN (orange) is shown. The surface diagrams were prepared using PyMOL. [FILM

**Extended Data Fig. 8:**
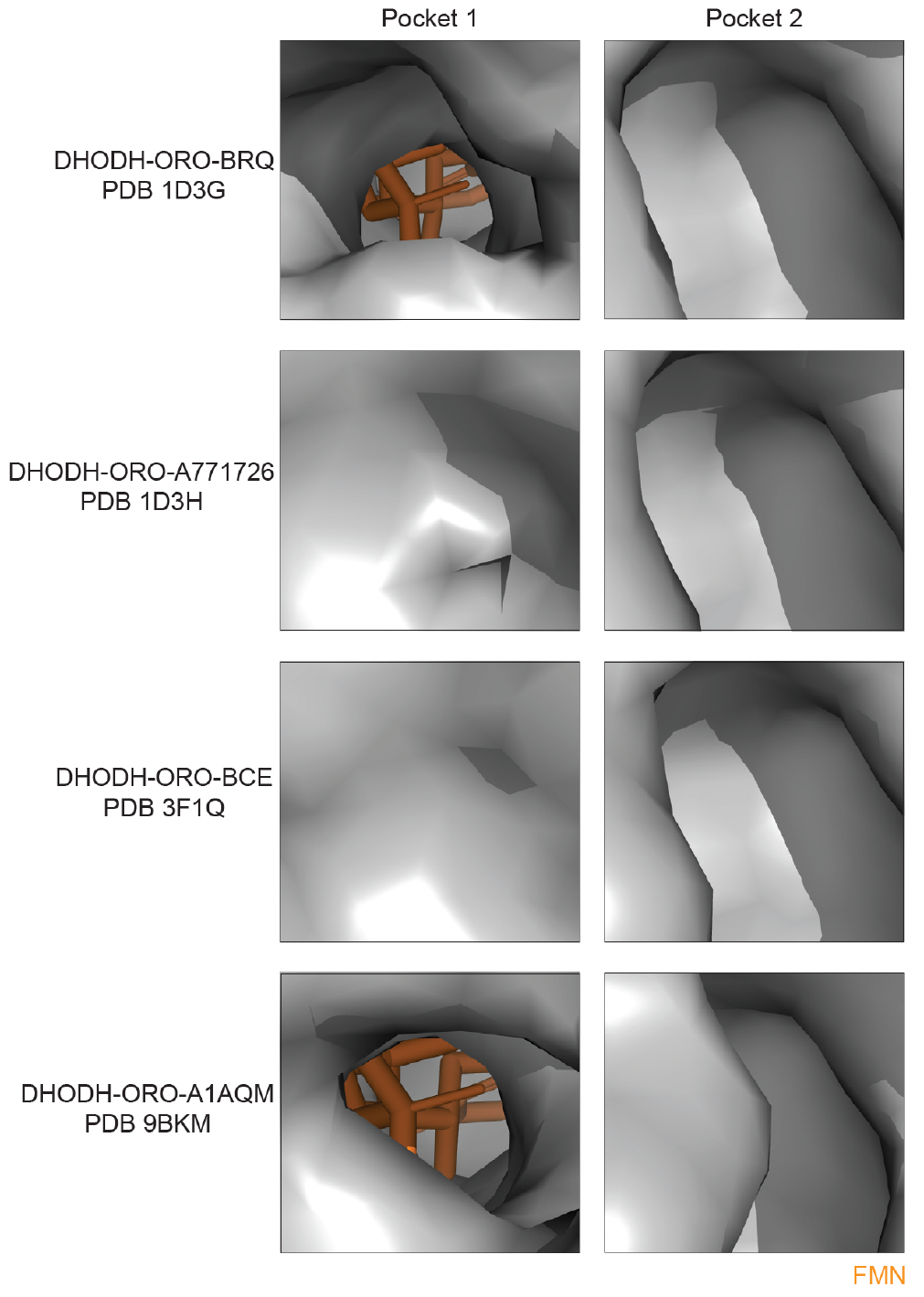
Surface views of pocket 1 and pocket 2 in DHODH-ORO-inhibitor complexes containing different clinically-relevant DHODH inhibitors. Close-up surface views of pocket 1 (left) and pocket 2 (right) in DHODH-ORO-BRQ (PDB 1D3G), DHODH-ORO-A771726/teriflunomide (PDB 1D3H), DHODH-ORO-BCE/leflunomide derivative 1 (PDB 3F1Q) and DHODH-ORO-A1AQM/JNJ-74856665 lead molecule (PDB 9BKM), as indicated. FMN (orange) is shown.

**Extended Data Fig. 9: The L2 loop over the first redox site and the α-2 helix of the ubiquinone access tunnel are highly flexible in holo-DHODH.** Film-overlay of the surface representations of DHODH-DCU and DHODH-ORO showing the movement of the lid over the first redox site (pink) and the movement of the resolved portions of the α-helices of the ubiquinone access tunnel (cyan). Pocket 2, adjacent to the ubiquinone access tunnel, remains open throughout. [FILM]

**Extended Data Fig. 10:**
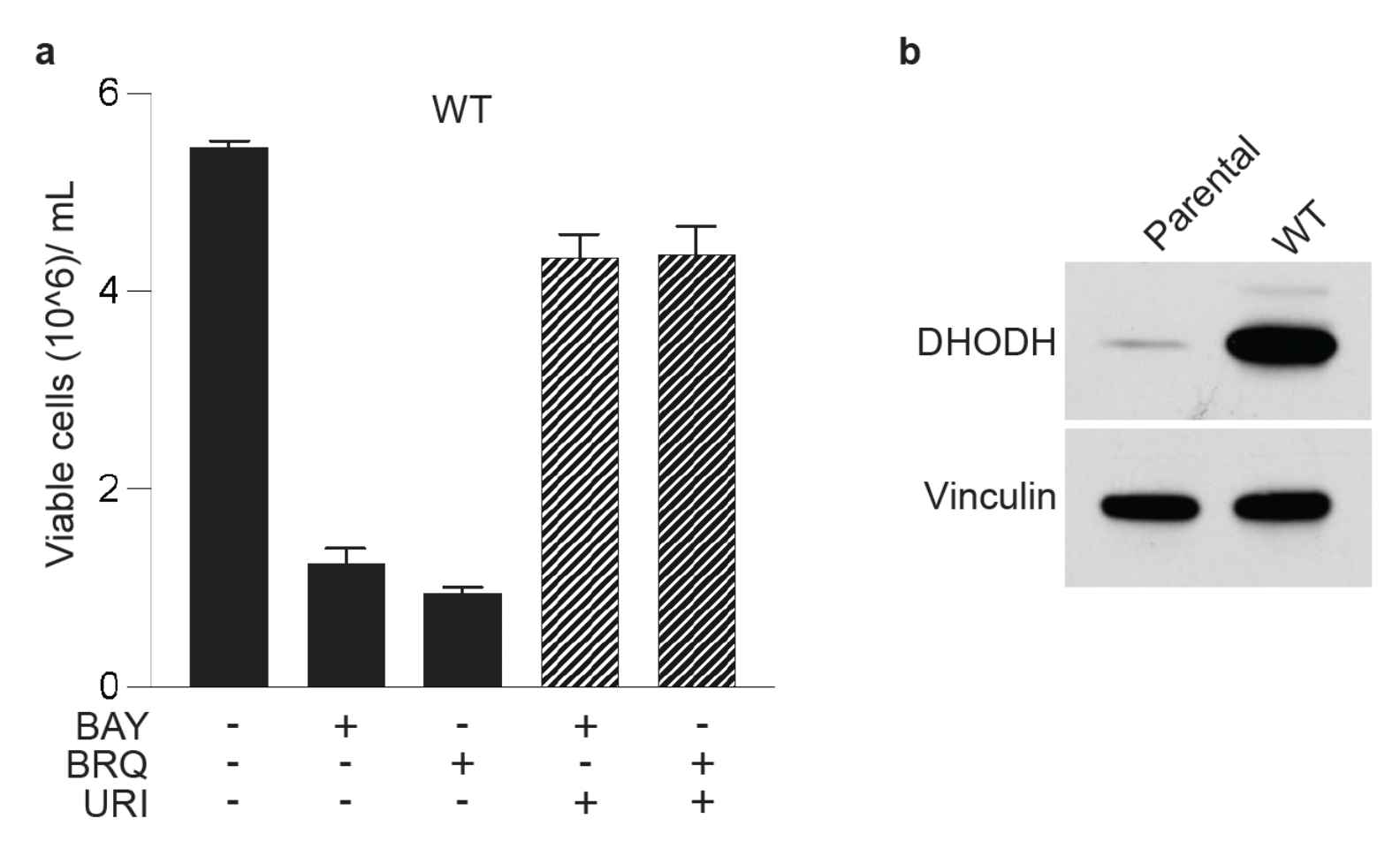
The cytotoxic effects of screening doses of BAY and BRQ act ‘on-target.’ **(a)** Number of viable TF-1 cells overexpressing DHODH^WT^ after treatment for six days with screening doses of BAY 50 nM or BRQ 5 μM in the presence or absence of uridine (URI) 50 μM, as indicated. **(b)** Immunoblot analysis of DHODH (top) and vinculin (bottom) expression in parental TF-1 cells (parental) and TF-1 cells ectopically expressing DHODH^WT^ (WT). Shown are representative results from two independent experiments.

**Extended Data Fig. 11:**
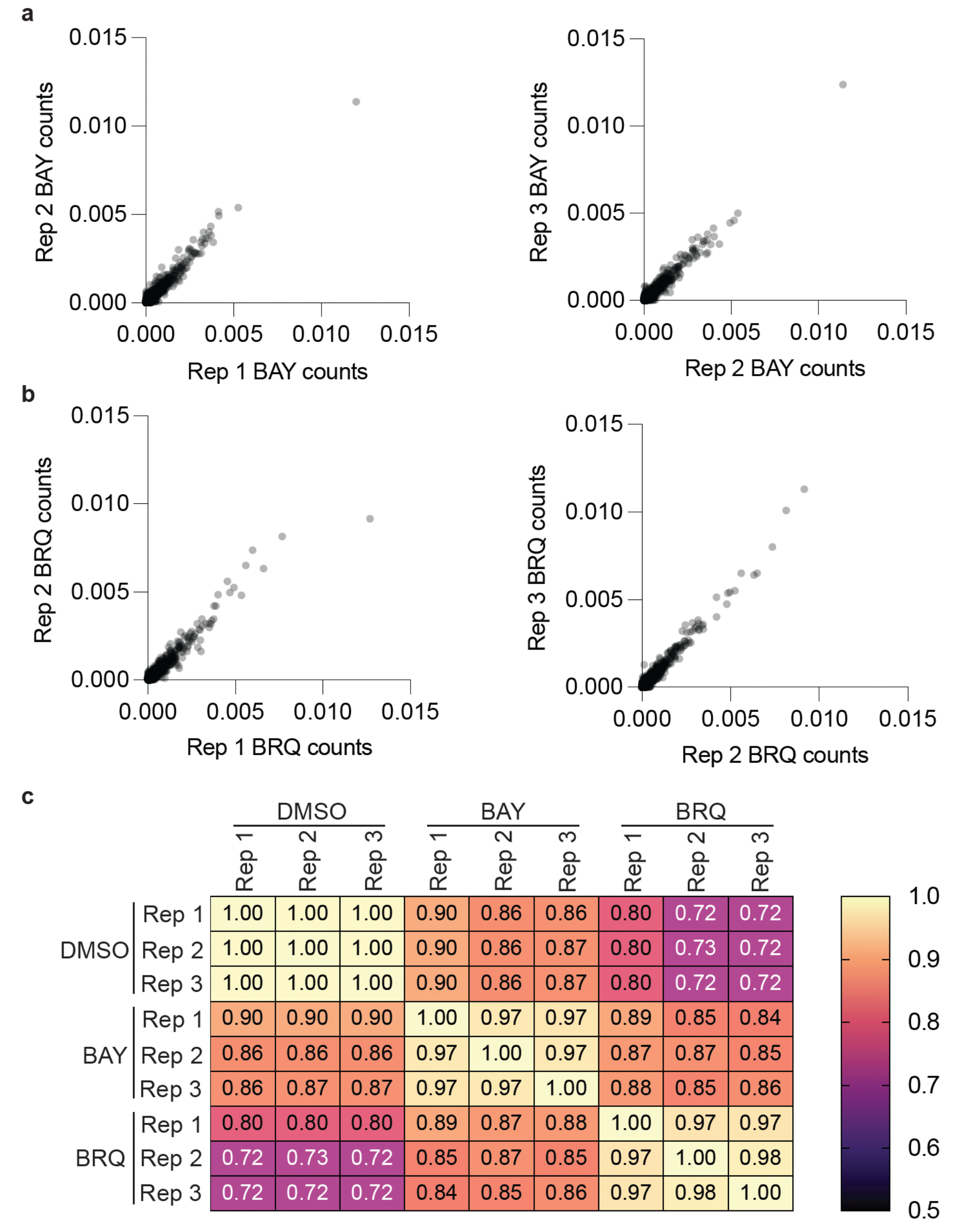
Quality control assessment of the DMS screens. **(a-b)** Scatter plots of read counts from replicate 1 vs. replicate 2 and replicate 2 vs. replicate 3 of the BAY (a) and BRQ (b) DMS drug-resistance screens. **(c)** Correlation heatmap of the results from (a) and (b).

**Extended Data Fig. 12:**
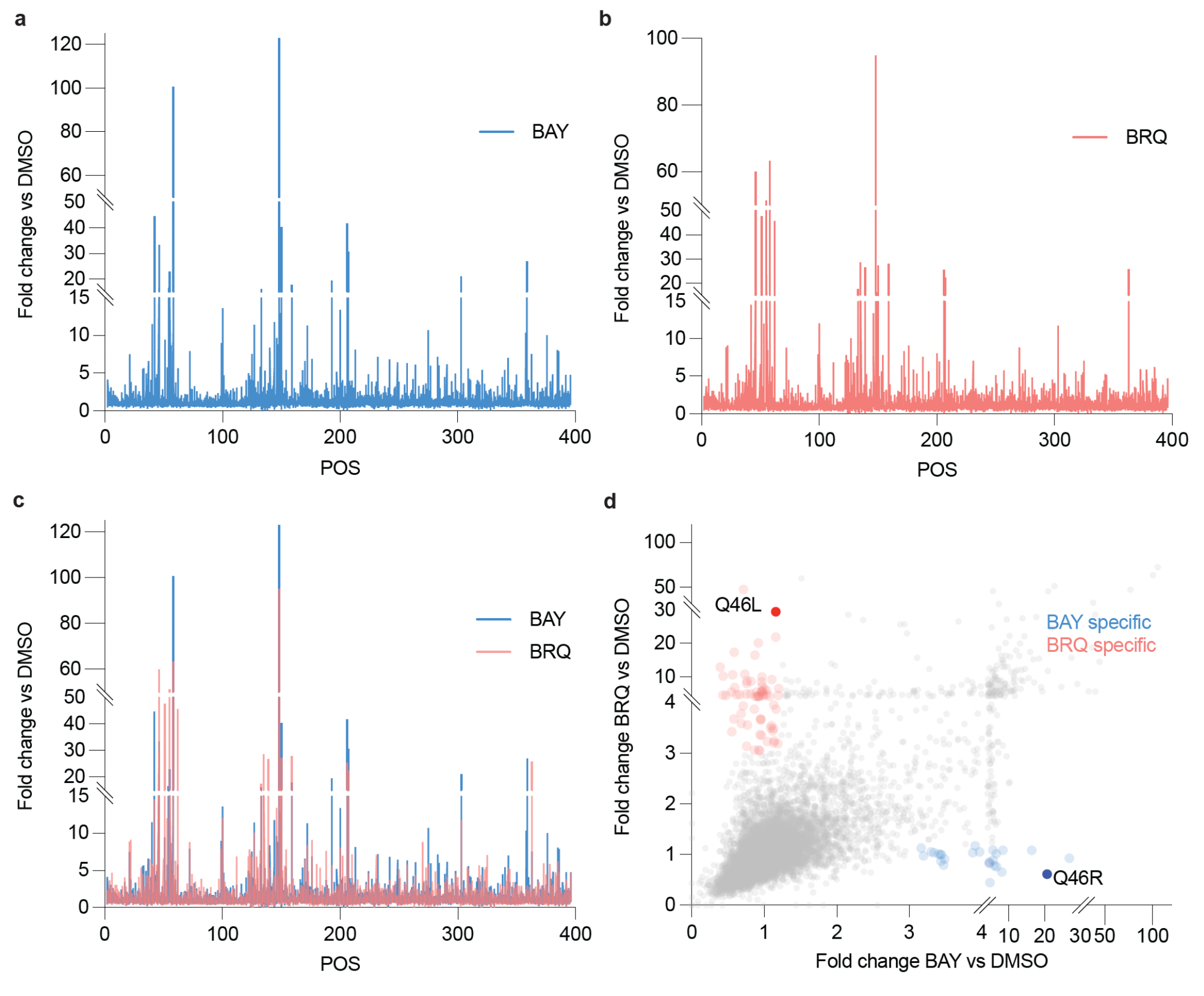
Residue mapping of amino acid substitutions in DHODH that confer resistance to BAY and/or BRQ. **(a-b)** Histogram of the combined fold change of all the substitutions at each residue in DHODH that confer resistance to BAY (a) or BRQ (b). **(c)** Overlay of the data from (a) and (b). **(d)** Scatter plot of the fold change for each variant that confers resistance to BAY (x-axis) and/or BRQ (y-axis). Mutations that specifically confer resistance to BAY (blue) or to BRQ (red) are highlighted.

**Extended Data Fig. 13:**
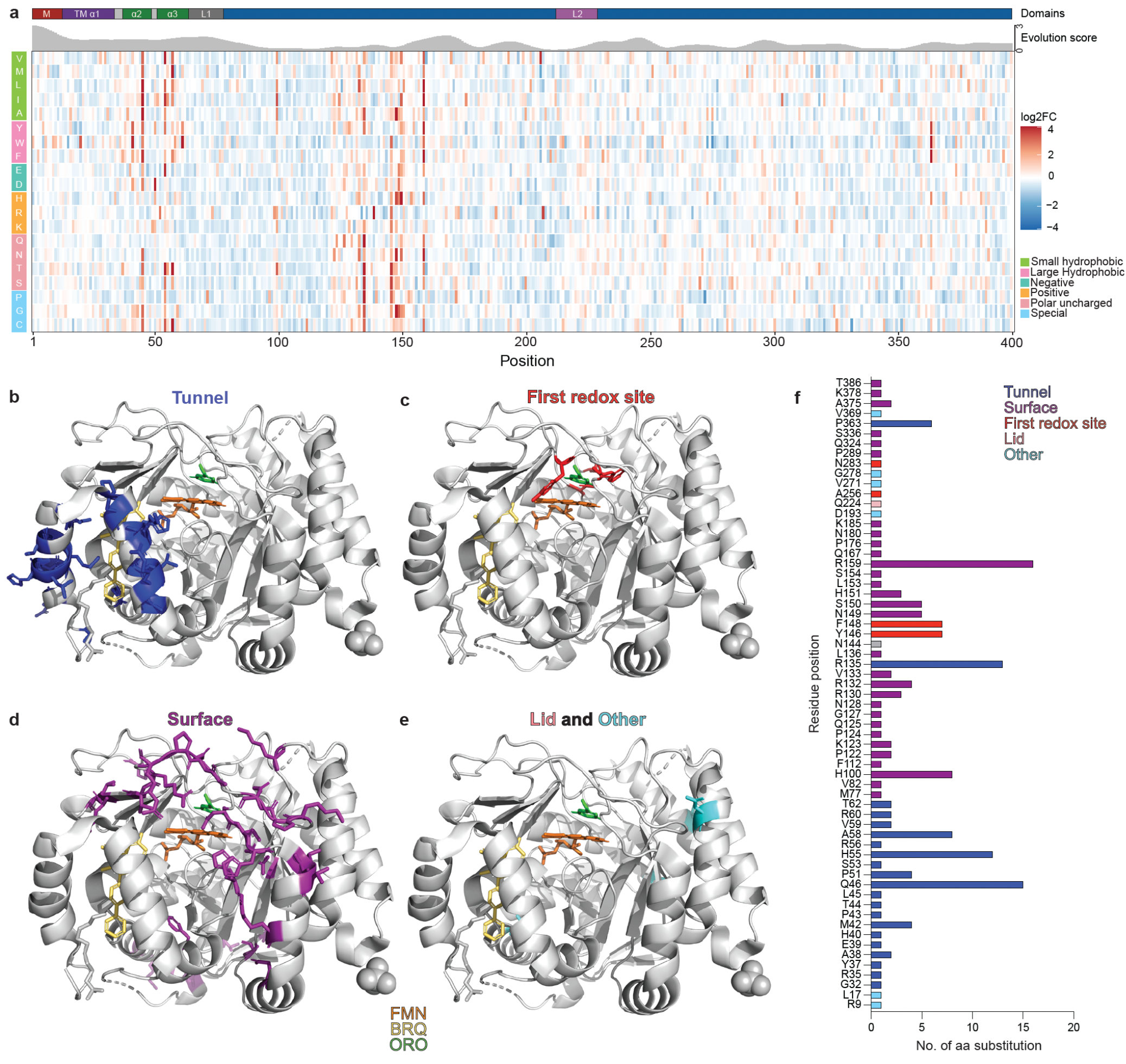
Structural mapping of BRQ-resistant DHODH variants. **(a)** Heatmap showing the log2 fold change (log2FC) in variant enrichment (red) or depletion (blue) on day 8 of the BRQ DMS screen relative to day 8 of the vehicle-control DMS screen. Each column represents an amino acid position and each row represents a substituted residue. Residues are grouped biochemically: small hydrophobic (green), large hydrophobic (pink), negatively charged (teal), positively charged (orange), polar uncharged (coral), and special (light blue). The locations in the DHODH secondary structure of the mitochondrial localization signal (M), transmembrane α1 helix (TM-α1), α2 and α3 helices, L1 loop, and L2 loop, are annotated above the heatmap. Shown below the annotation is the evolutionary score of each residue, as determined using Aminode. Lower scores indicate more conserved residues. **(b-e)** Ribbon representations of DHODH-ORO-BRQ (PDB 1D3G) highlighting the locations of mutations that confer resistance to BRQ in the ubiquinone access/drug-binding tunnel (blue) (b); first redox site (red) (c); surface (purple) (d); and lid (pink) and other locations (teal) (e). FMN (orange), BAY (yellow) and ORO (green) are highlighted. **(f)** Number of different substitutions that confer resistance to BRQ at each indicated residue position. Locations of the residues in the 3-dimensional structure of DHODH are color-coded as in (b-e). Analogous data from the BAY DMS screen are presented in **Fig. 3**. Ribbon representations were prepared using PyMOL.

**Extended Data Fig. 14:**
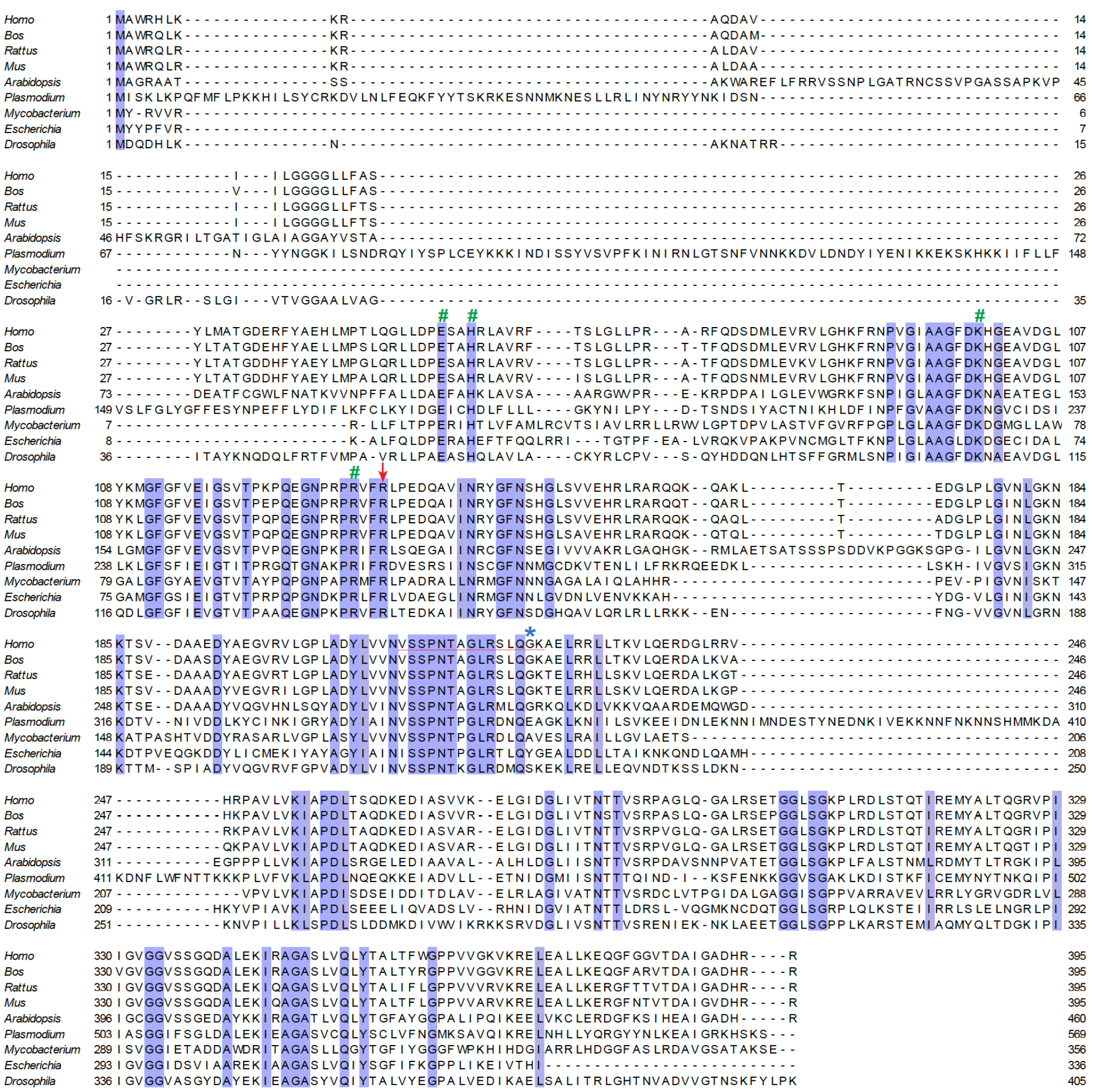
Sequence alignment of quinone-dependent DHODH orthologs. Alignment of DHODH sequences from *Homo sapiens* (*Homo*, Accession ID: Q02127), *Bos taurus* (*Bos*, Accession ID: Q5E9W3), *Rattus novergicus* (*Rattus*, Accession ID: Q63707), *Mus musculus* (*Mus*, Accession ID: O35435), *Arabidopsis thaliana* (*Arabidopsis*, Accession ID: P32746), *Plasmodium falciparum* (*Plasmodium*, Accession ID: Q08210), *Mycobacterium leprae* (*Mycobacterium*, Accession ID: P46727), *Escherichia coli* (*Escherichia*, Accession ID: P0A7E1), and *Drosophila melanogaster* (*Drosophila*, Accession ID: P32748). The residues 100% conserved across all species are shown in blue boxes. Tunnel residue R135 (red arrow), catalytic lid residues V212 to K226 (red line), pocket 2 residues E52, H55, K99, and R132 (green hashtags), and species-specific residue G225 (blue asterisk) in human DHODH are highlighted. Protein sequences were obtained from UniProt and were aligned using Clustal Omega. The alignment representation was prepared using Jalview.

**Extended Data Fig. 15:**
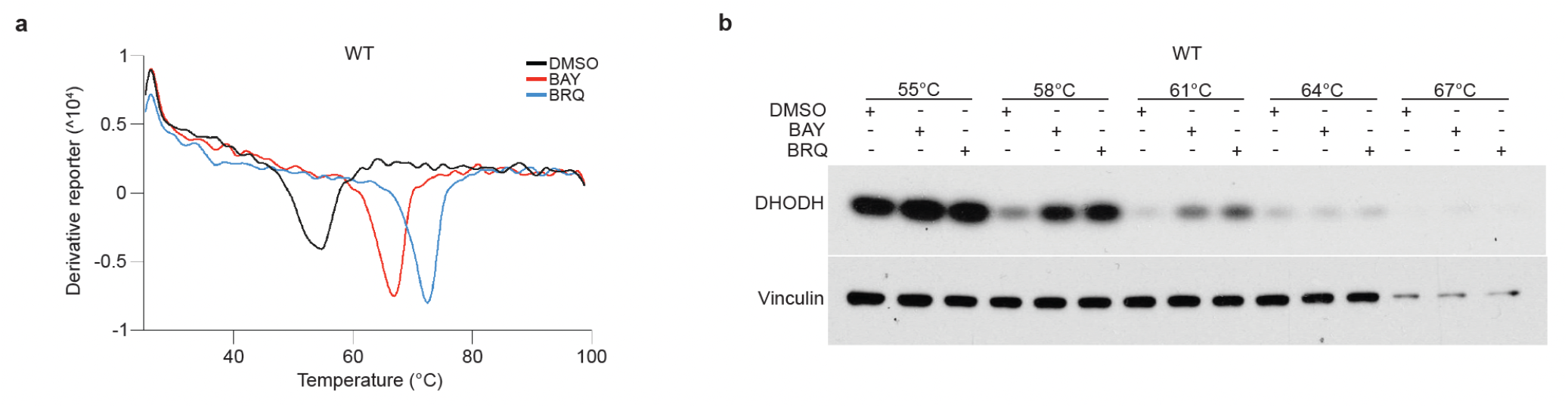
N-terminal-truncated recombinant DHODH^WT-ΔN^ and full-length DHODH^WT^ bind BAY and BRQ. **(a)** Thermal shift assay of recombinant DHODH^WT-ΔN^ treated with vehicle (DMSO; black), BAY 10 μM (red), or BRQ 10 μM (blue), as indicated. **(b)** Cellular thermal shift assay of lysates from TF-1 cells expressing DHODH^WT^ treated *in vitro* with DMSO, BAY 50 nM, or BRQ 50 nM, as indicated. Shown are representative results from three or more independent experiments.

**Extended Data Fig. 16:**
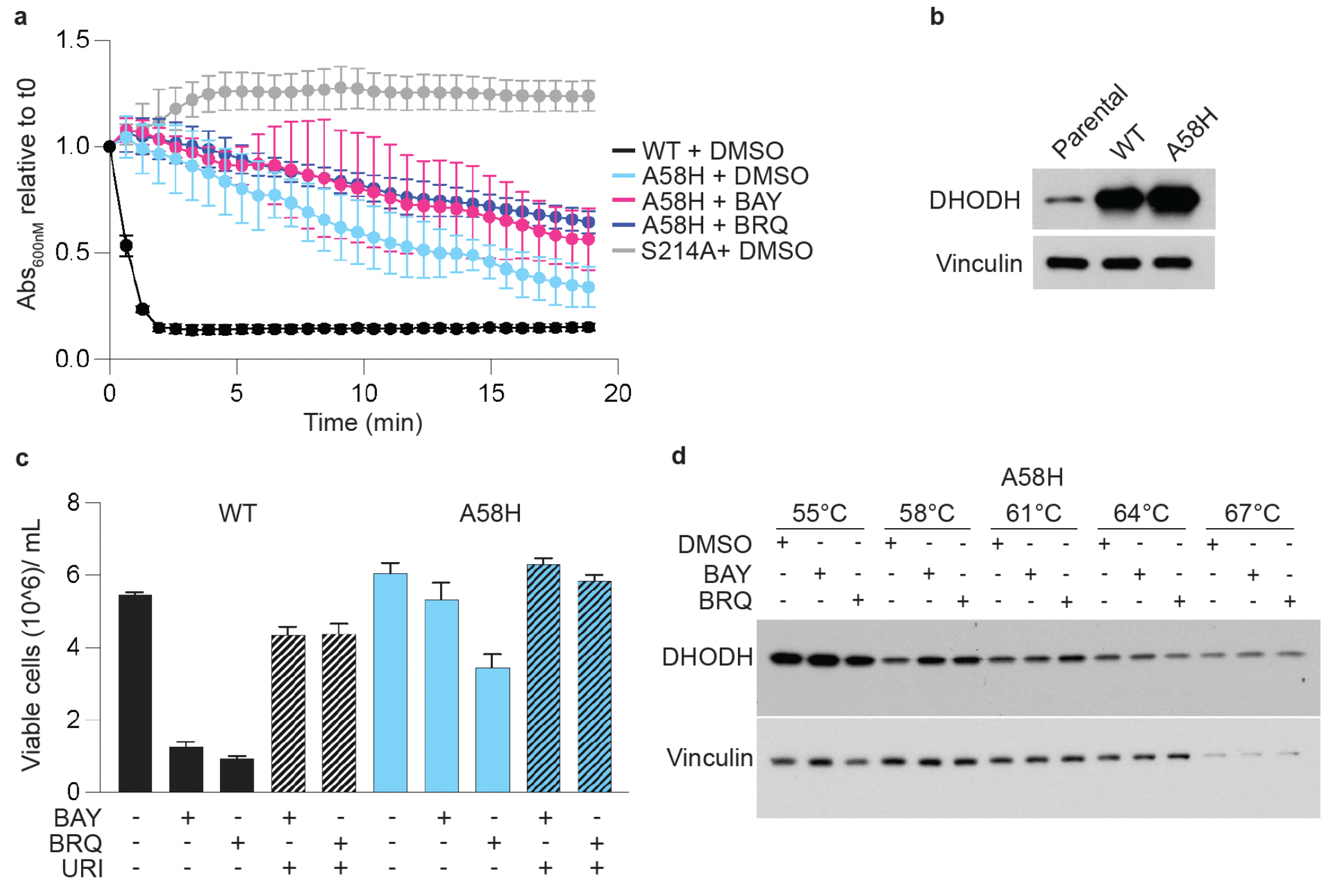
Drug-resistance mutation *DHODH^A58H^* impairs inhibitor binding. **(a)** *In vitro* catalytic activity of recombinant DHODH^WT-ΔN^, DHODH^A58H-ΔN^, and DHODH^S214A-ΔN^ in the presence of vehicle (DMSO), BAY 10 µM, or BRQ 10 µM, as indicated, as measured by DCIP assay. **(b)** Immunoblot analysis of DHODH (top) and vinculin (bottom) expression in parental TF-1 cells (Parental) and TF-1 cells expressing DHODH^WT^ (WT) or DHODH^A58H^ (A58H), as indicated. **(c)** Growth of the cells from (b) after six-day treatment with DMSO, BAY 50 nM, or BRQ 5 µM, with or without uridine (URI) 50 µM, as indicated. **(d)** CETSA of lysates from TF-1 cells expressing DHODH^A58H^ treated *in vitro* with DMSO, BAY 50 nM, or BRQ 50 nM, as indicated. Immunoblots for DHODH^A58H^ (top) and vinculin (bottom) are shown. Shown are representative results from three or more independent experiments.

**Extended Data Fig. 17:**
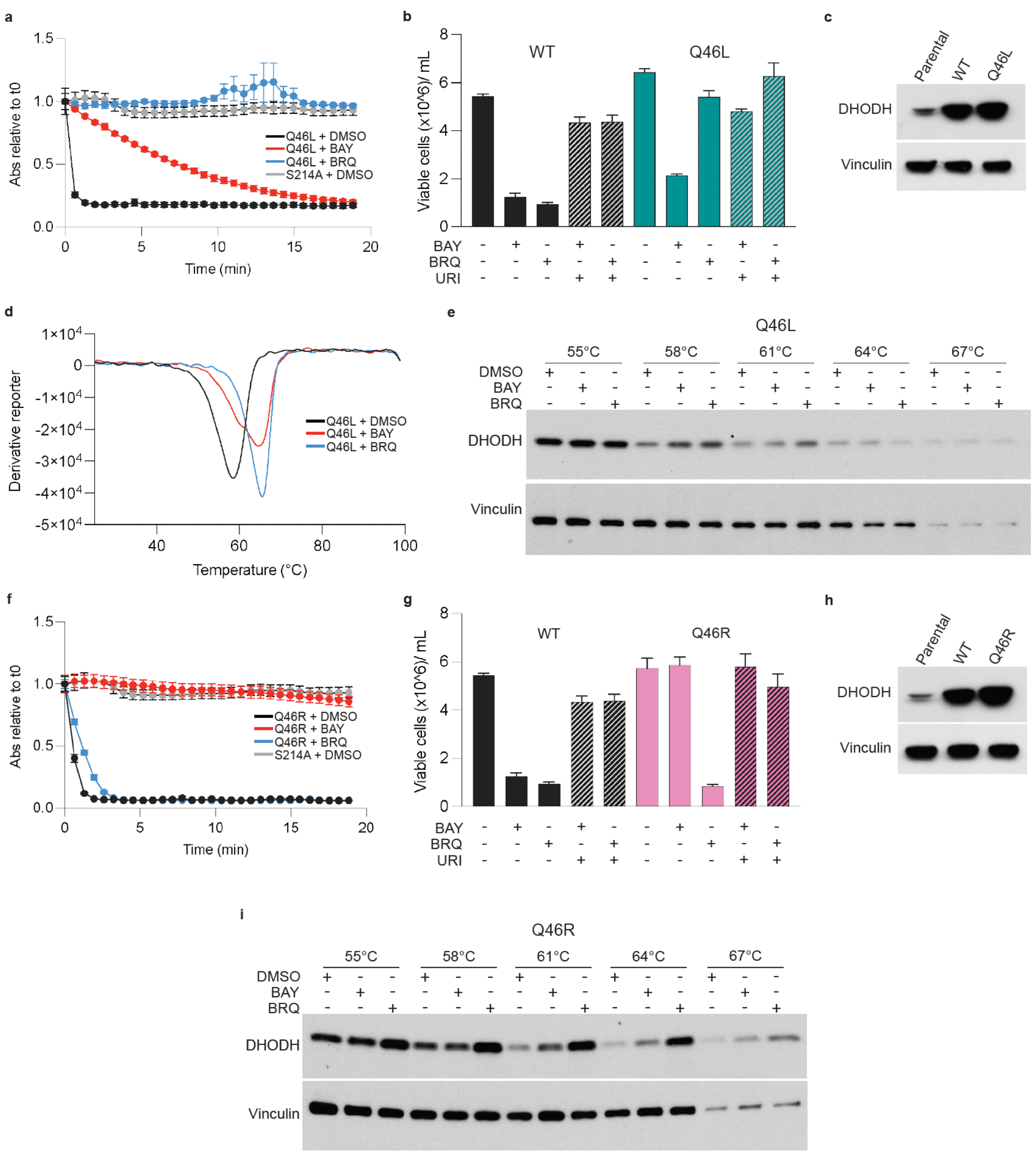
DHODH^Q46L^ and DHODH^Q46R^ bind inhibitors to which they are resistant. **(a)** *In vitro* catalytic activity of recombinant DHODH^Q46L-ΔN^ and DHODH^S214A-ΔN^ in the presence of vehicle (DMSO), BAY 10 µM, or BRQ 10 µM, as indicated, as measured by DCIP assay. **(b)** Growth of TF-1 cells expressing DHODH^WT^ (WT) or DHODH^Q46L^ (Q46L) after six-day treatment with DMSO, BAY 50 nM, or BRQ 5 µM, with or without uridine (URI) 50 µM, as indicated. **(c)** Immunoblot analysis of DHODH (top) and vinculin (bottom) expression in parental TF-1 cells (Parental) and TF-1 cells expressing DHODH^WT^ (WT) or DHODH^Q46L^ (Q46L), as indicated. (**d**) TSA of DHODH^Q46L-ΔN^ treated with DMSO (black), BAY 10 μM (red), or BRQ 10 μM (blue), as indicated. **(e)** CETSA of lysates from TF-1 cells expressing DHODH^Q46L^ treated *in vitro* with DMSO, BAY 50 nM, or BRQ 50 nM, as indicated. Immunoblots for DHODH (top) and vinculin (bottom) are shown. **(f)** *In vitro* catalytic activity of recombinant DHODH^Q46R-ΔN^ and DHODH^S214A-ΔN^ in the presence of DMSO, BAY 10 µM, or BRQ 10 µM, as indicated, as measured by DCIP assay. **(g)** Growth of TF-1 cells expressing DHODH^WT^ (WT) or DHODH^Q46R^ (Q46R) after six-day treatment with DMSO, BAY 50 nM, or BRQ 5 µM, with or without uridine (URI) 50 µM, as indicated. **(h)** Immunoblot analysis of DHODH (top) and vinculin (bottom) expression in parental TF-1 cells (Parental) and TF-1 cells expressing DHODH^WT^ (WT) or DHODH^Q46R^ (Q46R), as indicated. **(i)** CETSA of lysates from TF-1 cells expressing DHODH^Q46R^ treated *in vitro* with DMSO, BAY 50 nM, or BRQ 50 nM, as indicated. Immunoblots for DHODH (top) and vinculin (bottom) are shown. Shown are representative results from three or more independent experiments.

**Extended Data Fig. 18:**
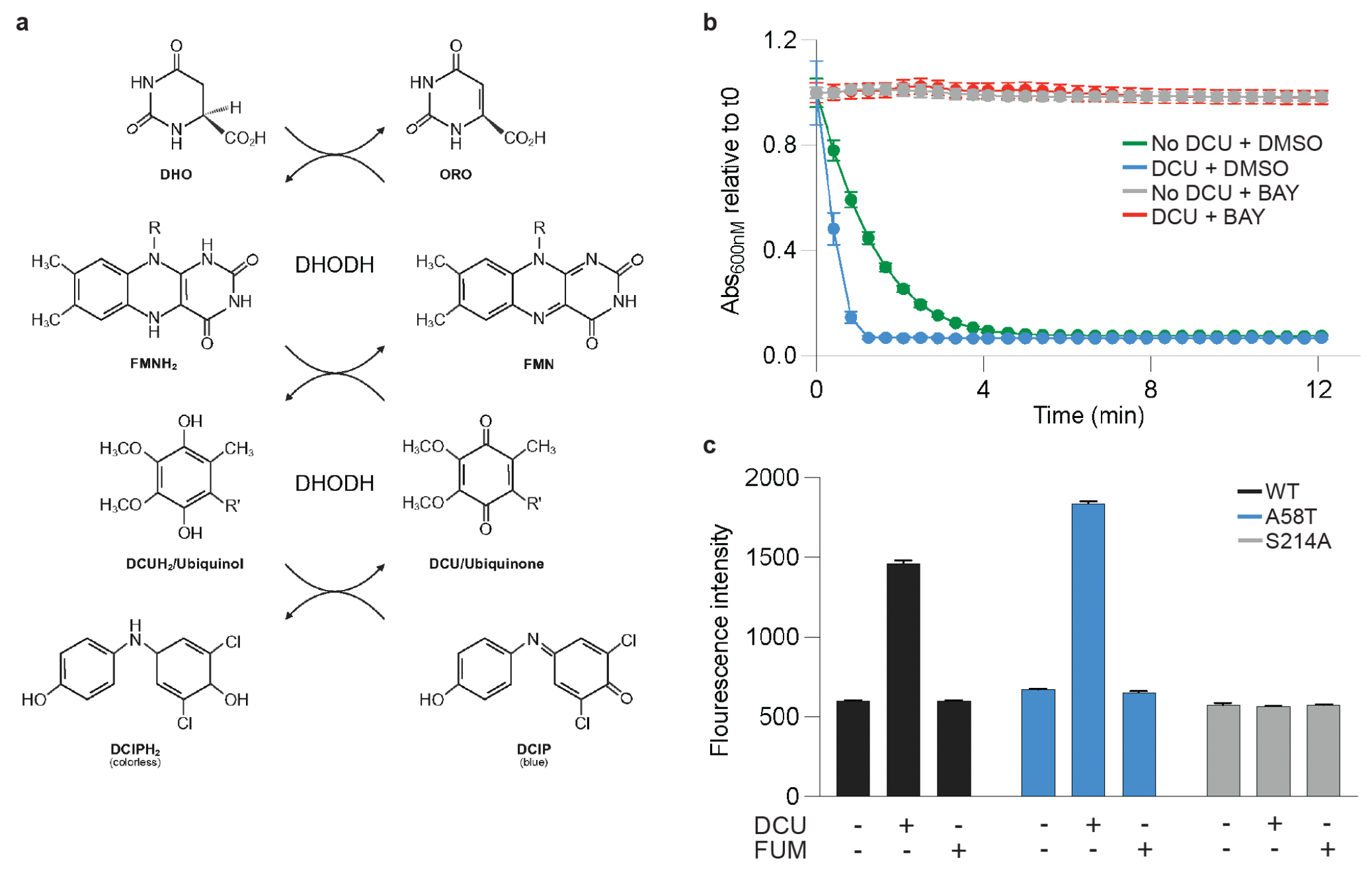
Recombinant human DHODH cannot use fumarate or molecular oxygen as alternative terminal electron acceptors. **(a)** Schematic of the DCIP absorbance assay, prepared using BioRender. (b) *In vitro* catalytic activity of recombinant DHODH^WT-ΔN^ in the presence of vehicle (DMSO; green), DCU 100 µM (blue), BAY 100 µM (grey), or DCU 100 µM + BAY 100 µM (red) as indicated, as measured by DCIP assay. (c) *In vitro* catalytic activity of recombinant DHODH^WT-ΔN^ (black), DHODH^A58T-ΔN^ (blue), or DHODH^S214A-ΔN^ (gray) in the presence of DMSO (-), DCU 500 µM (+), or fumarate 500 µM (+) as indicated, as measured by OFA. Shown are representative results from three or more independent experiments.

**Extended Data Fig. 19:**
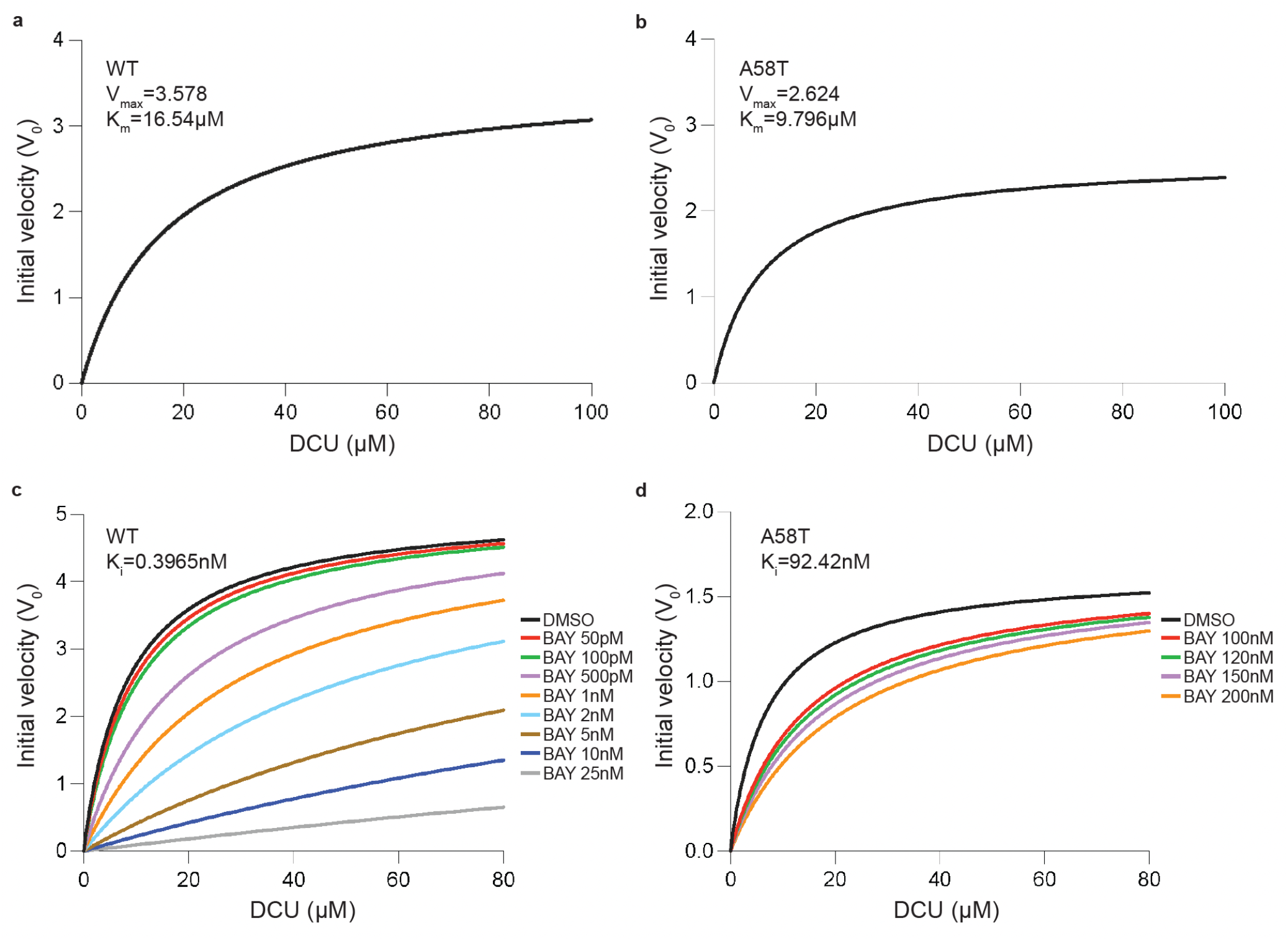
Enzyme kinetics analysis of DHODH^WT^ and DHODH^A58T^. **(a-b)** Michaelis-Menten kinetics at a saturating DHO concentration (500 µM) and a range of concentrations of DCU for DHODH^WT-ΔN^ (a) and DHODH^A58T-ΔN^ (b). **(c-d)** Michaelis-Menten kinetics at a saturating DHO concentration (500 µM) and a range of concentrations of DCU and BAY for DHODH^WT-ΔN^ (c) and DHODH^A58T-ΔN^ (d). At limiting concentrations of DCU, BAY acts as competitive inhibitor of both variants. The analyses were performed using the GraphPad Prism competitive inhibition equation. Shown are representative results from three or more independent experiments.

**Extended Data Fig. 20:**
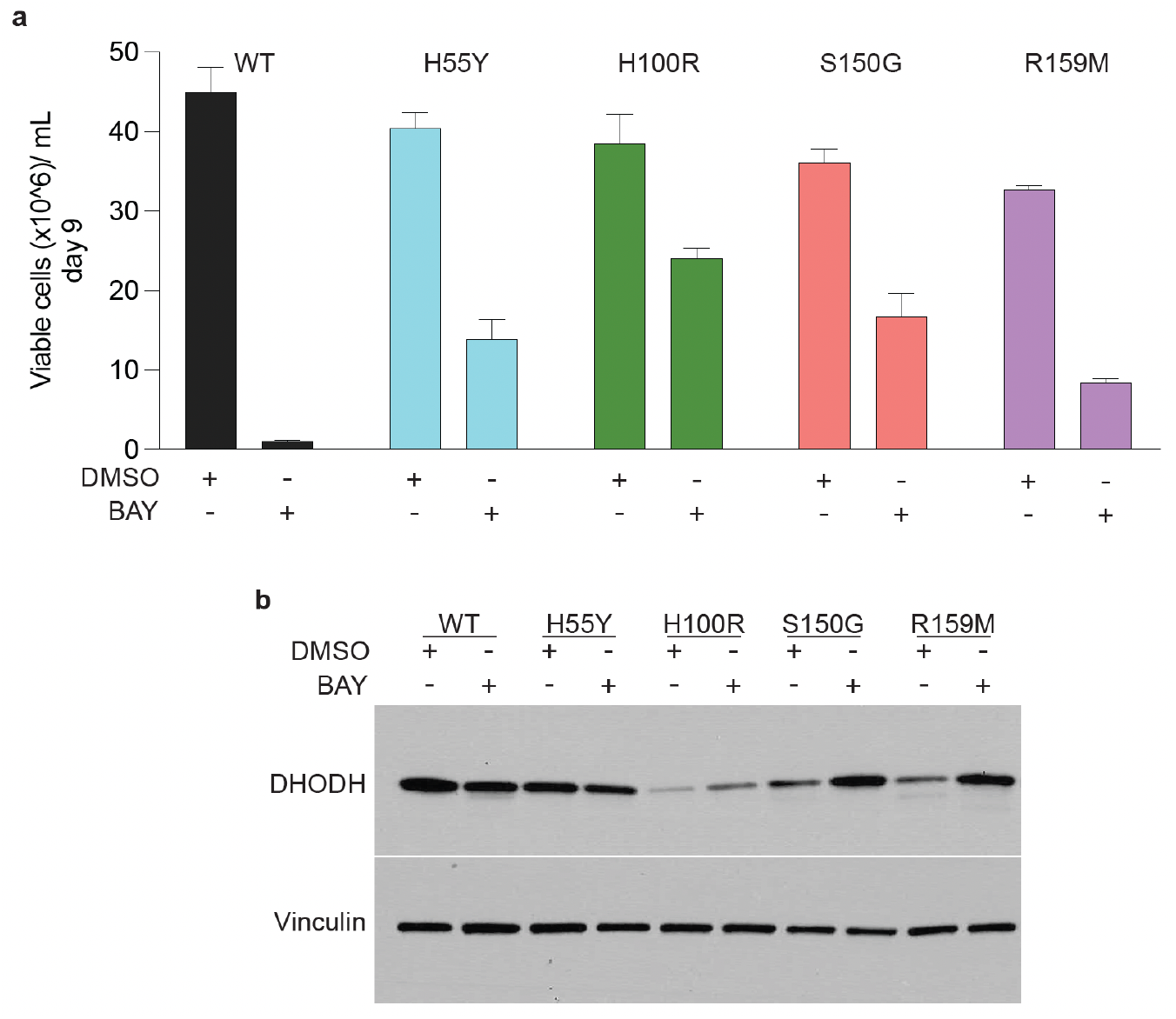
Several drug-resistant DHODH variants with substitutions located near pocket 2 are poorly expressed in cells and are upregulated by inhibitor treatment. **(a)** Growth of TF-1 cells expressing DHODH^WT^ (WT), DHODH^H55Y^ (H55Y), DHODH^H100R^ (H100R), DHODH^S150G^ (S150G), or DHODH^R159M^ (R159M) after nine-day treatment with vehicle (DMSO) or BAY 50 nM, as indicated. **(b)** Immunoblot analysis of DHODH (top) and vinculin (bottom) expression in the cells from (a) following overnight treatment with DMSO or BAY 50 nM, as indicated. Shown are representative results from three or more independent experiments.

**Extended Data Fig. 21:**
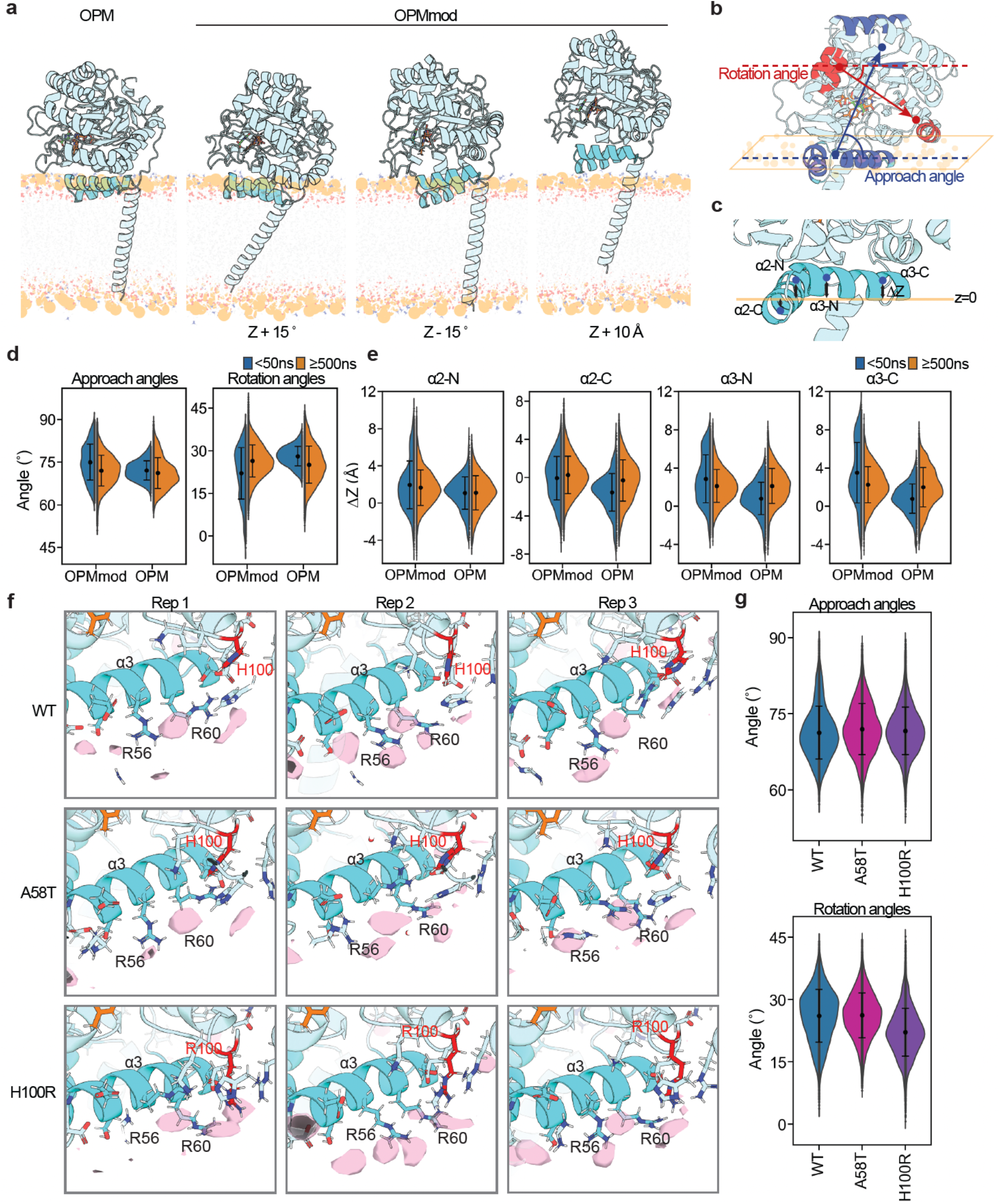
MD simulation of DHODH-membrane interactions. **(a)** Initial MD simulation systems with varied orientations. The ‘OPM’ geometry was generated using PPM 3.0 web server. The structure of full-length DHODH^WT^ was constructed from our apo-state crystal structure (DHODH-ORO), complemented with the loop region from our DHODH-DCU structure and the transmembrane span from the AlphaFold3 model. Modified ‘OPMmod’ geometries were generated by rotating DHODH by 15° about the z-axis perpendicular to the membrane plane in both directions (Z + 15° and Z - 15°) or by translating DHODH 10 Å upward along the z-axis (Z + 10 Å). **(b)** Schematic cartoon of the geometry of DHODH relative to membrane. ‘Approach angle’ is defined as the angle between the membrane plane of the upper leaflet (orange parallelogram) and the vector connecting the center of the membrane-embedding helices (α2: residues 35–47; α3: residues 52–63) to the center of β10 (residues 251–254) and α11 (residues 262–273), highlighted in blue. ‘Rotation angle’ is defined as the angle between the upper leaflet membrane plane and the vector connecting the center of α12 (residues 339–346) and α13 (residues 310–315) to the center of α6 (residues 154–161) and β7 (residues 177–179), highlighted in red. **(c)** Schematic cartoon of membrane embedding. For each membrane-embedding helix, N-terminal and C-terminal segments were selected (α2-N: residues 35-39; α2-C: residues 43-47; α3-N: residues 53-57; α3-C: residues 59-63). **(d)** Violin plots showing the distribution of approach and rotation angles, as defined in (B), for DHODH^WT^. ‘OPMmod’ depicts the results of simulations using the three modified geometries shown in (a); ‘OPM’ depicts the results of simulations using the OPM geometry shown in (a). 1-µs simulations were performed in triplicate for each initial geometry. The blue plots on the left show the distribution for the first 50 ns, and the orange plots on the right show the distribution for the last 500 ns. Black dots represent averages and error bars represent standard deviations. **(e)** Violin plots showing the distribution of membrane-embedding depths for each N-terminal and C-terminal segment of the membrane-embedding helices, as defined in (c). The z-height to the center of each segment from the membrane plane of the upper leaflet was measured. Plots are presented in the same manner as in (d). **(f)** Snapshots showing the volumetric density of POPE phosphates (pink) in each replicate of the 1-µs simulations for DHODH^WT^ (top), DHODH^A58T^ (middle), and DHODH^H100R^ (bottom). FMN (orange), membrane-embedded helices (cyan), and H100 or R100 (red) are indicated. **(g)** Violin plots showing the distributions of approach (top) and rotation (bottom) angles, as defined in (d), for DHODH^WT^ (blue), DHODH^A58T^ (magenta), and DHODH^H100R^ (purple). Data from 1-µs simulations performed in triplicate for each variant are shown. Black dots represent averages and error bars represent standard deviations.

**Extended Data Fig. 22:**
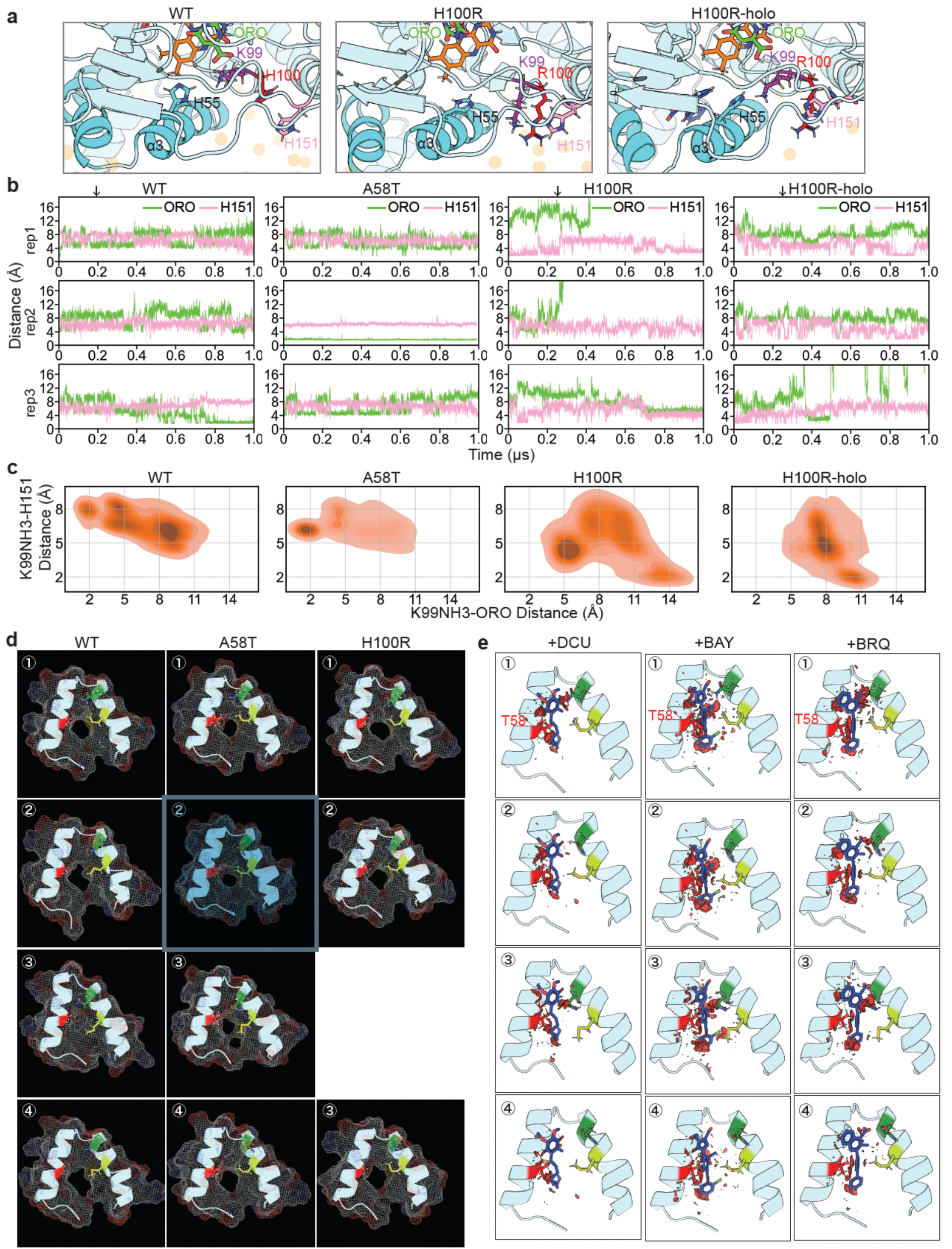
MD simulation of the effects of drug-resistance mutations on ligand-binding. **(a)** Representative snapshots from MD simulations of inhibitor-free DHODH^WT^ (WT), inhibitor-free DHODH^H100R^ (H100R), and DHODHi-bound DHODH^H100R^ (H100R-inh), showing how the H100R mutation may influence the local interaction network and ORO-binding. POPE phosphates (tan spheres), FMN (orange), ORO (green), membrane-embedding helices (cyan), the mutated H/R-100 site (red), and key surrounding residues K99 (purple) and H151 (pink) are indicated. **(b)** Line plots showing the minimum distances between the ammonium group of K99 and either ORO (green) or H151 (pink) over time for each replicate. The black arrows indicate the frames at which snapshots shown in (a) were taken. **(c)** Contour plots showing the distribution of distances between the ammonium group of K99 and H151 (K99NH3-H151) and between K99NH3 and ORO (K99NH3-ORO) for each variant. Frames collected after ORO dissociation (defined as K99NH3-ORO distances > 17 Å) were excluded from the analysis. **(d)** Representative snapshots of the unique tunnel conformations outlined by orange boxes in Figure 7 (f). Residues A/T-58 (red), M42 (light green) and Q46 (dark green) are indicated. Protein surfaces are shown as meshes. **(e)** Representative snapshots of tunnel structures from MD simulations of DHODH^A58T^ bound to DCU, BAY, or BRQ, as indicated. The ligands were positioned in the tunnel by aligning the tunnel structures with our DHODH-DCU structure (+DCU); PDB 6QU7 (+BAY), or PDB 1D3G (+BRQ). Potential clashes with ligands are indicated by red discs.

**Extended Data Fig. 23:**
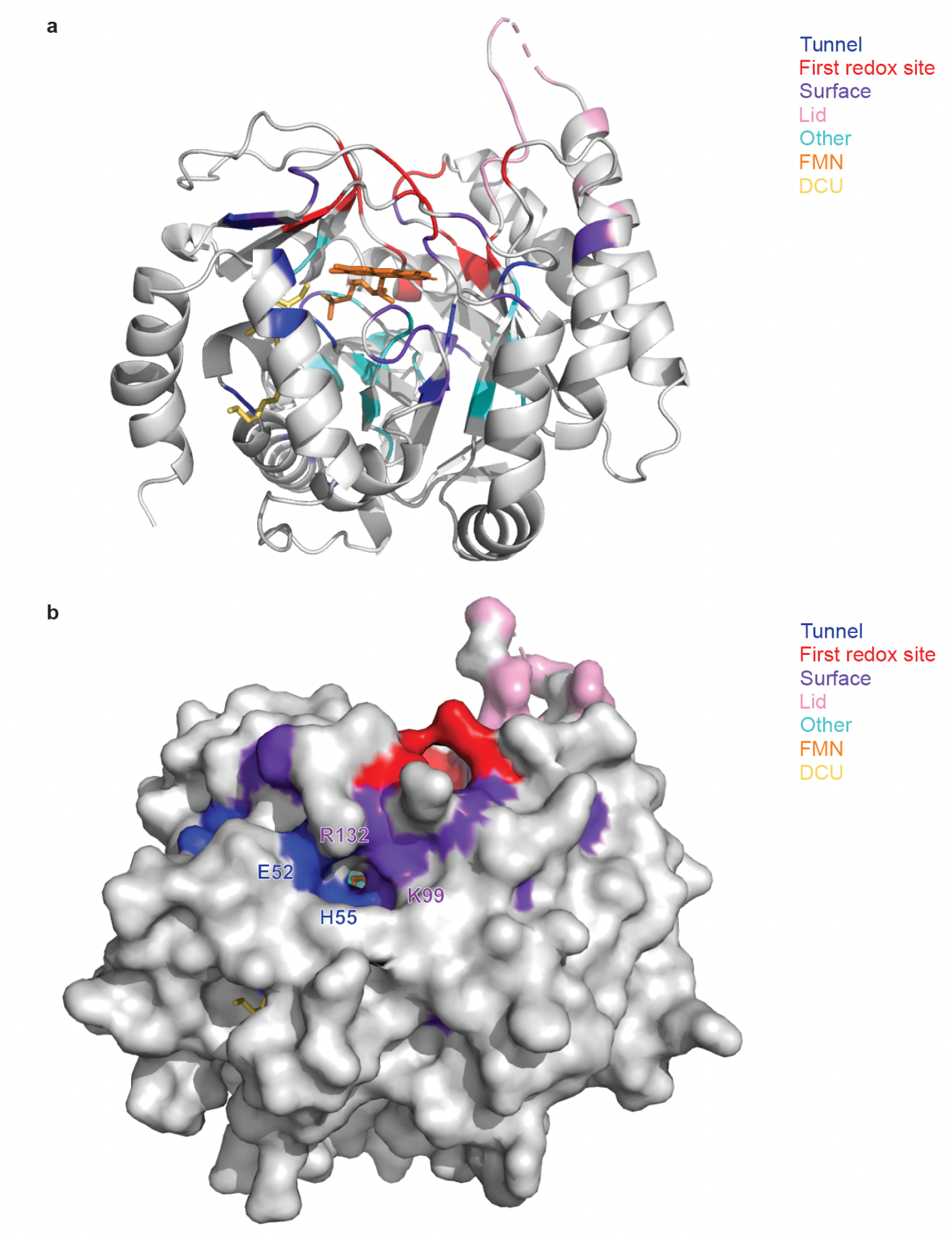
Evolutionarily conserved residues in class 2 DHODH orthologs. **(a-b)** Ribbon representation (a) and surface representation (b) of DHODH-DCU with 100% conserved residues from **Extended Data Figure 14** highlighted. Fully conserved residues in the region around pocket 2 are annotated in (b). Residues are color-coded based on their locations in DHODH: ubiquinone access/drug-binding tunnel (blue), first redox site (red), surface (purple), lid (pink), and other locations (teal). The structural representations were prepared using PyMOL.

**Extended Data Table 1:** X-ray crystallography data collection and refinement statistics. Metrics from x-ray crystal structure data collection and processing are indicated. Values in parentheses are for the highest-resolution shell.

|  | **DHODH-DCU** | **DHODH-ORO** |
| --- | --- | --- |
| **Wavelength** | 0.9793 | 0.9201 |
| **Resolution range** | 47.48-2.21 (2.24-2.21) | 51.24 -3.01 (3.2 -3.01) |
| **Space group** | P 1 21 1 | P 63 |
| **Unit cell** | 48.25 181.23 119.91 90 103.06 90 | 126.74 126.74 87.09 90 90 120 |
| **Total reflections** | 1578411 (12727) | 207101 (35320) |
| **Unique reflections** | 99742 (3271) | 15882 (2618) |
| **Multiplicity** | 15.8 (3.9) | 13.0 (13.5) |
| **Completeness (%)** | 99.59 (99.82) | 99.71 (99.24) |
| **Mean I/sigma(I)** | 4.9 (1.69) | 9.41 (0.66) |
| **Wilson B-factor** | 19.59 | 99.32 |
| **R-merge** | 0.5255 (0.7845) | 0.2178 (4.906) |
| **R-meas** | 0.5424 (0.9084) | 0.2267 (5.098) |
| **R-pim** | 0.1309 (0.4528) | 0.06257 (1.379) |
| **CC1/2** | 0.98 (0.337) | 0.999 (0.295) |
| **CC\*** | 0.995 (0.71) | 1 (0.675) |
| **Reflections used in refinement** | 99548 (3269) | 15858 (2599) |
| **Reflections used for R-free** | 4865 (145) | 784 (128) |
| **R-work** | 0.2108 (0.3358) | 0.2136 (0.3376) |
| **R-free** | 0.2488 (0.3602) | 0.2633 (0.3507) |
| **Number of non-hydrogen atoms** | 12228 | 5351 |
| **Macromolecules** | 10944 | 5266 |
| **Ligands** | 235 | 84 |
| **Solvent** | 1049 | 1 |
| **Protein residues** | 1440 | 691 |
| **RMS(bonds)** | 0.005 | 0.001 |
| **RMS(angles)** | 0.66 | 0.37 |
| **Ramachandran favored (%)** | 98.11 | 97.64 |
| **Ramachandran allowed (%)** | 1.89 | 2.36 |
| **Ramachandran outliers (%)** | 0 | 0 |
| **Rotamers outliers (%)** | 0.96 | 0 |
| **Clashscore** | 21.63 | 7.65 |
| **Average B-factor** | 30.74 | 132.64 |
| **Macromolecules** | 30.4 | 133.18 |
| **Ligands** | 29.03 | 99.28 |
| **Solvent** | 34.7 | 76.29 |

**Extended Data Table 2:** Protein-ligand interaction profiler output. Indicated are distance parameters of non-covalent interactions between DCU, BAY, or BRQ, as indicated, with specific side-chains in the ubiquinone access tunnel of DHODH, derived from the experimentally determined x-ray crystal structures using protein-ligand interaction profiler (PLIP).

| DHODH-DCU |  | DHODH-ORO-BAY<br>(PDB 6QU7) |  | DHODH-ORO-BRQ<br>(PDB 1D3G) |  |
| --- | --- | --- | --- | --- | --- |
| Residue | Distance (Å) | Residue | Distance (Å) | Residue | Distance (Å) |
| **Hydrophobic interactions** |  | **Hydrophobic interactions** |  | **Hydrophobic interactions** |  |
| A54 | 3.76 | Y37 | 3.99 | L45 | 3.58 |
| F61 | 3.6 | L45 | 3.89 | E46 | 3.9 |
| F97 | 3.93 | A54 | 3.88 | P51 | 3.89 |
| F97 | 3.85 | A58 | 3.28 | A54 | 3.8 |
| L358 | 3.82 | F61 | 3.74 | H55 | 3.47 |
| P363 | 3.82 | L67 | 3.52 | A58 | 3.55 |
| **Hydrogen bonds** |  | V133 | 3.95 | L67 | 3.48 |
| Y355 | 2.3 | L358 | 3.79 | Y355 | 3.79 |
|  |  | T359 | 3.89 | T359 | 3.55 |
|  |  | P363 | 3.94 | P363 | 3.72 |
|  |  | **Hydrogen bonds** |  | **Hydrogen bonds** |  |
|  |  | E46 | 2.12 | E46 | 2.13 |
|  |  | Y355 | 1.96 | P51 | 3.54 |
|  |  | **Halogen bonds** |  | **Water bridges** |  |
|  |  | A58 | 3.28 | T359 | 2.63 |
|  |  |  |  | **Salt bridges** |  |
|  |  |  |  | R135 | 3.46 |

**Extended Data Table 3:**
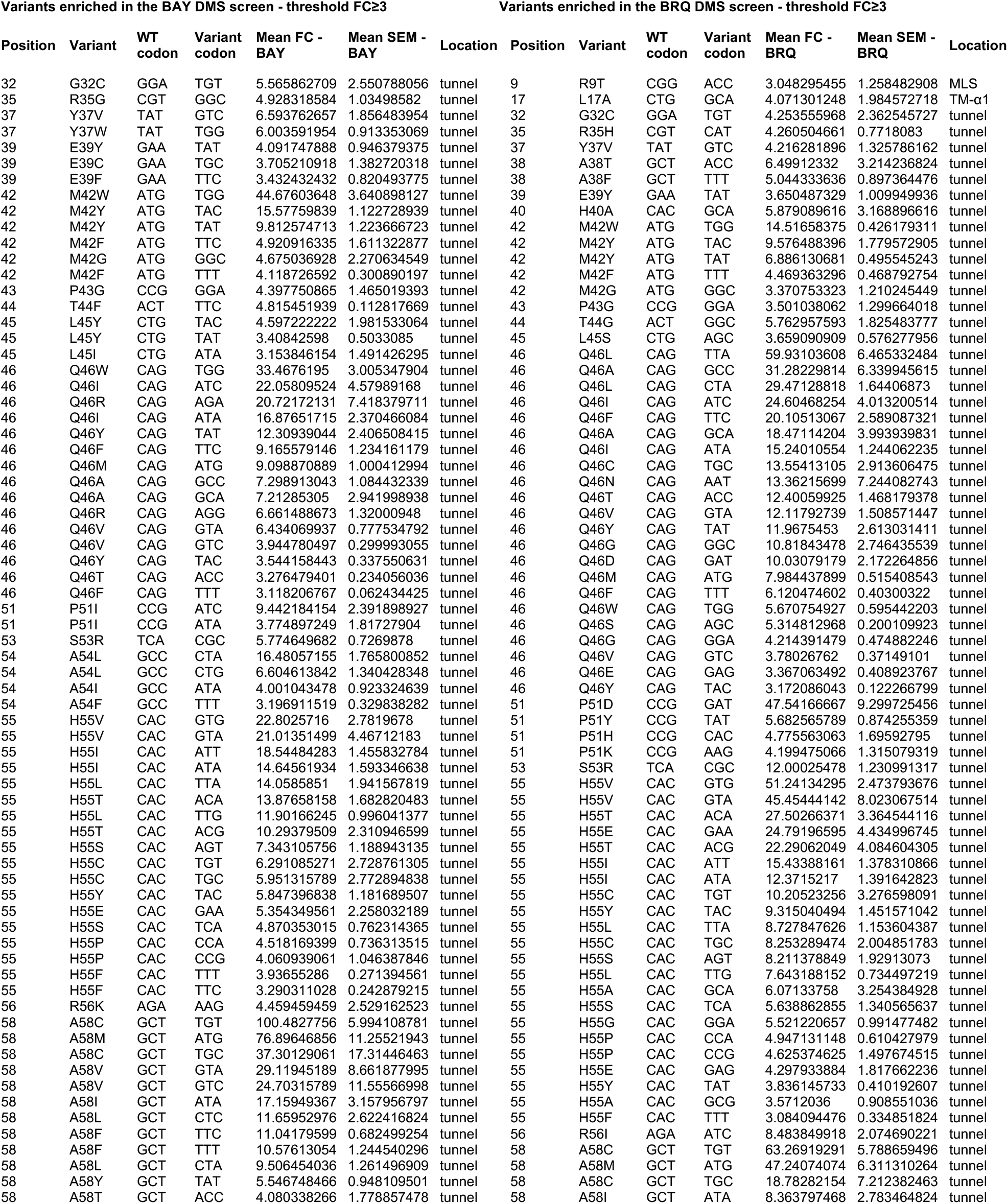
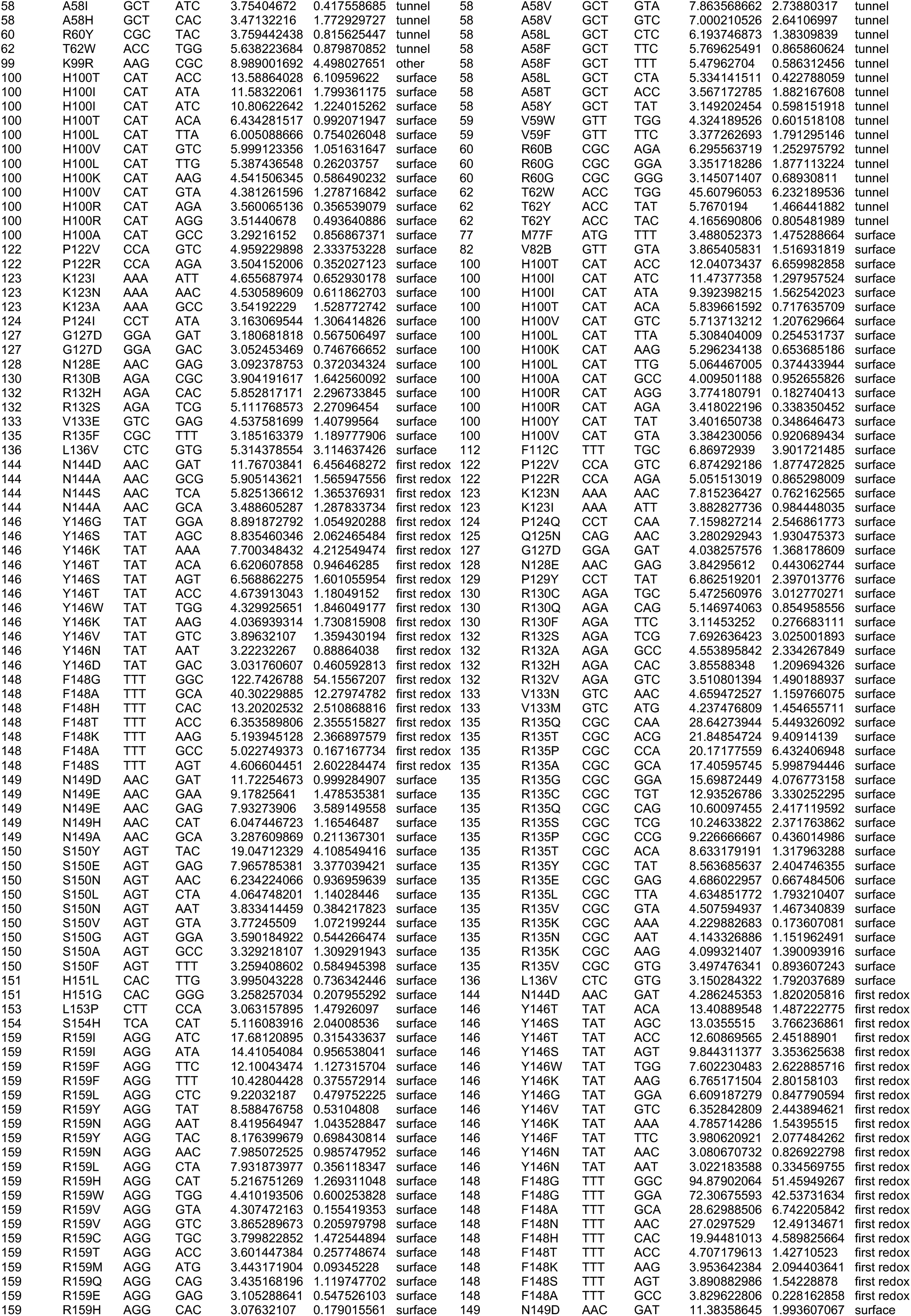
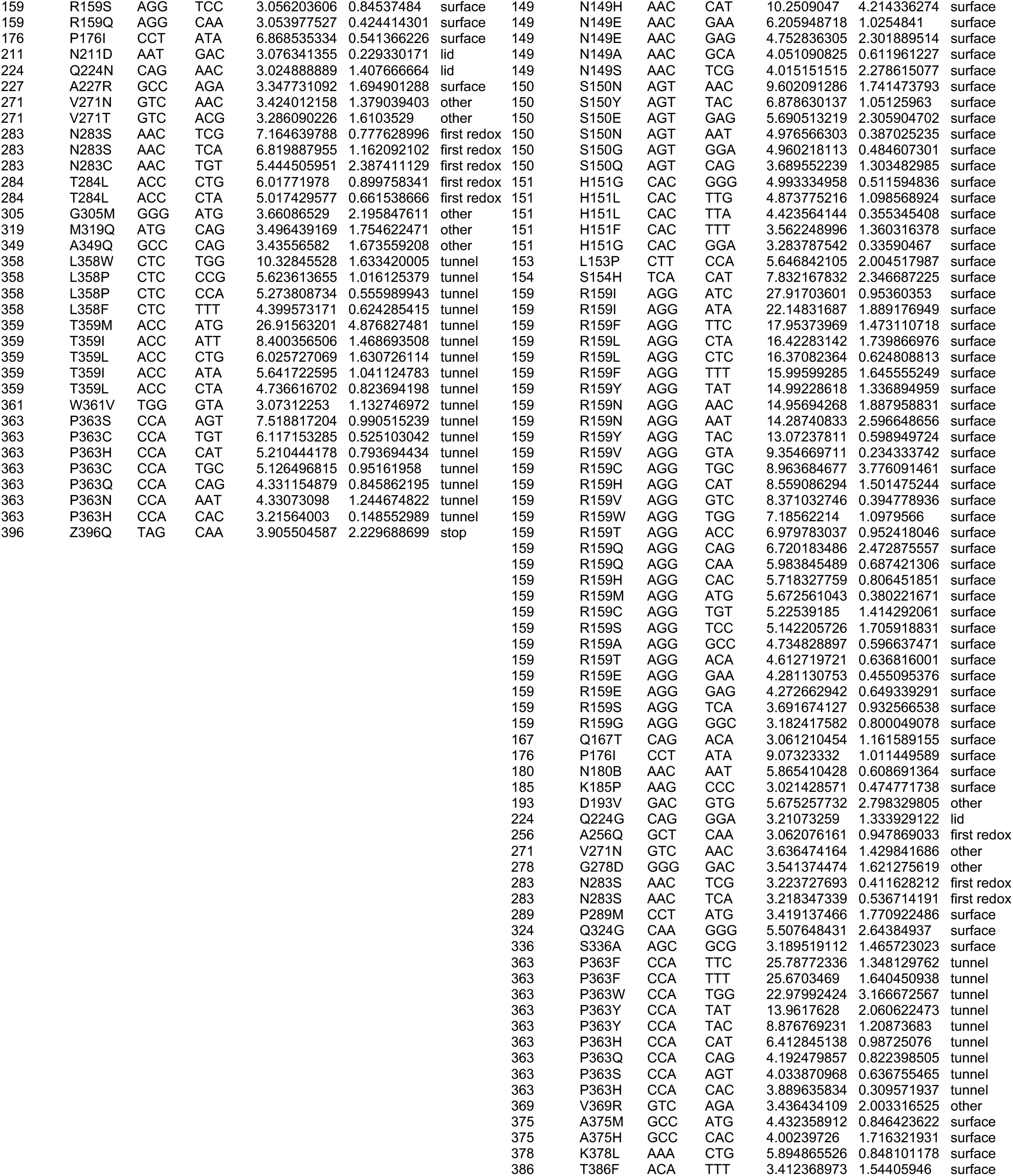
List of DHODH variants enriched ≥ 3-fold in the BAY or BRQ DMS screens. . Variants are arranged by amino acid position; wild-type and variant codons, mean fold-change (mean FC), standard error of the mean (SEM), and location of each substitution in the 3-D structure of DHODH are indicated (MLS, mitochondrial localization signal; TM-α1, putative transmembrane α-helix).

**Extended Data Table 4:** List of substitutions at each residue in DHODH that are enriched ≥ 3-fold in the BAY or BRQ DMS screens. Variants are arranged by amino acid position; wild-type codon, total number of drug-resistant substitutions, and alphabetical list of drug-resistant substitutions are indicated.

| Position | WT codon | # of BAY-resistant substitutions | List of BAY-resistant substitutions | # of BRQ-resistant substitutions | List of BRQ-resistant substitutions |
| --- | --- | --- | --- | --- | --- |
| 9 | R9 | N/A | N/A | 1 | T |
| 17 | L17 | N/A | N/A | 1 | A |
| 32 | G32 | 1 | C | 1 | C |
| 35 | R35 | 1 | G | 1 | H |
| 37 | Y37 | 2 | V, W | 1 | V |
| 38 | A38 | N/A | N/A | 2 | F, T |
| 39 | E39 | 3 | C, F, Y | 1 | Y |
| 40 | H40 | N/A | N/A | 1 | A |
| 42 | M42 | 4 | F, G, W, Y | 4 | F, G, W, Y |
| 43 | P43 | 1 | G | 1 | G |
| 44 | T44 | 1 | F | 1 | G |
| 45 | L45 | 2 | I, Y | 1 | S |
| 46 | Q46 | 9 | A, F, I, M, R, T, V, W, Y | 15 | A, C, D, E, F, G, I, L, M, N, S, T, V, W, Y |
| 51 | P51 | 1 | I | 4 | D, H, K, Y |
| 53 | S53 | 1 | R | 1 | R |
| 54 | A54 | 3 | F, I, L | N/A | N/A |
| 55 | H55 | 10 | C, E, F, I, L, P, S, T, V, Y | 12 | A, C, E, F, G, I, L, P, S, T, V, Y |
| 56 | R56 | 1 | K | 1 | I |
| 58 | A58 | 9 | C, F, H, I, L, M, T, V, Y | 8 | C, F, I, L, M, T, V, Y |
| 59 | V59 | N/A | N/A | 2 | F, W |
| 60 | R60 | 1 | Y | 2 | B, G |
| 62 | T62 | 1 | W | 2 | W, Y |
| 77 | M77 | N/A | N/A | 1 | F |
| 82 | V82 | N/A | N/A | 1 | B |
| 99 | K99 | 1 | R | N/A | N/A |
| 100 | H100 | 7 | A, I, K, L, R, T, V | 8 | A, I, K, L, R, T, V, Y |
| 112 | F112 | N/A | N/A | 1 | C |
| 122 | P122 | 2 | R, V | 2 | R, V |
| 123 | K123 | 3 | A, I, N | 2 | I, N |
| 124 | P124 | 1 | I | 1 | Q |
| 125 | Q125 | N/A | N/A | 1 | N |
| 127 | G127 | 1 | D | 1 | D |
| 128 | N128 | 1 | E | 1 | E |
| 130 | R130 | 1 | B | 3 | C, F, Q |
| 132 | R132 | 2 | H, S | 4 | A, H, S, V |
| 133 | V133 | 1 | E | 2 | M, N |
| 135 | R135 | 1 | F | 13 | A, C, E, G, K, L, N, P, Q, S, T, V, Y |
| 136 | L136 | 1 | V | 1 | V |
| 144 | N144 | 3 | A, D, S | 1 | D |
| 146 | Y146 | 8 | D, G, K, N, S, T, V, W | 7 | F, G, K, N, T, V, W |
| 148 | F148 | 6 | A, G, H, K, S, T | 7 | A, G, H, K, N, S, T |
| 149 | N149 | 4 | A, D, E, H | 5 | A, D, E, H, S |
| 150 | S150 | 8 | A, E, F, G, L, N, V, Y | 5 | E, G, N, Q, Y |
| 151 | H151 | 2 | G, L | 3 | F, G, L |
| 153 | L153 | 1 | P | 1 | P |
| 154 | S154 | 1 | H | 1 | H |
| 159 | R159 | 14 | C, E, F, H, I, L, M, N, Q, S, T, V, W, Y | 16 | A, C, E, F, G, H, I, L, M, N, Q, S, T, V, W, Y |
| 167 | Q167 | N/A | N/A | 1 | T |
| 176 | P176 | 1 | I | 1 | I |
| 180 | N180 | N/A | N/A | 1 | B |
| 185 | K185 | N/A | N/A | 1 | P |
| 193 | D193 | N/A | N/A | 1 | V |
| 211 | N211 | 1 | D | N/A | N/A |
| 224 | Q224 | 1 | N | 1 | G |
| 227 | A227 | 1 | R | N/A | N/A |
| 256 | A256 | N/A | N/A | 1 | Q |
| 271 | V271 | 2 | N, T | 1 | N |
| 278 | G278 | N/A | N/A | 1 | D |
| 283 | N283 | 2 | C, S | 1 | S |
| 284 | T284 | 1 | L | N/A | N/A |
| 289 | P289 | N/A | N/A | 1 | M |
| 305 | G305 | 1 | M | N/A | N/A |
| 319 | M319 | 1 | Q | N/A | N/A |
| 324 | Q324 | N/A | N/A | 1 | G |
| 336 | S336 | N/A | N/A | 1 | A |
| 349 | A349 | 1 | Q | N/A | N/A |
| 358 | L358 | 3 | F, P, W | N/A | N/A |
| 359 | T359 | 3 | I, L, M | N/A | N/A |
| 361 | W361 | 1 | V | N/A | N/A |
| 363 | P363 | 5 | C, H, N, Q, S | 6 | F, H, Q, S, W, Y |
| 369 | V369 | N/A | N/A | 1 | R |
| 375 | A375 | N/A | N/A | 2 | H, M |
| 378 | K378 | N/A | N/A | 1 | L |
| 386 | T386 | N/A | N/A | 1 | L |
| 396 | Z396 | 1 | Q | N/A | N/A |

**Extended Data Table 5:** Free Energy Perturbation (FEP) Analysis. Average ΔΔG values are indicated; < 0 = higher affinity; > 0 = lower affinity. Results that are discordant with experimentally-determine phenotypes are highlighted in red.

| Variant | Calculated affinity change (BAY) | BAY resistance phenotype | Calculated affinity change (BRQ) | BRQ resistance phenotype |
| --- | --- | --- | --- | --- |
| **Ubiquinone-access-tunnel/drug-binding-domain mutants** |  |  |  |  |
| Q46L | 1.51 | sensitive | 4.1 | resistant |
| Q46R | 4.38 | resistant | -1.35 | sensitive |
| A58H | 86.16 | resistant | 165.03 | resistant |
| A58T | -1.78 | resistant | 1.91 | resistant |
| R135C | 3.83 | sensitive | 15.5 | resistant |
| **Pocket 2 mutants** |  |  |  |  |
| H55Y | 3.58 | resistant | 0.51 | resistant |
| H100R | 0.93 | resistant | 3.25 | resistant |
| S150G | -0.01 | resistant | -0.07 | resistant |
| R159M | 0.2 | resistant | 0.42 | resistant |

**Extended Data Table 6:**
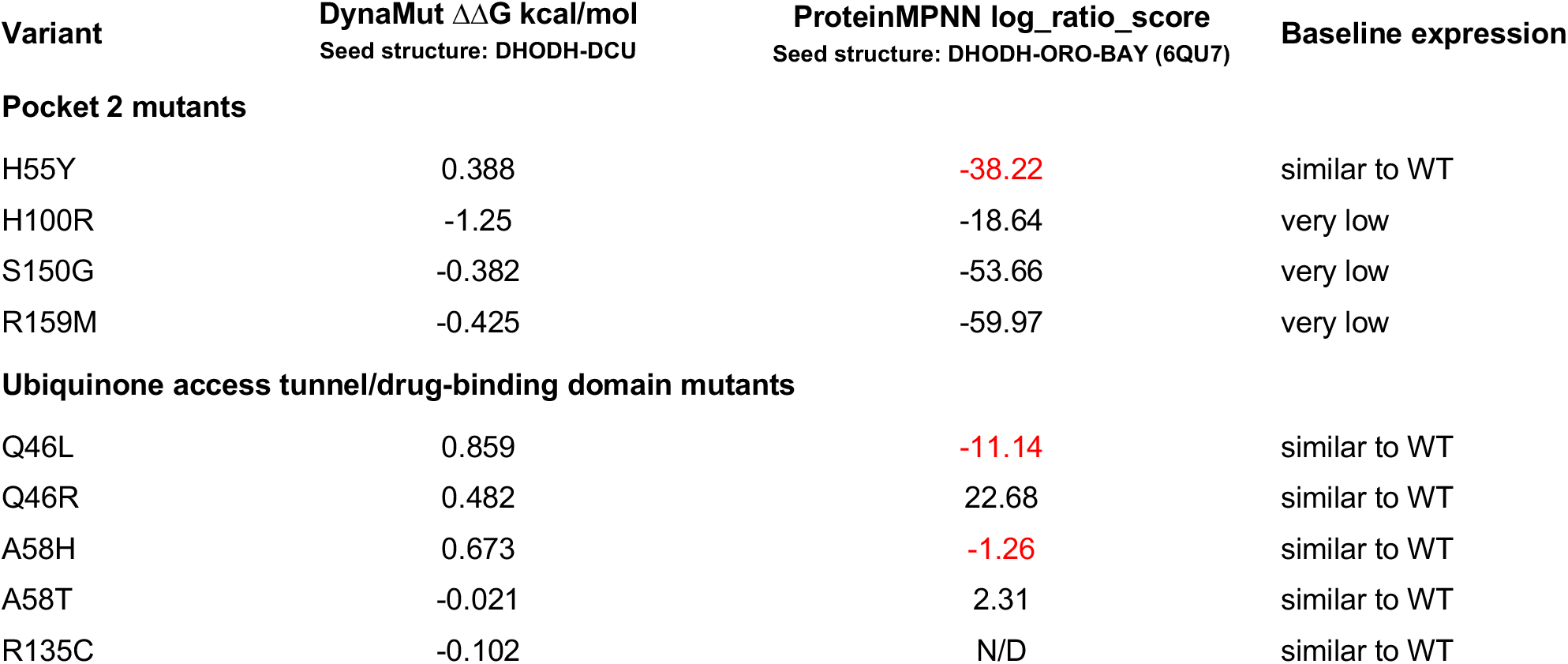
*In silico* analysis of the effects of specific substitutions on DHODH protein stability. Average ΔΔG values are indicated; < 0 = lower stability; > 0 = higher stability. Results that are discordant with experimentally-determine phenotypes are highlighted in red.

| Variant | DynaMut $\Delta\Delta G$ kcal/mol<br>Seed structure: DHODH-DCU | ProteinMPNN log_ratio_score<br>Seed structure: DHODH-ORO-BAY (6QU7) | Baseline expression |
| --- | --- | --- | --- |
| **Pocket 2 mutants** |  |  |  |
| H55Y | 0.388 | -38.22 | similar to WT |
| H100R | -1.25 | -18.64 | very low |
| S150G | -0.382 | -53.66 | very low |
| R159M | -0.425 | -59.97 | very low |
| **Ubiquinone access tunnel/drug-binding domain mutants** |  |  |  |
| Q46L | 0.859 | -11.14 | similar to WT |
| Q46R | 0.482 | 22.68 | similar to WT |
| A58H | 0.673 | -1.26 | similar to WT |
| A58T | -0.021 | 2.31 | similar to WT |
| R135C | -0.102 | N/D | similar to WT |

## Acknowledgements

We thank Jon Clardy, Stephen Blacklow, Dyann Wirth, Patrick Fischer, and Haribabu Arthanari at Harvard Medical School and Thomas Wales at Northeastern University College of Science for helpful discussions and guidance with crystallography experiments. We thank William G. Kaelin, Jr. and William Sellars at the Dana-Farber Cancer Institute for helpful discussions throughout the project. We thank Andreas Janzer, Andreas Bernthaler, Stefan Gradl, and Atanas Kamburov at Bayer for help analyzing and interpreting the DMS data. We thank members of the Losman, Dhe-Paganon, and Kaelin laboratories for technical help; Cigall Kadoch and members of her laboratory for help with protein production; Jun Qi and members of his laboratory for help with TSA development; and members of the Haigis and Vander Heiden laboratories for help developing the metabolomics assays. The metabolomics analysis was performed at the Metabolite Profiling Core Facility of the Whitehead Institute at Massachusetts Institute of Technology. Diffraction data were collected at the Center for Bio-Molecular Structure (CBMS) facility at the National Synchrotron Light Source (NSLS-II). NSLS-II is a U.S. Department of Energy (DOE) Office of Science User Facility operated under contract #DE-SC0012704. CBMS is primarily supported by the National Institutes of Health - National Institute of General Medical Sciences through a Center Core grant (P30GM133893) and by a grant from the DOE Office of Biological and Environmental Research (KP1607011). This publication resulted from data collected using beamtime obtained through NECAT BAG proposal #311097. J.-A. Losman was supported by the Dana-Farber Innovations Research Fund and Dana-Farber Cancer Institute start-up funds. M. Mravic was supported by NIH R35GM166357. M. Murakoso was supported by the Olson-King Fellowship in Computational Biology and the JASSO fellowship. S. E. Trojan acknowledges support from the Kosciuszko Foundation. The DMS screen was supported by a collaboration between the Broad Institute and Bayer.

## Author contributions

R. Pokhriyal performed all of the *in vitro* experiments and, together with J.-A. Losman, designed the experiments, analyzed the data, and assembled and wrote the manuscript. H. S. Seo and S. Dhe-Paganon performed the x-ray crystallography. M. Mravic and M. Murakoso performed the molecular dynamic simulations. S. E. Trojan, S. Joshi, M. Haigis, and M. Vander Heiden helped develop and perform the metabolomics assay. L. Evans performed the DMS screen and the initial validation experiments. J. K. O’Neil and K. Chauhan assisted with validation experiments. H. Meyer performed preliminary metabolomics analyses. A. Friberg and J. Guenther performed the FEP calculations. S. Giese performed the ProteinMPNN modeling. S. Christian assisted with initial validation experiments. X. Yang helped analyze the DMS data. R. Lintner designed and constructed the DMS library. D. E. Root leads the GPP. All of the authors helped to edit the manuscript.

## MATERIALS AND METHODS

### Cell lines and tissue culture

HEK-293T cells (ATCC, RRID:CVCL_0063) were maintained at low passage number in DMEM (Thermo Fisher #11965118) containing 10% fetal bovine serum (FBS; Thermo Fisher #A5670701) and 1% penicillin/streptomycin (P/S; Thermo Fisher #15140122) and were split every 2–3 days before reaching confluence. After being thawed, HEK-293T cells were passaged a maximum of 5 times (2 weeks) prior to use in experiments. TF-1 cells (ATCC, RRID:CVCL_0559) were maintained in RPMI (Thermo Fisher #11875119) containing 10% FBS, 1% P/S, and 2 ng/mL recombinant human GM-CSF (R&D Systems #215-GM). After being thawed, TF-1 cells were passaged a maximum of 3 times (10 days) prior to use in experiments. All cell lines were authenticated by Short Tandem Repeat profiling (ATCC) and were periodically tested for mycoplasma by Venor GeM Mycoplasma Detection Kit (Sigma Aldrich #MP0025). Most recent negative test: April 8^th^, 2026.

### Bacterial expression constructs

The pET19b vector was used to inducibly express recombinant DHODH^ΔN^ variants in bacteria. DHODH^ΔN^ variants were cloned into pET19b as follows: pET19b revcomp-SHP2_PTP, a gift from Stephen Blacklow (Harvard Medical School, Boston, MA), was digested with restriction enzymes Nde1 (NEB #R0111S) and BamH1 (NEB #R3136) to release the cDNA fragment, and digested pET19b backbone was gel-purified by QIAEX II gel extraction kit (QIAGEN #20021). The N-terminal-truncated wild-type DHODH cDNA sequence encoding amino acids 29-396 was PCR-amplified from pET22b-DHODH, a gift from Dyann Wirth (Harvard Medical School, Boston, MA), using KOD polymerase (Novagen #710863) and the following primers:

forward primer: 5’ GGAATTCCATATGGCCACGGGAGATGA 3’
reverse primer: 5’ CGCGGATCCTCACCTCCGATGATCT 3’

The PCR-amplified insert was digested with Nde1 and BamH1, gel-purified, and ligated into pET19b using T4 DNA ligase (NEB #M0202) for 16h at 16°C. The resulting plasmid harbors a 10X His-tag at the N-terminal end of the DHODH^WT-ΔN^ cDNA. The ligation mixture was transformed into XL10-Gold ultracompetent cells (Agilent #200315) and pET19b-DHODH^WT-ΔN^ clones were screened by full-plasmid sequencing (Plasmidsaurus). Point mutations to generate DHODH^Q46L-ΔN^, DHODH^Q46R-ΔN^, DHODH^A58H-ΔN^, DHODH^A58T-ΔN^, DHODH^H100R-ΔN^, DHODH^R135C-ΔN^, and DHODH^S214A-ΔN^ were introduced into pET19b-DHODH^WT-ΔN^ by QuickChange II site-directed mutagenesis (Agilent #200523) and the sequences of the selected clones were confirmed by full-plasmid sequencing.

### Mammalian expression constructs

The lentiviral expression vector pMT040 was used to express full-length DHODH variants in TF-1 cells. pMT040 was derived from pMT025 (RRID:Addgene_158579). pMT040 contains a CMV promoter-driven open reading frame (ORF) that contains a multiple cloning site followed by an IRES sequence followed by a puromycin resistance gene. The wild-type human DHODH cDNA sequence (NCBI Reference Sequence NM_001361.5) was synthesized by GenScript and was cloned into pMT040 by restriction-ligation cloning using Nhe1 (NEB #R3131) and BamH1. Point mutations to generate DHODH^H55Y^, DHODH^Q46L^, DHODH^Q46R^, DHODH^A58H^, DHODH^A58T^, DHODH^H100R^, DHODH^R135C^, DHODH^S150G^, DHODH^R159M^, DHODH^S214A^, DHODH^A58T/H100R^ and DHODH^A58TS150G^ expression vectors were introduced into pMT040-DHODH^WT^ using QuickChange II and the sequences of the selected clones were confirmed by full-plasmid sequencing. The lentiviral expression vector pLX304 was used to express DHODH variants in DHODH-knock-out TF-1 cells. Empty pLX304 and pLX304-DHODH^WT^ vectors, which contain a blasticidin-resistance cassette, were a gift from Samuel McBrayer (University of Texas at Southwestern, Dallas, Texas). To generate the CRISPR-resistant DHODH^WT^ cDNA, a silent C-to-T substitution at the PAM sequence at position 309 in DHODH^WT^ was introduced using QuickChange II. CRISPR-resistant DHODH^A58T^, DHODH^H100R^, DHODH^S150G^, DHODH^R159M^, and DHODH^S214A^ cDNAs were generated from CRISPR-resistant pLX304-DHODH^WT^ vector using QuickChange II site-directed mutagenesis. For constructs co-expressing Cas9 and a control or DHODH-targeting sgRNA, the following forward and reverse sgRNA sequences were annealed and cloned into lentiCRISPR-v2 (RRID:Addgene_52961), which contains a puromycin-resistance cassette.

sgControl forward primer: 5’ CACCGATATTTTATGACATAAAAAT 3’
sgControl reverse primer: 5’ AAACATTTTTATGTCATAAAATATC 3’
sgDHODH forward primer: 5’ CACCGCATCTTATAAAGTCCGTCCA 3’
sgDHODH reverse primer: 5’ AAACTGGACGGACTTTATAAGATGC 3’

### Expression and purification of recombinant DHODH^ΔN^ proteins

pET19b-DHODH^ΔN^ constructs were transformed in BL21 (DE3) competent E. coli (NEB #C2527I) and recombinant DHODH^ΔN^ proteins were expressed and purified according to a previously published protocol^9^, with the following modifications. To prepare the starter culture, a single colony was inoculated into 2 mL terrific broth (see stock solution list) containing 100 µg/mL ampicillin (see stock solution list) and was incubated at 37°C with shaking at 160 rpm for 16 h. A 500 µL aliquot of starter culture was used to inoculate 1 L rich media (see stock solution list) containing 100 µg/mL ampicillin. Protein expression was induced with 1 µM isopropyl β-D-thiogalactoside (IPTG; see stock solution list) when the culture reached an OD_600_ of 0.6, and the culture was then incubated at 25°C with shaking at 160 rpm for 20 h. Cells were harvested by centrifugation at 8,000 rpm for 15 min. The cell pellet from 1L culture was lysed in 100 mL ice-cold bacterial lysis buffer (see stock solution list) by slow mixing at 4°C for 30 min. The lysate was sonicated on ice for 5 min. Triton X-100 (Sigma Aldrich #T92840) was added to a final concentration of 1% and the lysate was centrifuged at 10,000 rpm for 1 h at 4°C. The supernatant was collected and maintained on ice. 1 mL TALON resin (Cytiva #28957499) was equilibrated with bacterial lysis buffer and incubated with supernatant with constant mixing at 4°C. The mixture was loaded on an Econo-Pac chromatography column (Bio-Rad #7321010) and the unbound fraction was collected as flow-through. Triton X-100 was exchanged by washing the resin with 10 mM N, N-dimethyl un-decyl-amine N-oxide (C11DAO; Cayman Chemical #20650). The resin was subsequently washed three times with 10 mL buffer containing 0, 8 or 15 mM imidazole (Sigma Aldrich #I2399). Elution was done with 160 mM imidazole in bacterial lysis buffer and the eluted fraction was buffer-exchanged into size exclusion chromatography (SEC) buffer containing 50 mM HEPES pH 7.7 (Sigma Aldrich #H3375) and 100 mM NaCl (Sigma Aldrich #S3014). Protein expression was analyzed by SDS-PAGE under reducing conditions followed by Coomassie Blue (Bio-Rad #1610786) staining. To prepare protein for crystallization, eluted fractions were subjected to size-exclusion chromatography on a Superdex 200 16/600 column (Cytiva #28989335) using SEC buffer as the running buffer, which also served as the final buffer exchange step.

### Crystallization and structure solution-refinement details

For crystallization, 200 nL protein solution was mixed with 200 nL precipitation solution. For DHODH-DCU, crystals were grown by sitting-drop vapor diffusion at 4°C using a precipitation solution of 200 µM DHODH and 600 µM DCU in 2-(N-morpholino)-ethanesulfonic acid (MES) buffer, pH 6.9 containing 17% (w/v) PEG 20000 and 1 mM tris(2-carboxyethyl)-phosphine (TCEP). For DHODH-ORO, crystals were obtained by sitting-drop vapor diffusion at 20°C using a precipitation solution of 180 µM DHODH and 2 mM orotate equilibrated against a reservoir solution of 100 mM sodium citrate, pH 6.5, 21% (w/v) PEG 3350, and 50 mM ammonium nitrate. Prior to data collection, crystals were briefly transferred into their respective precipitation solutions supplemented with 25% glycerol as a cryoprotectant and were flash-frozen in liquid nitrogen. Diffraction data were collected at beamlines 17-ID-1 (AMX)^65^ and 17-ID-2 (FMX)^66^ at the National Synchrotron Light Source II (NSLS-II; Brookhaven National Laboratory). Datasets were integrated and scaled using Xia2 package^67^. Structures were solved by molecular replacement with Phaser^68^, using the DHODH-2-hydroxypyrazolo[1,5-a]pyridine structure (PDB 6FMD) as the search model. Iterative cycles of manual model building and refinement were performed using Coot and Phenix^69,70^. Final model statistics are summarized in Extended Data Table 1.

### *In silico* structural analysis

PyMOL (Schrödinger, LLC version 3.1.6.1) was used to align molecules and prepare 3-dimensional representations. PLIP (https://plip-tool.biotec.tu-dresden.de/plip-web/plip/index) was used to identify protein-ligand interactions using default parameters. MoleOnline (https://mole.upol.cz) was used to detect tunnels in DHODH-DCU using FMN as the starting point and using default parameters. Aminode (http://www.aminode.org/search) was used to determine evolution scores. Protein Free Energy Perturbation (FEP) calculations were performed using Schrödinger’s FEP tool [Schrödinger Release 2020-3, Schrödinger, LLC, New York, NY] and applying default parameters for all selected mutations in PDB 6QU7 (for BAY) and PDB 1D3G (for BRQ) after structure preparation in Schrödinger’s Maestro tool. Calculated ΔΔG values were derived as arithmetic means over three independent runs. DynaMut (https://biosig.lab.uq.edu.au/dynamut/) was used to calculate the effects of single amino acid substitutions on the stability of DHODH-DCU using default parameters. For DHODH sequence alignments, the following amino acid sequences were obtained from UniProt (https://www.uniprot.org): *Homo sapiens* (Accession ID: Q02127), *Bos taurus* (Accession ID: Q5E9W3), *Rattus novergicus* (Accession ID: Q63707), *Mus musculus* (Accession ID: O35435), *Arabidopsis thaliana* (Accession ID: P32746), *Plasmodium falciparum* (Accession ID: Q08210), *Mycobacterium leprae* (Accession ID: P46727), *Escherichia coli* (Accession ID: P0A7E1), and *Drosophila melanogaster* (Accession ID: P32748). Multiple sequence alignments were generated using Clustal Omega (https://www.ebi.ac.uk/jdispatcher/msa/clustalo) and were modified using JalView (version 2.11.5.2)^71^.

### DMS library production

The DHODH DMS library, which contains all possible single amino acid substitutions in all 394 amino acid positions in human DHODH, for a total of 6688 variants, was constructed by the Genetic Perturbation Platform (GPP) at the Broad Institute of MIT and Harvard. A detailed description of the method of library construction is available at https://www.biorxiv.org/content/10.1101/2021.06.16.448102v7. Briefly, each variant was synthesized on an array, encoded into a 150-base DNA oligo (90-base tiles flanked by 30-base constant regions). Adjacent ORF tiles were synthesized on separate arrays. Oligonucleotide tile pools were amplified by emulsion PCR using primers to the 30-base flanking regions. Tile PCR products were isolated on 2% agarose gel and extracted for cloning. The entry vector pUC57 was linearized using Phusion polymerase (NEB #M0530) and tile region specific primers. The linearized vector was purified on 1% agarose gel and treated with Dpn1 (NEB #R0176). Dpn1-treated linear backbone was mixed with the relevant PCR-amplified tile and assembled by *in vitro* recombination with NEBuilder HiFi DNA Assembly Master Mix (NEB #E2621). Assembly reactions were column purified, electroporated into TG1 *E. coli* cells (Lucigen #60502), and recovered for 1 h at 37 °C in Recovery Media (Lucigen #80030). Aliquots from the transformations were used to inoculate overnight cultures of LB containing 25 μg/mL Zeocin (Thermo Fisher #R25005). Cells were harvested by centrifugation and plasmid DNA was isolated using Midiprep Plus kits (QIAGEN #12143). Plasmid pools, each corresponding to a tile, were verified by Nextera XT sequencing (Illumina). Plasmid tiles were pooled equally and the pool was subjected to restriction digest with Nhe1 and BamH1. After purification by 1% agarose gel, the excised linear fragment library was cloned into the pMT_040 expression vector that had been pre-processed with a compatible Nhe1-BamH1 restriction site pair. Ligation was carried out with a 5:1 insert-to-vector molar ratio, using T7 DNA ligase at room temperature for 2 h and was cleaned up with isopropanol precipitation. The resulting DNA pellet was used to transform Stbl4 bacterial cells (Thermo Fisher #11635018). Plasmid DNA (pDNA) was extracted from harvested colonies using Maxi Prep kits (QIAGEN #12162). The resulting pDNA library was sequenced on the Nextera XT platform to determine the distribution of variants in the library. Lentivirus was produced as outlined in http://www.broadinstitute.org/rnai/public/resources/protocols. Briefly, HEK-293T viral packaging cells were transfected using TransIT-LT1 transfection reagent (Mirus Bio #2306) with the pDNA library, a psPAX2 packaging plasmid containing gag, pol and rev genes (RRID:Addgene_12260), and a pMD2.G envelope plasmid containing VSV-G (RRID:Addgene_12259). Media was changed 6-8 h after transfection and virus was harvested 24 h thereafter. Viral titer was measured by infecting 3 x 10^3^ A549 cells (ATCC, RRID:CVCL_0023) per well in a 96-well cell culture plate with a series of dilutions of virus of known titer, for a standard curve, or a series of dilutions of library virus to find data points that lie in the linear range of the standard curve. 24 h post-infection, cells were subjected to puromycin selection for 2 days. Cell number post-selection was quantified with alamarBlue (Thermo Fisher #A50101).

### DMS library screening

The DHODH DMS screens were performed as follows: for each of the three replicates, 3.2 x 10^7^ TF-1 cells (∼5,000 cell per DHODH variant) were resuspended in 32 ml media containing 8 µg/ml polybrene (Santa Cruz #134220). Cells were split across 12-well plates at 2 ml per well and an appropriate volume of pooled high-titer lentivirus was added to each well to achieve a multiplicity of infection of 0.3. Spin infections were performed by centrifugation at 2000 rpm for 2 h at room temperature. After spin infection, cells were incubated for 4 h at 37°C and were then combined into a single large culture and fresh media was added. Puromycin was added 24 h after infection and cells were selected and expanded for 10 days. On day 10 post-infection, cells were counted and three aliquots of 3.2 x 10^7^ cells were plated at a concentration of 3.2 X 10^5^ cells/mL. Cells were passaged for 8 days in media containing DMSO (Thermo Fisher #D12345), 50 nM BAY2402234 (Selleck #S8847), or 5 µM brequinar (Sigma Aldrich #SML0113). Every two days, cells were counted and 3.2 x 10^7^ cells per condition were plated in fresh media with DMSO or drug. On day 0 and day 8 of screening, aliquots of 3.2 x 10^7^ cells were collected and stored at -80°C. Following completion of the screens, genomic DNA (gDNA) was isolated using a gDNA midi prep kit (QIAGEN #51183) and 75 µg gDNA per sample was submitted to the GPP for sequencing. The abundance of each variant in each replicate was determined as follows: DHODH cDNAs were PCR-amplified using the following primers:

forward primer: 5’-CCAAAATGTCGTAACAACTCCGC-3’
reverse primer: 5’-CGATCGCAGATCCTTTTAATTAATGTCTGC-3’

The volume of each PCR reaction was 50 µL and contained an optimized amount of gDNA. Q5 DNA polymerase (NEB #M0491) was used. All the PCR reactions for each gDNA sample were pooled and purified by gel electrophoresis. Bands of expected size were excised and DNA was purified, first by QIAquick (QIAGEN #28104) and then by AMPure XP (Beckman Coulter #A63881). Next-generation sequencing (NGS) samples were prepared according to the Nextera XT protocol. Each Nextera reaction used 1 ng purified PCR product. Each reaction was indexed with a unique i7/i5 index pair. After a limited-cycle PCR step, Nextera XT reactions were purified by AMPure XP and subjected to NGS on a NovaSeq S4 cell (Illumina). Sequence data processing and variant calling were performed using the variant detection software AnalyzeSaturationMutagenesis version 1.0 (ASMv1.0), as described in https://www.biorxiv.org/content/10.1101/2021.06.16.448102v7.

### Generation of stable TF-1 cell lines

Lentiviral supernatants were generated by Lipofectamine 2000 (Life Technologies #11668019) co-transfection of HEK-293T cells with cDNA-expressing or Cas9- and sgRNA-co-expressing lentiviral vectors, along with lentiviral packaging constructs psPAX2 (RRID:Addgene_14887) and pMD2.G (RRID:Addgene_14888) in a 2:2:1 ratio. To perform lentiviral infections, 2 x 10^6^ TF-1 cells were plated in media with lentiviral supernatant and 8 µg/mL polybrene, and the cells were centrifuged at 2000 rpm for 2 hours at room temperature. To generate TF-1 cells expressing DHODH variants, parental TF-1 cells were infected with a given pMT040 vector; 24 h post-infection, 1 µg/mL puromycin (Thermo Fisher #BP2956100) was added; cells were selected for 3 days; and DHODH expression was assessed by immunoblot analysis. To generate DHODH-knock-out TF-1 cells expressing sgRNA-resistant DHODH variants, parental TF-1 cells were infected with a given pLX304 vector; 24 h post-infection, 10 µg/mL blasticidin (Thermo Fisher #NC9016621) was added; cells were selected for 10 days; cells were transferred to media containing 50 µM uridine (see stock solution list); cells were infected with a given lentiCrispr-v2-Puro vector; 24 h post-infection, puromycin was added; cells were selected for 3 days in antibiotic- and uridine-containing media; and DHODH expression and sgRNA-mediated DHODH knock-out were assessed by immunoblot analysis.

### Immunoblot analysis

Protein lysates were resolved on midi-protein gels (Thermo Fisher #WXP42026BOX) and transferred to 0.2 µm PVDF membranes (Thermo Fisher IPVH00010). Membranes were blocked in TBST (see stock solution list) with 5% non-fat milk, probed with primary antibodies, and detected with horseradish-peroxidase (HRP)-conjugated secondary antibodies. Primary antibodies: rabbit polyclonal anti-DHODH (Proteintech, RRID:AB_2091723); mouse monoclonal anti-vinculin (Sigma Aldrich, RRID:AB_477629). Secondary antibodies: anti-rabbit (Jackson ImmunoResearch, RRID:AB_2339149); anti-mouse (Cell Signaling Technology, RRID:AB_330924).

#### DHODH activity assays

* DCIP assay: The *in vitro* catalytic activity of DHODH^ΔN^ variants was measured by quantifying the reduction of 2,6-dichlorophenolindophenol (DCIP), as previously described^22^, with the following modifications. Purified recombinant DHODH^ΔN^ enzyme, 100 µM DCU (see stock solution list), and either DMSO or inhibitor were added to ice-cold DHODH activity buffer (see stock solution list) and incubated on ice for 45 min. 0.5 mM DCIP (Sigma Aldrich #D1878) was added to the reaction mixture and samples were incubated for 10 min at room temperature. 3-5 100 µL aliquots of each reaction mixture were transferred to 96-well plates (Greiner Bio-one #655101), reactions were initiated by addition of DHO (see stock solution list) to a final concentration of 500 µM, and the change in absorbance at 600 nM was monitored for 15 min using a FLUOstar Omega microplate reader (BMG Labtech #4153954).
* OFA assay: The quinone-dependent *in vitro* catalytic activity of DHODH^ΔN^ variants was measured by quantifying orotate production, as previously described^41^, with the following modifications. Purified recombinant DHODH^ΔN^ enzyme was incubated with DCU, and/or fumarate (Sigma Aldrich #47910), and/or vehicle or inhibitor in 100 µL DHODH activity buffer on ice for 45 min. DHO was added to initiate the reaction and the reaction solutions were incubated for 1 h at 37°C to allow for orotate production. A 50 µL aliquot of water was added to stop the reaction. To generate a fluorogenic product, 250 µL each of 4 mM 4-(Trifluoromethoxy)-benzamidoxime (4-TFMBAO; Sigma Aldrich #422231), 8 mM potassium hexacyanoferrate (III) (K_3_[Fe(CN)_6_]; Sigma Aldrich #244023) and 80 mM potassium carbonate (K_2_CO_3_; Sigma Aldrich #209619) were added to the reaction mixture and the mixture was heated at 80°C for 4 min. Samples were cooled on ice for 2 min and 3-5 100 µL aliquots of each reaction were transferred to a 96-well plate (Thermo Fisher #M33089). Fluorescence was measured at 320 nM excitation and 430 nM emission using a Molecular Devices microplate reader (SpectraMaxM5).

#### DHODH binding assays

* Thermal shift assay (TSA): TSA was performed according to the manufacturer’s protocol (Applied Biosystems #4462263), with the following modifications. A fresh 8X dilution of protein thermal shift dye was prepared for each experiment. In a pre-chilled tube, a 20 µL reaction mixture was prepared using 5.0 µL protein thermal shift buffer, 3 µL protein, 9.5 µL DMSO or inhibitor in SEC buffer, and 2.5 µL diluted protein thermal shift dye. Reaction components were mixed and transferred to a 384-well plate (Applied Biosystems #4309849), sealed with optical adhesive film (Applied Biosystems #4360954), and incubated on ice for 45 min. Plates were centrifuged at 1000 rpm for 2 min and TSA was performed on a ViiA 7 System (Thermo Fisher).

* Cellular thermal shift assay (CETSA): For each experiment, 2.2 X 10^7^ TF-1 cells stably expressing a given DHODH variant were collected by centrifugation at 300g for 3 min and washed with 10 mL ice-cold Dulbecco’s Phosphate-Buffered Saline (DPBS; Thermo Fisher #14190250). Samples were lysed by resuspension in 2.2 mL ice-cold CETSA buffer (see stock solution list) and were immediately flash-frozen in liquid nitrogen and thawed at 25°C. The freeze-thaw cycles were repeated three times. From this step onwards, samples were maintained on ice. Cell lysates were centrifuged at 17,000g for 30 min at 4°C and supernatants were collected without disturbing the pellets. Equal volumes of supernatant were divided among three pre-chilled microfuge tubes and were treated with DMSO, 50 nM BAY, or 50 nM BRQ for 45 min on ice. Following incubation, each sample was split into 100 µL aliquots in pre-chilled PCR tubes. Samples were heated for 3 min at 55°C, 58°C, 61°C, 64°C or 67°C, incubated at room temperature for 3 min, and put on ice. Samples were transferred to pre-chilled microfuge tubes and centrifuged at 17,000g for 30 min at 4°C. Supernatants were collected into pre-chilled microfuge tubes and immunoblot samples were prepared by mixing 20 µL supernatant with 8 µL CETSA loading dye (see stock solution list). Samples were heated for 10 min at 70°C and DHODH expression was assessed by immunoblot analysis.

### Cell proliferation assays

On day 0 of each experiment, 2 X 10^5^ cells/mL were plated in three 6-well plates and either DMSO, BAY, or BRQ was added to a plate. Cells were counted and split every 2-3 days. For uridine withdrawal assays, on day 0 of uridine withdrawal, cells were counted and pelleted at 500g for 5 min, washed three times with RPMI, resuspended at 2 X 10^5^ cells/mL in RPMI containing dialyzed FBS with or without 50 µM uridine, and plated in 6-well plates. Cells were counted and split every 2-3 days. Cell proliferation and viability were assessed by counting the number of viable cells/mL using a Vi-Cell Cell Viability Analyzer (Beckman Coulter) and the number of viable cells on day 6 or 9 of the experiment were plotted using GraphPad PRISM.

#### Amide-^15^N-glutamine tracing

DHODH-knock-out cells stably expressing empty vector (DHODH^Δ/EV^), full-length wild-type DHODH (DHODH^Δ/WT^), or full-length DHODH^H100R^ (DHODH^Δ/H100R^) were maintained in media supplemented with 50 µM uridine. 24 h prior to glutamine-tracing, each of the stable cell lines was transferred to isotope-tracing media (see stock solution list). On day 0 of the experiment, 1x10^6^ cells per cell line were washed once with glutamine-free RPMI, resuspended in isotope-tracing media, and cultured for 24 h in isotope-tracing media containing DMSO or 20 mM labeled L-Glutamine (L-glutamine_amide-^15^N, 98%; Cambridge Isotope Laboratories #NLM-557-1). Cells were collected by centrifugation at 500g for 5 min at 4°C, supernatant was removed, and cells were washed in 1 mL ice-cold Optima LC/MS-grade water (Fisher Scientific #W61) with 150 mM NaCl. Cells were centrifuged, supernatant was removed, and 300 µL isotope-tracing solution (see stock solution list) was immediately added to each sample and samples were vortexed for 5 sec. Samples were vortexed at level 10 for 10 min at 4°C and centrifuged at 13,300 rpm for 10 min at 4°C. 200 µL supernatant per sample was carefully withdrawn without disturbing the pellet, transferred to pre-chilled tubes, and dried using a speed-vac at 4°C. Dried polar metabolites were resuspended in 60 µL HPLC-grade water and vortexed for 5 sec. Samples were vortexed at level 10 for 10 min at 4°C followed by centrifugation at 13,300 rpm for 10 min. Finally, 25 µL of each sample was transferred to vials for liquid chromatography-mass spectrometry (LC–MS) analysis.

#### LC-MS metabolomics analysis of amide-^15^N-glutamine tracing

Metabolite profiling was conducted on a QExactive benchtop orbitrap mass spectrometer equipped with an Ion Max source and a Heated Electrospray Ionization (HESI-II) probe, which was coupled to a Dionex UltiMate 3000 HPLC system (Thermo Fisher). 2 µl of each sample was injected into a SeQuant ZIC-pHILIC 150 × 2.1 mm analytical column equipped with a 2.1 × 20 mm guard column (EMD Millipore). Both columns had particle sizes of 5 mm. The column oven and autosampler tray were maintained at 25°C and 4°C, respectively. The mobile phase consisted of buffer A (20 mM ammonium carbonate and 0.1% ammonium hydroxide) and buffer B (acetonitrile). Chromatographic separation was carried out at a flow rate of 0.150 mL min^-^¹ using a gradient that decreased from 80% to 20% buffer B over 0–20 min, followed by a rapid increase back to 80% buffer B between 20 and 20.5 min, and then a final hold at 80% buffer B from 20.5 to 28 min. The mass spectrometer was operated in full-scan mode with polarity switching enabled. Spray voltage was set to 3.0 kV, with capillary and HESI probe temperatures maintained at 275°C and 350°C, respectively. Sheath, auxiliary, and sweep gas flows were adjusted to 40, 15, and 1 arbitrary units, respectively. Data were acquired across an m/z range of 70–1,000 at a resolution of 70,000, with an automatic gain control target of 1 × 10⁶ and a maximum injection time of 20 msec. An additional scan (m/z = 220-700) was included in negative mode only to enhance detection of nucleotides. Metabolite peak areas were called using XCalibur v.2.2 (Thermo Fisher) with 5 ppm mass tolerance and referencing retention times of chemical standards. All ion counts were normalized to ^13^C9-^15^N1-phenylalanine as an internal standard.

#### Molecular dynamics (MD) simulations

* Construction of full-length DHODH structural models: The DHODH-ORO crystal structure was used as the initial model. The unresolved loop residues (67-72) in DHODH-ORO were modeled by stitching in the conformations of the loop residues from the DHODH-DCU crystal structure. The missing region of the N-terminal TM-α1 and the α2 helix (residues 1-50) were complemented by the AF3 model (AF-Q02127-F1)^72^. The DHODH^A58T^ model was derived from this full-length WT model by fixed-backbone mutagenesis in PyMOL (Schrödinger, LLC version 3.0; https://pymol.org/ (2024)). The DHODH^H100R^ model was derived from the full-length WT model by replacing residue 100 with the residue conformation from a DHODH^H100R^ AF3 model^73^. The initial model for the simulated DHODH^H100R^ BRQ-bound state was generated by replacing residues 40–62 from the corresponding region of the BRQ-bound DHODH^WT^ crystal structure (PDB 1D3G) with the BRQ coordinates and the N-terminal TM-α1 region from AF3. The orientation and embedding of the full-length models relative to membrane were predicted with PPM 3.0 server^52^. Models with varied initial membrane orientations were generated in PyMOL by rotating the DHODH model ±15° about the z-axis or translating it +10 Å along the z-axis.
* Construction of all-atom bilayer systems: All simulation systems were constructed using the CHARMM-GUI web server^74^. Topology files and parameter files for orotate and FMN were created using Ligand Reader & Modeler^75^. Simulation systems were generated using Membrane Builder^76^. DHODH models were embedded in 100% POPE bilayer 120 Å by 120 Å and solvated with 150 mM KCl with neutralization. Simulations were performed with GROMACS 2022.5 documentation (https://manual.gromacs.org/2022/reference-manual/references.htmlrefabraham2015) and CHARMM36 force field^77,78^. Water molecules were described with the TIP3 model. Systems were energy-minimized using steepest descent algorithm (5000 steps, force tolerance 1000 kJ/mol/nm) with position restraints applied to protein backbone (4000 kJ/mol/nm²) and side-chain (2000 kJ/mol/nm²) heavy atoms. A 2 fs timestep was used throughout equilibration and production, with bonds involving hydrogen constrained using LINCS. Electrostatics were treated with Particle Mesh Ewald (PME). Van der Waals interactions were treated using a force-switching function between 1.0 and 1.2 nm, with a Coulombic short-range cutoff of 1.2 nm. A 100 ps NVT equilibration phase retained the same backbone/side-chain protein restraints (4000/2000 kJ/mol/nm², no lipid restraints), with temperature controlled via velocity-rescaling coupling of protein, lipid, and solvent groups independently to a 310.15 K bath (time constant 0.1 ps). A subsequent 15 ns NPT equilibration phase was performed with 1000 kJ/mol/nm^2^ restraints on the backbone atoms of protein (side-chain and lipid restraints removed). The Berendsen thermostat was used, with protein, lipid, and solvent independently coupled to a 310.15 K bath with time constant 1.0 ps. The Berendsen barostat was used with semi-isotropic pressure coupling (1.0 bar, time constant 5.0 ps). Restraints were then removed for 1 µs production runs with temperature maintained via velocity-rescaling coupling (310.15 K, time constant 1.0 ps) and pressure maintained using a Parrinello–Rahman barostat with semi-isotropic coupling (1.0 bar, time constant 5.0 ps). Trajectory coordinates were written every 0.2 ns for analysis. GROMACS trajectory output was converted to DCD format using CatDCD - Concatenate DCD files version 4.0 (https://www.ks.uiuc.edu/Development/MDTools/catdcd, 2009) prior to analysis. All subsequent analyses were performed in Python using ProDy^79^. Trajectories were sampled at 0.2 ns intervals for all analyses unless otherwise noted.
* Protein orientation relative to the membrane: To quantify the spatial arrangement of DHODH relative to the membrane, three angular metrics were calculated from Cα coordinates for each frame: (i) a TM-α1 helix angle, defined by the vector DHODH^H100R^ connecting N- and C-termini of the TM-α1 domain (residues 7 and 25); (ii) approach angle, defined by the vector connecting the centroids of the membrane-embedding helices (α2: residues 35–47; α3: residues 52–63) to the center of β10 (residues 251–254) and α11 (residues 262–273), and (iii) rotation angle, defined by the vector connecting the centroids of α12 (residues 339–346) and α13 (residues 310–315) to the center of α6 (residues 154– 161) and β7 (residues 177–179). For each metric, the angle between the structural vector and the membrane-normal reference direction was calculated as the arccosine of the dot product of unit vectors and reported as the complement of this value (90° − θ) so that 0° corresponds to a vector lying within the membrane plane; values exceeding 90° were reflected (180° − θ) to constrain results to a physically meaningful range. Trajectories used for this analysis were pre-aligned to a common membrane reference frame prior to angle calculation.
* Membrane insertion depth (ΔZ): The vertical position of the TM-α1 helix and adjacent amphipathic helices (α2, α3) relative to the membrane surface was assessed by calculating, for each frame, the z-coordinate difference (ΔZ) between the Cα centroid of six backbone segments (N- and C-terminal boundaries of the TM-α1 helix, α2, and α3) and the center of the upper leaflet phosphate plane. The upper-leaflet reference was defined using lipid phosphate (P) atoms more than 9 Å from the protein and with z > −10 Å, to exclude lipids displaced by protein insertion.
* Root-mean-square fluctuation (RMSF): Per-residue RMSF was calculated for each replicate after superposition of the full trajectory ensemble onto protein Cα atoms of the first frame. The mean ± standard deviation across the three replicates at each residue position was plotted as a line plot.
* Substrate-tunnel width: The substrate tunnel width was measured by calculating the minimum heavy-atom distance between the Cβ atom of residue 58 (in the α3-helix) and residues 42 and 46 (in the α2-helix). Values from the simulated trajectories were compared against corresponding distances measured in reference crystal structures representing apo, ORO-bound, and DCU/BRQ-bound states.
* Substrate-tunnel volume: Volume of the substrate tunnel was calculated using POVME 3.0^80^. 10 frames were randomly selected from each cluster with distinct tunnel widths. Each extracted frame was first superposed onto a reference structure, and membrane lipids overlapping the region of interest were removed by spatial filtering prior to pocket volume calculation. Tunnel volume was then computed on a 1.0 Å spaced grid within a fixed inclusion box (17 × 20 × 27 Å, centered on the tunnel region), using an automatically identified, spatially contiguous pocket seeded near the tunnel center and a point-inclusion distance cutoff of 1.09 Å with convex-hull exclusion applied.
* K99 side-chain positioning: Two intermolecular distances around residue H/R100 were tracked: the minimum distance between the K99 side-chain amine (NZ and its hydrogens) and (i) orotate and (ii) residue 151.
* Lipid phosphate density mapping: Spatial density maps of membrane lipid phosphate atoms around DHODH were generated using MDAnalysis (https://doi.org/10.1093/bioinformatics/btr168) (https://doi.org/10.25080/majora-629e541a-00e)^81^. For each condition, trajectory frames were first superposed onto protein Cα atoms of a reference frame to remove protein rigid-body motion. Phosphate (P) atoms of POPE lipids were then used to compute a volumetric density map on a 1.0 Å grid.

#### Stock solution list

- **Ampicillin stock** (100 mg/mL) was prepared by dissolving ampicillin sodium salt (MP Biomedicals #0219452605) in water. The solution was filter-sterilized using a 0.22 µM syringe filter, split into 500 µL aliquots, and stored in -20°C.
- **Bacterial lysis buffer** was prepared fresh for each experiment by mixing 50 mM HEPES pH 7.7, 300 mM NaCl, 10% glycerol, and 1 tablet protease inhibitor (Roche #1873580).
- **Cell lysis buffer** was prepared fresh for each experiment. Stock EBC lysis buffer was prepared by mixing 50 mM Tris-HCL pH 8.0 (Sigma Aldrich #T3069), 120 mM NaCl, 0.5% NP-40 (Thermo Fisher #85124), and 5 mM EDTA pH 8.0 (Boston Bioproducts #BM-150). Stock EBC lysis buffer was stored at 4°C. For cell lysis, 1 tablet protease inhibitor (Roche #11836170001) and 1 tablet phosphatase inhibitor (Sigma Aldrich #04906837001) were solubilized in 1 mL EBC lysis buffer and the 10X cell lysis stock solution was split into 100 µL aliquots and stored at -20°C. Aliquots of 1X cell lysis working solution were prepared by adding 900 µL EBC lysis buffer to an aliquot of 10X stock solution.
- **CETSA buffer stock solution** (100X) was prepared by solubilizing 1 protease-inhibitor tablet in 1 mL stock EBC lysis buffer and the stock solution was split into 40 µL aliquots and stored at -20°C. The working solution was prepared by adding 960 µL EBC lysis buffer to the stock solution.
- **CETSA loading dye** was prepared by mixing 300 µL LDS sample buffer (Invitrogen #NP0007) and 36 µL β-mercapto-ethanol (Thermo Fisher #21985023).
- **DCU stock solution** (100 mM) was prepared by solubilizing DCU (Sigma Aldrich #D7911) in DMSO and was stored at -20°C
- **DHO stock solution** (1 M) was prepared by solubilizing DHO (Sigma Aldrich #D7128) in DMSO and was stored at -20°C.
- **DHODH activity buffer** was prepared by mixing 50 mM Tris-HCL pH 8.0, 150 mM KCl (Sigma Aldrich #P9541), 0.1% Triton X-100, and 2% DMSO.
- **IPTG stock solution** (100 mM) was prepared by dissolving IPTG (Roche Diagnostics #IPTG-RO) in water. The solution was filter-sterilized using a 0.22 µM syringe filter and stored at -20°C.
- **Isotope-tracing media** was prepared by mixing glutamine-free RPMI (Life Technologies #21870076), 10% dialyzed FBS (Thermo Fisher #A3382001), 1% P/S, 2 ng/mL recombinant human GM-CSF, and 50 µM uridine.
- **Isotope-tracing solution** was prepared by mixing 80% methanol in 20% HPLC-grade water containing 500 nM each ^13^C and ^15^N amino acid standards (Cambridge Isotope Lab #MSK-A2-1.2).
- **Rich media** was prepared in three steps. First, 2% bacto-tryptone (Thermo Fisher #211705), 1.5% yeast extract (Fisher Scientific #BP97272), 0.2% Na_2_HPO_4_ (Sigma Aldrich #S7907), 0.1% KH_2_PO_4_ (Sigma Aldrich #P0662), and 0.8% NaCl were mixed in water and autoclaved for 20 min. Second, 0.4 g/mL D-(+)-Glucose (Sigma Aldrich #G7528) was dissolved in water and filter-sterilized using a 0.22 µM syringe filter (Sigma Aldrich #CLS430769). Finally, rich media was prepared by adding 0.5% glucose solution to the tryptone solution.
- **TBS stock solution** (10X) was prepared by dissolving 87.75 g NaCl and 50 mL 2 M Tris-HCL pH 8.0 to a final volume of 1 L in distilled water. The 1X TBST working solution was prepared by diluting 100 mL 10X TBS and 1 mL Tween-20 (Roche Diagnostics #11332465001) in 899 mL distilled water.
- **Terrific broth** was prepared by dissolving 47.6 g Terrific Broth powder (Thermo Fisher #22711022) in 996 mL distilled water and adding 4 mL glycerol (Thermo Fisher #G331). The solution was autoclaved for 20 min.
- **Uridine stock** (100 mM) was prepared by dissolving uridine 5′-monophosphate disodium salt (Sigma Aldrich #U6375) in water. The solution was filter-sterilized using 0.22 µM syringe filter, split into 100 µL aliquots, and stored in -80°C.

## Data availability

Crystal structures were deposited into the Protein Data Bank with the following IDs: DHODH-DCU, PDB 12IG; DHODH-ORO, PDB 12IH.

